# Scalable, Robust, High-throughput Expression, Purification & Characterization of Nanobodies Enabled by 2-Stage Dynamic Control

**DOI:** 10.1101/2023.12.14.571655

**Authors:** Jennifer N. Hennigan, Romel Menacho-Melgar, Payel Sarkar, Michael D. Lynch

**Author notes:** To whom all correspondence should be addressed.

## Abstract

Nanobodies are single-domain antibody fragments that have garnered considerable use as diagnostic and therapeutic agents as well as research tools. However, obtaining pure VHHs, like many proteins, can be laborious and inconsistent. High level cytoplasmic expression in *E. coli* can be challenging due to improper folding and insoluble aggregation caused by reduction of the conserved disulfide bond. We report a systems engineering approach leveraging engineered strains of *E. coli*, in combination with a two-stage process and simplified downstream purification, enabling improved, robust soluble cytoplasmic nanobody expression, as well as rapid cell autolysis and purification. This approach relies on the dynamic control over the reduction potential of the cytoplasm, in combination with dynamic expression of chaperones and lysis enzymes needed for purification. Collectively, the engineered system results in more robust growth and protein expression, enabling efficient scalable nanobody production, and purification from high throughput microtiter plates, to routine shake flask cultures and larger instrumented bioreactors. We expect this system will expedite VHH development.

## Introduction

Nanobodies (VHHs) are derived from the variable domain of heavy-chain-only Camelidae antibodies ^1,2^. Nanobodies can bind to antigens with a single domain, and as a result, VHHs have several advantages over conventional antibodies and antibody fragments in terms of diffusivity, epitope binding, and stability ^3–5^. Specifically, VHHs (∼15kDa) have a size advantage over monoclonal antibodies (mAbs, ∼150kDa). In mouse models, this smaller size has resulted in improved VHH diffusivity and homogeneous intratumoral distribution compared to mAb treatment ^6^. Although mAbs can be fragmented to produce single chain variable fragments (scFv) of ∼30kDa, the compact size and structure of VHHs allows them to access epitopes along grooves and clefts that are inaccessible to scFvs which prefer flat epitopes ^4,7^. Additionally, VHHs have stability advantages over antibody fragments and scFvs^8^.

Due to these advantages, nanobodies have gained attention as potential therapeutic and diagnostic agents ^5,9,10^. Currently, the first nanobody therapeutic has been FDA approved, and there are 10 more in development ^10^. Additionally, nanobodies have demonstrated their versatility in several biotechnological domains, such as agriculture, protein purification, intracellular imaging, and controlled gene expression ^11–14^. As nanobodies gain importance across numerous applications, so do expression platforms that enable robust, high level expression and facile purification enabling rapid characterization ^15–17^. Such expression platforms can not only enable more rapid research and development but assist with clinical and commercial translation ^18,19^.

A range of expression hosts have been used to express VHHs including *E. coli* ^15,16,20^. *E. coli* has remained a preferred expression system due to its well-established characterization, ease of modification, cost-effective media, and rapid doubling time, enabling efficient and affordable protein expression ^15,16,21–23^. In *E. coli*, periplasmic expression of VHHs typically yields titers in the range of tens of milligrams per liter, whereas cytoplasmic expression has achieved up to ∼200 mg/L in both SHuffle and BL21(DE3) strains.^15,24^ Beyond intracellular processes, extracellular expression (via protein export) has resulted in more impressive yields with titers peaking at 400 mg/L^25^. However, this approach requires optimization of various factors including the signal peptide used for export, strain, and induction method for every protein target. Although *E. coli* is a popular host, yeast systems, specifically *S. cerevisiae*, have enabled even higher titers with a record of ∼600 mg/L when using a 10L fed-batch fermentation ^15,26^. However, yeast expression has its own challenges as there’s the potential for unwanted N-glycosylation of VHHs, which can interfere with antigen binding and elevate immunogenicity ^15,16,27^. Although these are promising results, the current expression platforms lack comprehensive validation to ensure consistent high-level expression across a diverse range of VHH sequences and production scales and can still require optimization for every protein. Given that *E. coli* doesn’t perform glycosylation, efforts to enhance its expression platforms remain of significant interest.

In *E. coli*, cytoplasmic expression of nanobodies presents a challenge as they contain at least one conserved disulfide bond. Expressing proteins with disulfide bonds can be a problem because the *E. coli* cytoplasm is reductive, where disulfide bonds can be reduced to thiols leading to improper folding and protein aggregation (Figure 1) ^24,28^. Yet, this reductive environment is crucial for optimal *E. coli* growth ^29^. Several essential host proteins rely on reduced cysteines for their activity, and certain native proteins act as redox state sensors through the presence of 1-2 disulfide bonds ^30–32^. When these proteins become oxidized, it can adversely affect the robust growth of *E. coli* ^33–35^.

**Figure 1.**
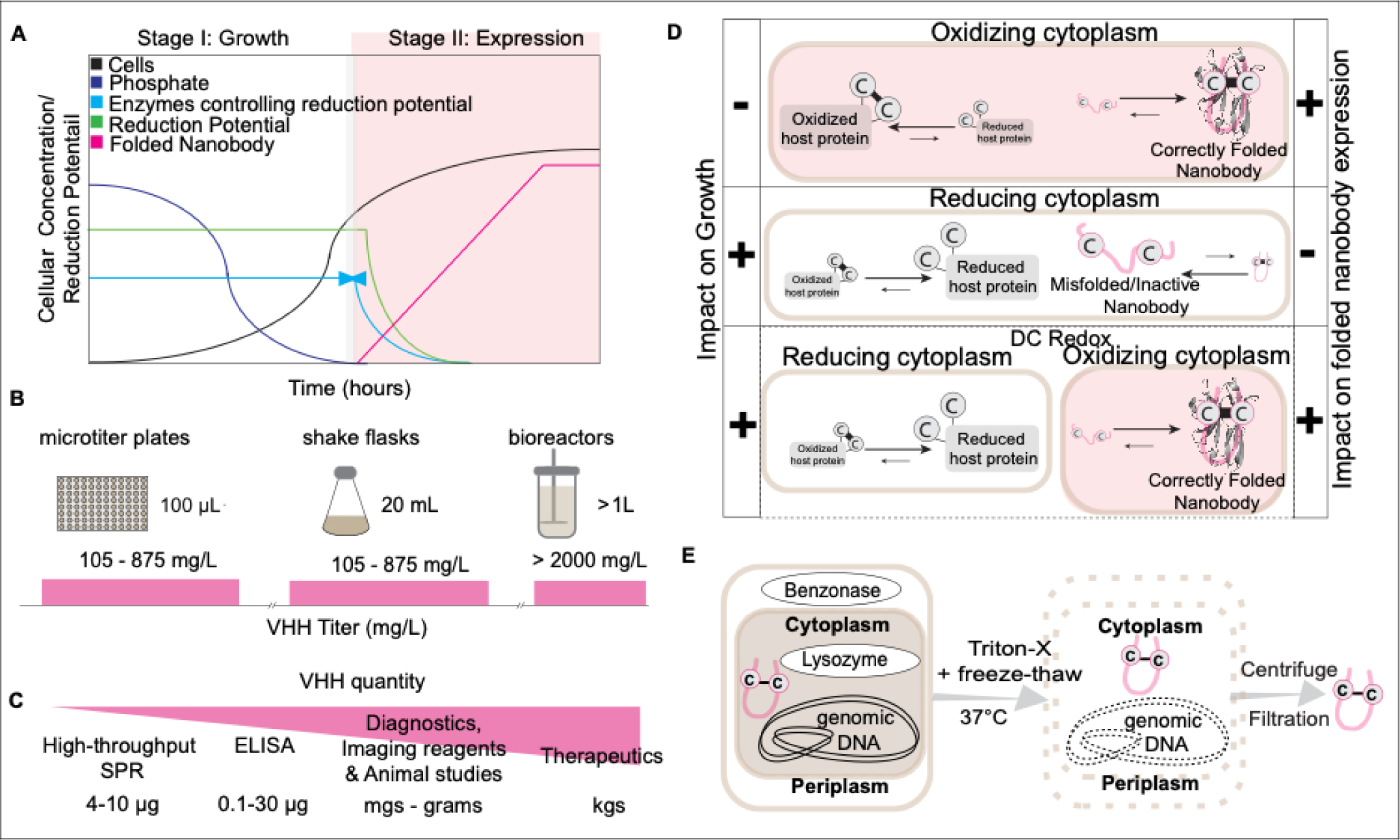
Dynamic control of cytoplasmic redox state overcomes traditional toxicity challenges and results in high titer VHH expression. A) Dynamic control strategically separates growth and production phases. Phosphate depletion in the media (dark blue) triggers VHH expression while dynamically reducing the expression of cytoplasmic reductases (light blue, valve), shifting the cytoplasm to a more oxidative state. B) Soluble VHH titer ranges at each scale with autoDC REdox. C) Ranges of VHH quantities for common evaluation studies and applications. D) Nanobodies (pink) possess a conserved disulfide bond, which is reduced in the cytoplasm (white, center). The impact of each approach on soluble VHH expression is summarized in the right table, while the effect on growth in the intracellular environment is summarized in the left table. Traditional engineering approaches aim to enhance disulfide bond formation by increasing cytoplasmic oxidation (red, top). Although this improves nanobody expression, it negatively affects growth and overall robustness. E) Autolysis integration in strain autoDC REdox streamlines VHH purification: Co-expressing lysis enzymes during phosphate-limited stationary phase, we achieve efficient cell lysis. After Triton X-100 addition, freeze-thaw cycle, and 37°C incubation, nanobodies are easily purified by filtration post-centrifugation. This innovation enhances our biomanufacturing pipeline, simplifying downstream purification for rapid and scalable applications.

Traditional engineering approaches to promote disulfide bond formation have engineered the cytoplasm to be more oxidative. Specifically, in commercially available strains such as SHuffle and Origami, cytoplasmic reductases have been deleted. These strains have compensatory mutations enabling growth ^36,37^. Alternatively, the chaperone sulfhydryl oxidase (Erv1p) can be expressed which catalyzes disulfide bond formation *de novo* ^38,39^. These prior approaches have several limitations. These approaches try to balance divergent goals of cellular growth and recombinant protein oxidation. Process optimization on a protein by protein basis is required which results in specialized case by case fermentation conditions ^36,40–42^. Additionally, existing expression hosts have primarily focused on enhancing protein oxidation, overlooking the integration of components to streamline the downstream purification process.

In this study, we took a systems engineering approach to develop an efficient nanobody biomanufacturing pipeline based on dynamic control (DC) in the context of a 2-stage fermentation ^43–46^, while additionally incorporating 2-stage expression of autolysis enzymes to streamline the purification process ^44,47^. This strategy, illustrated in Figure 1, separates the metabolic demands of recombinant protein production and cellular growth, mitigating resource competition and facilitating process development. During growth, the cytoplasm remains reductive and is dynamically converted to an oxidative environment only during production in the stationary phase, balancing growth with nanobody expression. We introduce and validate the automatic dynamic control over redox “autoDC REdox (automatic dynamic control over redox state)” strain to ensure consistent robust growth and protein expression and rapid purification, achieving consistently high titers across a library of VHH sequences. Collectively, this platform provides a robust tool for rapid VHH development and scalable production, to expedite not only VHH characterization and development, but rapid deployment in research settings as well as industrial translation.

## Results

### Strain Engineering

In this work, dynamic control leverages phosphate depletion to activate desired protein expression while reducing levels of unwanted proteins in the production phase. During the phosphate limited stationary phase, phosphate depletion initiates expression of our target protein, as well as the expression of additional proteins, such as chaperones, to enhance expression ^48^. Simultaneously, levels of unwanted proteins are reduced either using gene silencing or inducible proteolysis ^49^. Proteins targeted for proteolysis are modified with a C-terminal DAS+4 tag, which leads to ClpXP mediated degradation, in the presence of the SspB adapter protein ^49,50^. In our strains, the native *sspB* gene has been deleted and a copy placed under control of a promoter induced by phosphate depletion ^49,50^. For silencing, we utilize *E. coli*’s native Type I-E Cascade/CRISPR system. A deletion in the *cas3* gene eliminates DNA cleavage and converts the Cascade system into a silencing system when paired with gRNAs targeting the promoter of a given gene ^51,52^. Both the Cascade complex and a guide RNA (specific to the target gene) are governed by promoters induced by phosphate depletion to inhibit gene transcription in a gRNA dependent manner ^49,51–53^. In this study, we employed both methods to reduce the levels of proteins controlling the redox state of the cytoplasm, thereby enabling an oxidative state during the protein production stage of our 2-stage process. (Figure 2A).^48^

**Figure 2.**
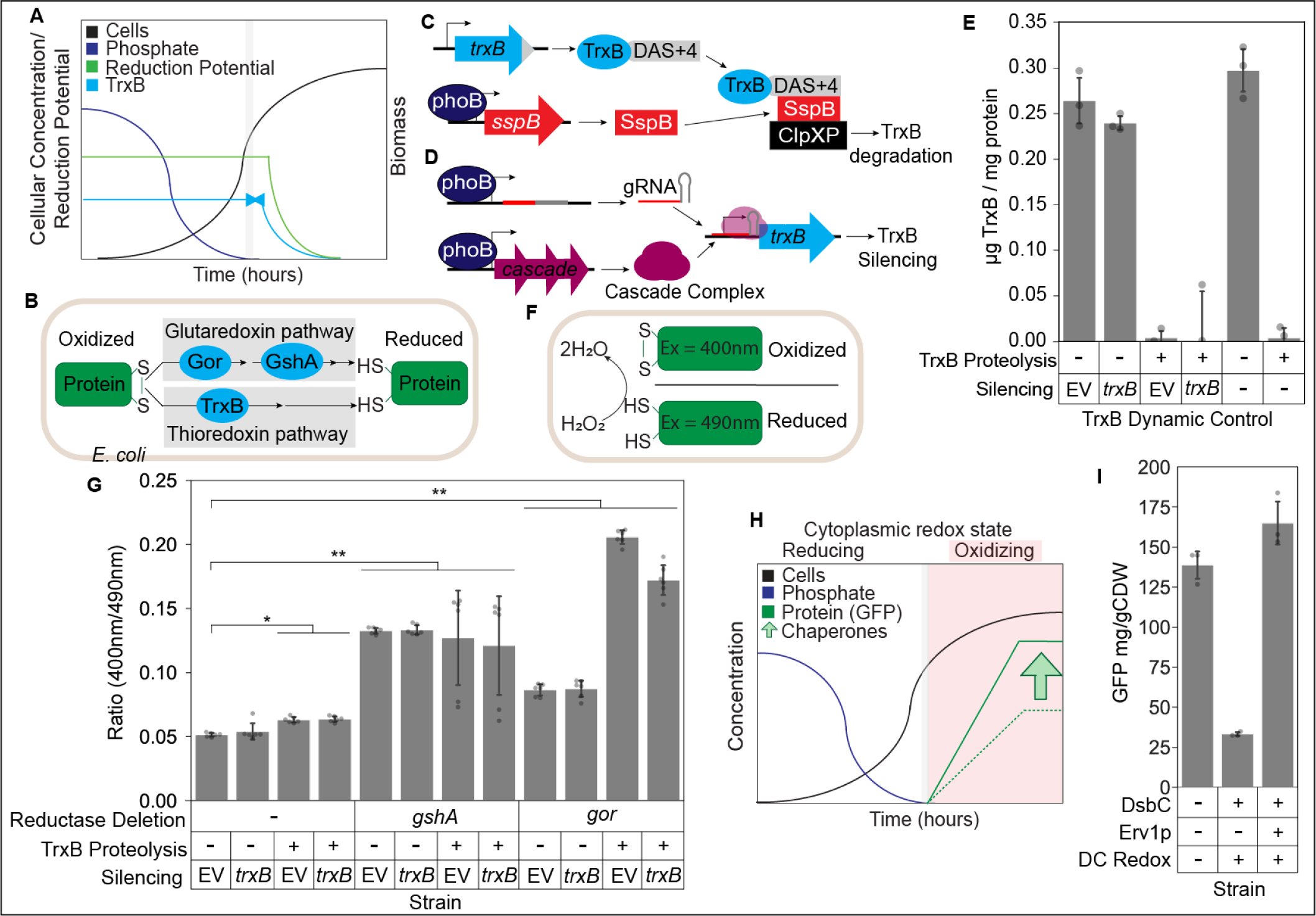
Overview of reductase modifications and the impact dynamic control on both cytoplasmic redox state and protein expression. A) The 2-stage cytoplasmic redox state is facilitated by phosphate depletion in the media (depicted in dark blue), triggered by the controlled suppression of cytoplasmic reductase TrxB (represented in light blue, valve). This regulatory mechanism induces a shift in the cytoplasm to a more oxidative state (illustrated in green). B) *E. coli* cytoplasm has two pathways (glutaredoxin and thioredoxin) to reduce disulfide bonds to thiols. Key reductases that were altered to control redox state are shown in light blue. C) Proteolysis overview: TrxB is tagged with C-terminal DAS+4 (gray), connected to ClpXP protease (black) via SspB adapter (red), under phoB promoter control. D) Silencing of *trxB* gene: Guide RNA (gRNA, red and gray) introduced on a plasmid, along with phoB promoter-controlled Cascade complex (pink), blocks *trxB* transcription during phosphate depletion in stationary phase. E) Comparison of TrxB expression levels (n = 3) in dynamic control strains and the control strain (DLF_S0025) without proteolysis or silencing. “EV” represents the empty vector plasmid, “-” indicates control without the silencing plasmid. F) Overview of redox sensitive GFP (roGFP) assay: Oxidation of roGFP alters its excitation spectrum. Increase in emission ratio (400nm/490nm) indicates greater roGFP oxidation. The bar plots represent the mean values, with error bars indicating the standard deviation. The raw data points on the bar plots correspond to the number of biological replicates. G) Evaluation of cytoplasmic oxidation using the roGFP excitation ratio in *E. coli* strains with reductase control. Dynamic control of TrxB and glutaredoxin pathway deletions led to significant excitation ratio increases (*p<0.05, **p<0.01, Welch’s t-test). H) Assessing two-stage chaperone expression impact on basal protein (GFP) in JNH_Redox5. Cytoplasmic redox state is color-coded (white for reductive, red for oxidative). Dashed line: GFP expression without chaperones; solid green line: GFP expression with chaperones (light green arrow). I) Erv1p impact on GFP expression in dynamic control strains (n=3): DLF_S0025 (control), JNH_Redox6, and JNH_Redox6E.

The parent strain used in this study, DLF_S0025, has been previously engineered to possess the DC capabilities described above (Figure 2B) ^49,51^. In this work this strain was further modified with either deletions in the glutaredoxin reductase pathway (Δ*gor* or Δ*gshA*) or dynamic control over the thioredoxin reductase pathway through either silencing and proteolysis of TrxB (Figure 2C, 2D^54^). To assess the efficacy of DC methods on TrxB, we measured protein levels in response to either gene silencing or proteolysis alone and in combination. To accomplish this, we incorporated a C-terminal superfolder GFP tag onto TrxB, either alone or upstream of the DAS+4 tag, and quantified the expression levels using a GFP ELISA (Figure 2E). Based on this data, we concluded that proteolysis alone was sufficient to significantly reduce TrxB levels.

Next we systematically evaluated strains with reduced reductase levels (*gor*, *gshA, trxB*), using the redox-sensitive GFP reporter (roGFP) to determine the impact of reductase modifications on the cytoplasmic redox state^55^. This reporter changes its excitation spectrum when oxidized due to a pair of surface cysteines forming a disulfide bond. As depicted in Figure 2F, roGFP’s excitation intensifies at 400nm upon oxidation^56^, with a rising 400nm/490nm emission ratio signaling increased cytoplasmic oxidation – a behavior validated by adding hydrogen peroxide (Figure S1). Relative to the control strain (DLF_S0025), individual modifications including TrxB proteolysis, Δ*gshA*, or Δ*gor* notably improved cytoplasmic oxidation (Figure 2G). However, combining Δ*gor* with TrxB proteolysis yielded the most pronounced oxidation. This observation was reinforced by comparing hydrogen peroxide response between the control (DLF_S0025) and the best strain from the roGFP assay (Δ*gor* + TrxB proteolysis), termed JNH_Redox5. Notably, JNH_Redox5 exhibited higher resilience to hydrogen peroxide, indicating a predominantly oxidized cytoplasm without added hydrogen peroxide. Additionally, roGFP data for JNH_Redox5 suggests a more oxidative environment than when 10mM hydrogen peroxide is introduced to the control strain.

Given the dynamic control over reducing pathways in JNH_Redox5, we compared its cytoplasmic redox state with the common commercial strain SHuffle. SHuffle lacks two reductases (Δ*gor*, Δ*trxB*) but includes a mutant peroxidase to maintain some reducing power, compensating for growth limitations ^36,37,57^. Additionally, SHuffle includes constitutive expression of the chaperone disulfide bond isomerase (DsbC). Thus, we introduced dynamic overexpression of the chaperone DsbC in strain into JNH_Redox5 to make JNH_Redox6. Biolector studies confirmed comparable roGFP ratios between JNH_Redox5, JNH_Redox6, and SHuffle (Figure S2).

Having validated the improvement in cytoplasmic redox state upon dynamic control, we wanted to evaluate if any changes to the parent strain (DLF_S0025) affected basal (non-redox sensitive) protein expression (Figure 2H). To do so, we assessed the expression of GFP (non-redox sensitive) using microfermentations in AB autoinduction media at 37°C across dynamic control strains. Notably, dynamic control over reductases (strains JNH_Redox5 and JNH_Redox6) drastically reduced basal protein expression compared to the parent strain (DLF_S0025) (Figure 2I and S3). However, considering our overarching goal of producing proteins with disulfide bonds, we introduced dynamic overexpression of the chaperone Erv1p, known for catalyzing disulfide bond formation, into strains JNH_Redox5 and JNH_Redox6, resulting in JNH_Redox5E and JNH_Redox6E. Fortunately, expression of chaperone Erv1p restored basal expression levels in both dynamic control strains to make JNH_Redox5E and JNH_Redox6E (Figure 2I and S3). To build upon this success and to further our systems engineering approach, we also integrated dynamic control of autolysis enzymes into JNH_Redox6E, culminating in the development of autoDC REdox for subsequent robustness studies ^58^. It is worth noting that we use the “yibDp gene” promoter because of its well-characterized robustness ^59^.

### Growth & expression robustness studies

We next turned to comparing the performance of autoDC REdox against the two commonly-used oxidative folding approaches, SHuffle and CyDisCo strains, depicted in Figure 3A. As previously mentioned, SHuffle strain lacks two reductases (Δ*gor*, Δ*trxB*) but includes a compensatory mutation ^36,37,57^. In contrast, CyDisCo maintains both reducing pathways but pre-expresses Erv1p and protein disulfide isomerase (PDI)^60,61^. Notably, autoDC REdox diverges from both approaches by integrating the dynamic control over reductases and dynamic overexpression of the chaperones DsbC and Erv1p. Because our approach separates the metabolic demands of cellular growth from protein expression, we hypothesized autoDC REdox would result in improved growth robustness and higher levels of protein expression.

**Figure 3.**
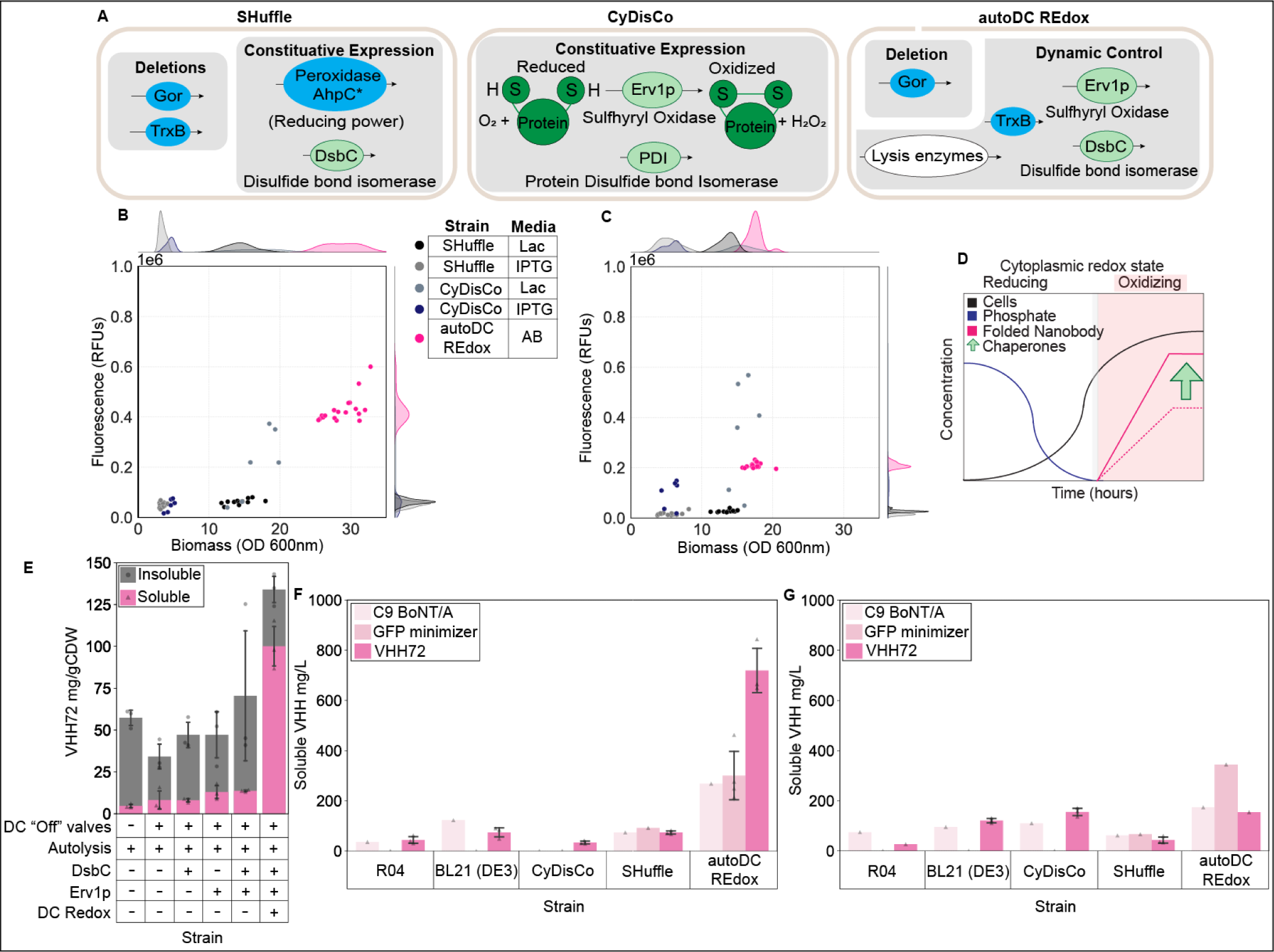
AutoDC REdox strain robustness and impact on soluble VHH expression. A) Overview of genetic modifications in each strain, highlighting enzymes with reducing power (light blue), chaperones (light green), and lysis enzymes (white). B) Growth (OD 600nm) and GFP expression robustness of SHuffle, CyDisCo, and autoDC REdox strains in microfermentations at 37 degrees. Color represents the strain-media combination: Studiers lactose autoinduction media (Lac), IPTG in LB (IPTG), phosphate autoinduction broth (AB). C) Growth and GFP expression robustness of the same strains from 30-degree microfermentations. D) Combining two-stage chaperone expression and reductase control to enhance nanobody expression in autoDC REdox. Cytoplasmic redox state is color-coded (white for reductive, red for oxidative). Dashed line: nanobody expression without chaperones; solid pink line: nanobody expression with chaperones (light green arrow). E) VHH72 shake flask expression (n=3) in dynamic control strains, stratified by soluble and insoluble fractions within cells. DC “off” valves refer to strain modifications shown in Figure 2B and Figure 2C. F-G) Soluble VHH yields obtained from shake flask expression at 37 degrees (F) and 30 degrees (G) for each strain. Color represents the expressed VHH sequence. Raw data points indicate the number of replicates, while error bars in all bar plots represent the standard deviation.

Thus, we conducted robustness studies aiming for consistent, high biomass levels and protein expression. Because lowering temperature is a common approach for oxidative folding, microfermentations were carried out at both 37°C and 30°C, probing temperature’s effect on protein expression. For these studies, AB autoinduction media was employed to test GFP expression in autoDC REdox, while SHuffle and CyDisCo strains relied on IPTG or Studier’s lactose autoinduction for their T7 induction system. The results are shown in Figures 3B and 3C, where strains closer to the top right corner indicate stronger growth and GFP expression. Under IPTG induction, both SHuffle and CyDisCo achieved an OD ∼5 at both temperatures. Notably, CyDisCo outperformed SHuffle in GFP expression, with its peak occurring at the lower temperature of 30°C. Transitioning to Studier’s lactose autoinduction media, both strains displayed increased biomass. Despite this, CyDisCo maintained better GFP expression over SHuffle, especially at 30°C. However, it’s important to note that CyDisCo’s expression lacked robustness, with replicates showing varied fluorescence. In contrast, the autoDC REdox strain grown in AB media demonstrated the most consistent growth and optimal protein expression, with minimal variation. The highest protein expression was consistently observed at 37°C, with autoDC REdox the closest to the upper right quadrant of the plot. Supplementary data from additional microfermentations is available in Figure S4.

Additionally, BioLector studies were conducted at 37°C to evaluate the growth and protein expression of SHuffle, CyDisCo, and autoDC REdox (Figure S5). SHuffle displayed limited GFP expression compared to CyDisCo, with both strains exhibiting atypical growth curves. Unlike the standard log phase, their growth slowed as GFP expression increased. In contrast, autoDC REdox exhibited consistent growth and protein expression, indicating that the integration of dynamic redox control, chaperones, and autolysis enzymes did not impede strain growth or protein expression.

### VHH solubility

To evaluate the impact of each genetic modification on nanobody expression and solubility, we utilized VHH72, a nanobody known for its binding to the spike protein of SARS-CoV-1 and SARS-CoV-2^62,63^ (Figure 3D). In this study, we incorporated autolysis enzymes into the control strain DLF_S0025, while strain R04, originally used for autolysis development^58^, but lacking the dynamically controlled “off” components, served as an additional control with intact reducing pathways. Initially, we assessed VHH72 expression levels with dynamic chaperone expression (DsbC and Erv1p) alone and in combination with dynamic redox control (TrxB proteolysis, Δ*gshA*, or Δ*gor)*. Figure 3E and Figure S6 display the expression of VHH72 in both soluble and insoluble fractions at 37°C. VHH72 is predominantly expressed in the insoluble fraction until all genetic modifications are combined in autoDC REdox strain, allowing for soluble nanobody expression. Having established the benefits of autoDC REdox strain for soluble VHH expression, we compared its performance with previously reported SHuffle and CyDisCo strains^15,24,60,64^. Considering that protein sequences can influence expression, it is essential to evaluate consistency in platform strains to enable broader applicability. We examined the soluble titers of three VHHs from the literature, including VHH72, VHH C9 BoNT/A, and VHH GFP minimizer ^65–67^. We performed expression in shake flasks using autoinduction media at both 37°C and 30°C (Figure 3F, 3G, and S7). In both sets of experiments, the autoDC REdox strain exhibited the highest titers, demonstrating its suitability as an improved platform for VHH expression across different temperatures.

### Autolysis & purification

Next we aimed to demonstrate the utility of the autoDC REdox strain and process for efficient purification of VHHs. As mentioned above, this process relies on the dynamic expression of the autolysis/autohydrolysis enzymes benzonase and lysozyme ^58^. As previously reported, we lysed cells and hydrolyzed nucleic acids using Triton X-100 and a freeze-thaw cycle, followed by a 37°C incubation step. Next, due to their small size it was possible to purify VHHs via a single filtration step (Figure 4A). We selected a 30kDa MWCO filter, close to the size of VHH (∼15kDa), to allow VHHs to pass through while retaining larger *E. coli* proteins ^68–70^. This method was validated using two nanobodies, VHH GFP enhancer and VHH GFP minimizer, known for their effects on GFP emission upon binding ^67^. Five samples of each nanobody were purified, resulting in >99% purity for VHH GFP enhancer and 81±8% purity for VHH GFP minimizer, as determined by densitometry (Figure 4B, S8). Subsequently, an in vitro binding assay with wild-type GFP confirmed the activity of the purified nanobodies (Figure 4C)^67^. The fluorescence measurements (Figure 4D) demonstrated that both VHH sequences effectively modulate GFP fluorescence, consistent with literature values^67^. These results highlight the capability of the autoDC REdox strain to facilitate VHH expression and purification. Importantly, enough active VHH is produced in shake cultures for many diagnostic applications ^71–75^(Figure 1). Overall, the autoDC REdox strain simplifies VHH expression and purification, making it a powerful tool for nanobody development.

**Figure 4.**
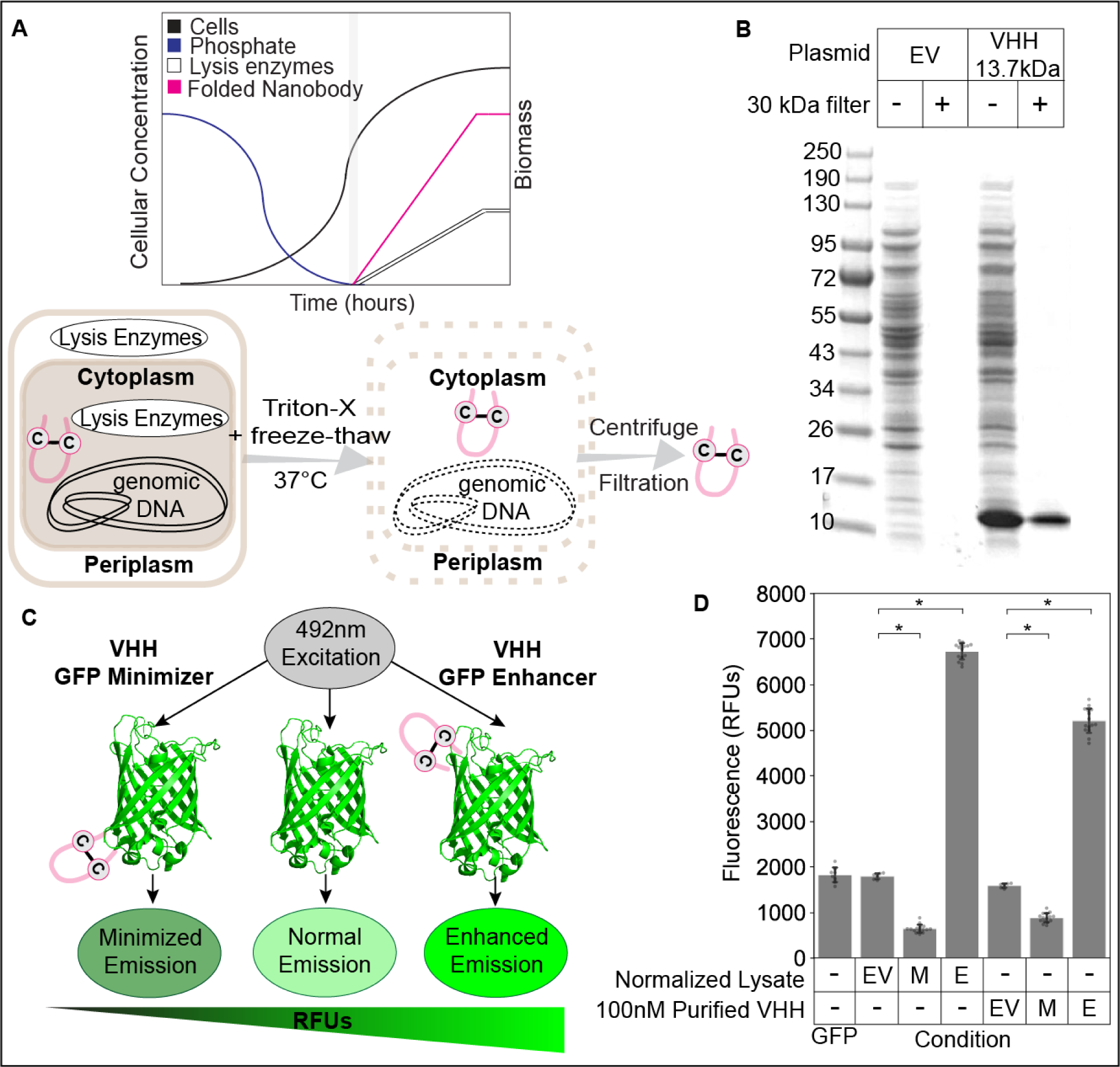
GFP VHH filtration and activity. A) Autolysis integration in strain autoDC REdox for simplified VHH purification: Lysis enzymes are co-expressed during phosphate-limited stationary phase. Following Triton X-100 addition, freeze-thaw cycle, and 37°C incubation, lysis enzymes act on their substrates. Nanobodies are purified by filtration after centrifugation. B) SDS-PAGE analysis of VHH GFP enhancer (∼13.7 kDa) filtration compared to empty vector control (EV): Filtrate shown after 30kDa filtration (+) and compared to the unfiltered soluble fraction (-). C) VHH GFP enhancer and VHH GFP minimizer activity assay overview. D) Confirmation of VHH activity through in vitro fluorescence binding assay: Three sets of conditions were evaluated to assess the activity of GFP minimizer (VHH Min) and GFP enhancer (VHH Enh). The conditions included 100nM GFP control in PBS (left), lysate addition normalized to 1 mg/mL (middle) with 100nM GFP, and 100nM purified VHH addition to 100nM GFP (right). Statistical significance (*p<0.01) was determined using Welch’s t-test, comparing the results to the EV control in the purification condition. The filtered EV samples (EV) were supplemented with the average volume of filtered VHH necessary for 100nM addition. The analysis involved 5 biological replicates, with additional raw data points representing technical replicates. Error bars represent standard deviation.

### Diversity & Scalability

Next, in assessing this platform’s versatility, we expanded our VHH library to include 15 sequences, three of which (GFP minimizer, 1B5, and JM3) had an additional disulfide bond (Figure S9) ^65–67,76–85^. We conducted shake flask expression of these VHHs in both the control strain and the autoDC REdox strain at 37°C and 30°C. Reducing the incubation temperature improved the solubility of only 25% of the library in the control strain (Figure S10, S11). In contrast, the autoDC REdox strain demonstrated 80% solubility across the VHH library at both temperatures (Figure 5A, S10), with the highest yields obtained at 37°C. All VHHs, except one, surpassed the literature benchmark for periplasmic expression, and even those with two disulfide bonds exceeded this range. Over 50% of the VHH library achieved higher expression levels than the highest reported cytoplasmic expression in *E. coli*, establishing the autoDC REdox strain as a robust platform for soluble VHH expression.

**Figure 5.**
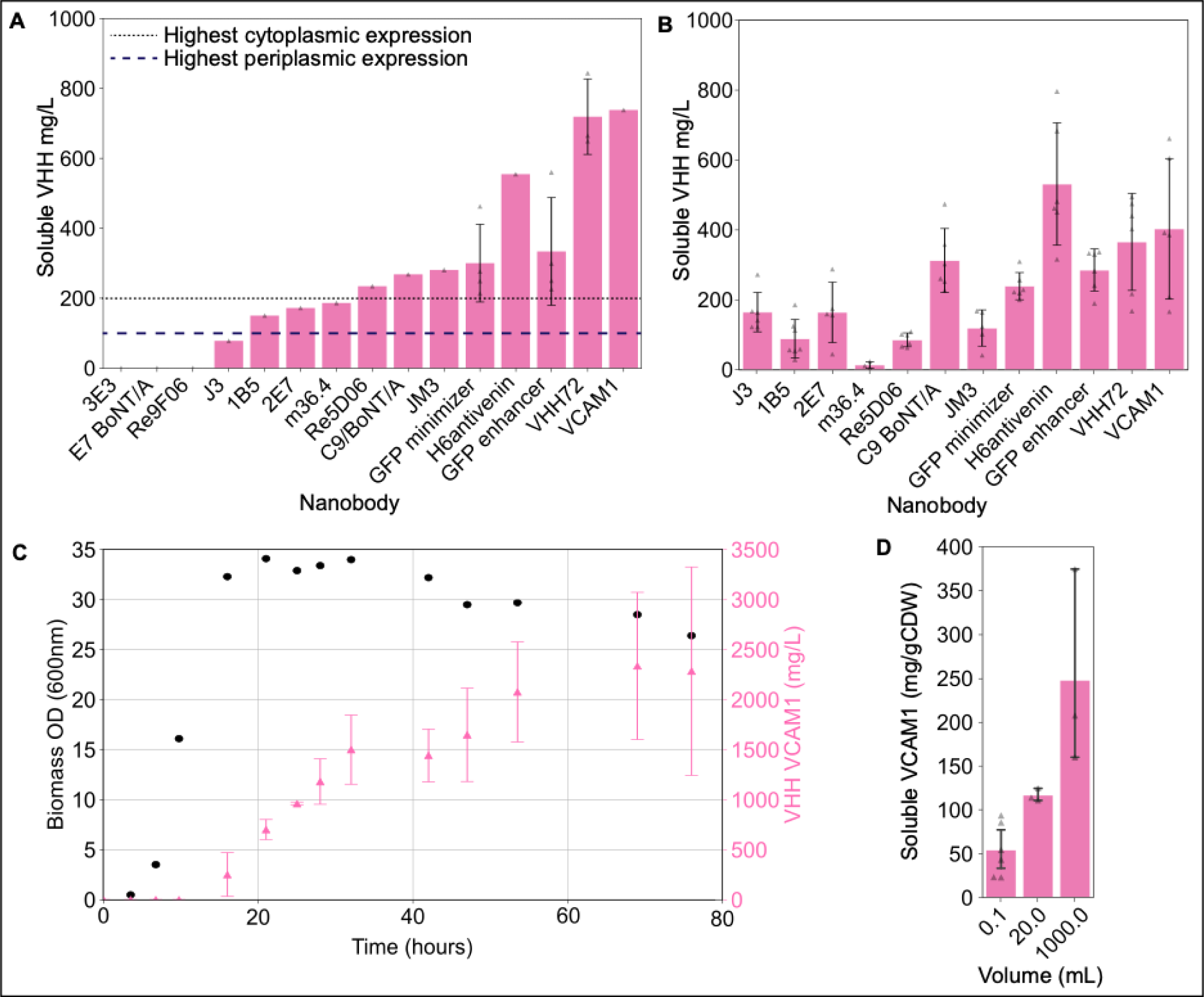
Diversity of VHHs, and Scalability of VHH production. Bar plots show the mean values and error bars represent standard deviation. A) Nanobody titers obtained from the autoDC REdox strain in shake flasks with reference to *E. coli* cytoplasmic expression in literature (gray dotted line, ∼200 mg/L) and periplasmic expression (blue dashed line, <100 mg/L)^15,24,98^. Data points represent biological replicates. B) Nanobody titers obtained from autoDC REdox expression in microfermentations at 37°C (n=3) additional data points represent technical replicates. C) Biomass (black circles) and VHH VCAM1 (pink triangles) titer achieved from a 1L bioreactor. Error bars represent standard deviation in technical replicates. D) Soluble VHH VCAM1 expression level from each scale including: microfermentations (0.1mL), shake flasks (20mL), and bioreactor (1L).

Lastly, we also evaluated the scalability of VHH production with the autoDC REdox strain, ranging from small-scale microfermentations to a 1L bioreactor ^43,86^. We confirmed high-titer expression in 96-well plates (Figure 5B and S10) by leveraging autolysis enzymes for efficient lysis, a crucial component of our engineered system that overcomes potential challenges in the absence of this mechanism ^58,87–89^. Titers matched shake flask expression (Figure S10D). Figure 5B illustrates that the majority of the library exhibited expression levels exceeding 100 mg/L, with the highest reaching an average yield of ∼500 mg/L in microfermentations, demonstrating consistent high expression levels for research and development purposes. Importantly, 100 mg/L titers in a 100 μL microtiter plate culture allows for production of up to 10 μg of protein, more than enough for higher throughput characterization via high throughout SPR shown in Figure 1 (personal communication, Carterra) ^90^. These findings validate the consistency of high VHH expression levels, in small-scale production settings crucial for research and development efforts and high-throughput screens. Furthermore, we successfully scaled up production of VHH VCAM1 in a 1 L bioreactor (Figure 5C). The VHH titer plateaued at around 70 hours, peaking at ∼2 g/L, surpassing the previously reported maximum VHH titer of ∼600 mg/L produced in *S. cerevisiae* from a 10L fed-batch fermentation (∼600 mg/L)^15,24,26^. Therefore, we concluded that the autoDC REdox strain surpasses current methods at both small-scale lab production and larger-scale biomanufacturing. Additionally, higher titers are feasible in fermentations with increased biomass level ^43^.

## Discussion

An ideal biomanufacturing process to produce recombinant nanobodies should possess several key features. Firstly, it should be user-friendly to facilitate straightforward experimental procedures. Secondly, rapid and robust growth is essential to achieve consistent and scalable high cell densities. However, as mentioned previously, in *E. coli* this goal often conflicts with strain engineering efforts to promote oxidized protein production. It is also worth noting that a strain requiring extensive optimization experiments related to temperature, aeration, or exogenous inducers is not an ideal tool for protein expression because it reduces the ease of use. Thirdly, ensuring consistency across multiple protein sequences is essential, as the host strain should enable reliable and robust expression regardless of the specific protein sequence or fermentation conditions. Additionally, an ideal process should be capable of producing high levels of oxidized protein using simple and affordable media, making it accessible for research and development purposes. Lastly, a strain that can facilitate rapid and efficient lysis, can enable high throughput screening and streamline downstream processes and protein purification.

The two-stage bioprocess utilizing strain autoDC REdox, addresses each of these desired features, distinguishing itself from all existing processes and presenting a comprehensive solution for protein expression in the context of oxidative VHH folding. Specifically, this strain combines dynamic control of cytoplasmic redox state, chaperone expression, nanobody expression, and autolysis to achieve high levels of soluble nanobodies within 24 hours of inoculation using an autoinduction approach. Furthermore, the process enables parallel sample lysis with mild conditions and minimal hands-on time. This is in contrast to current approaches for oxidative folding that rely on physical lysis (sonication or homogenization), enzymatic lysis, or chemical lysis. These methods have limitations such as the need for specialized equipment, throughput limitations, or inconsistent and inefficient lysis methods ^58,87–89^. Therefore this process provides a user-friendly and time-effective platform that allows for high protein yields with minimal effort, making it a highly efficient and convenient option.

In this study, we were successful in expressing nanobodies with the two-stage bioprocess utilizing strain autoDC REdox. Specifically, 80% of our nanobody library was expressed in the soluble fraction, with 68% surpassing routine yields for periplasmic expression (<100 mg/L)^15^. Additionally, 50% of the nanobodies exceeded the highest reported titer for cytoplasmic expression (200 mg/L)^15,24^, with the highest titer reaching >700 mg/L in a shake flask. Therefore, our results not only surpassed the highest nanobody titers reported in *E. coli* but also exceeded the highest titers reported in other microbes, including *S. cerevisiae* fed-batch fermentation (∼600 mg/L)^15,26^. Importantly, our results were achieved in shake flasks, highlighting the efficacy of our approach. In addition to the simplified upstream process for VHH expression, the integrated process simplifies the downstream purification. By utilizing a combination of autolysis and 30kDa filtration, we achieved high purities of 81±8% and >99% for two nanobodies, which meets the purity requirements (85-95%) for in vitro diagnostic applications ^91–93^. This result highlights the potential of autoDC REdox strain for manufacturing nanobodies for use in commercial diagnostics, as well as numerous research applications.

In conclusion, the results of this study positions the two-stage bioprocess utilizing strain autoDC REdox as a key enabling technology for future research and development of nanobodies and with further advances potentially other interesting proteins with disulfide bonds.

## Author contributions

JNH and MDL conceived the study and prepared the manuscript. JNH, RMM, PS and MDL performed experiments. All authors edited and revised the manuscript.

## Declaration of Competing interest

JNH, RMM and MDL have financial interests in Roke Biotechnologies, LLC. MDL has a financial interest in DMC Biotechnologies, Inc.

## Acknowledgements

This study was supported by NIH grant 3R61AI140485.

## Methods

### Reagents and Media

Unless otherwise stated, all materials and reagents were of the highest grade possible and purchased from Sigma (St. Louis, MO) or Bio Basic (Ontario, Canada). Luria Broth, lennox formulation with lower salt, was used for strain construction and plasmid propagation and is referred to as LB media in the text. Autoinduction Broth (AB), and Studier’s lac autoinduction media were prepared as previously reported ^46,58,94^. For the preparation of AB media, it is crucial to use the specific yeast extract (ThermoFisher, Cat#288620) and casamino acids (ThermoFisher Cat#223120) used in the development of the media. Working antibiotic concentrations were as follows: kanamycin (35 µg/mL), chloramphenicol (35 µg/mL), ampicillin (100 µg/mL), tetracycline (10 µg/mL), gentamicin (25 µg/mL), and puromycin (125 µg/mL). Puromycin selection was performed with LB supplemented with 50 mM potassium phosphate buffer (pH = 8.0) to maintain pH for adequate selection.

### Strains and strain construction

Three commercially available strains were used in this study. E. cloni 10G (Lucigen, Middleton WI) was used for cloning and plasmid propagation. SHuffle T7 Express and BL21(DE3) *E. coli* used in protein expression studies were both obtained from New England Biolabs (Ipswich, MA, Cat# C3029J and Cat# C2527H). The dynamic control strain DLF_S0025 (DLF_Z0025) was constructed as previously described^49^. Strain DLF_S0025 was subsequently modified for glutaredoxin pathway deletions or TrxB C-terminal DAS4+ tag modifications (with or without superfolder-GFP tags) to alter the cytoplasmic redox state. The strains constructed in this study, along with the synthetic DNA used to make these modifications are listed in Supplemental Table 2 and 3, respectively. Briefly, the linear DNA (gBlocks,IDT Coralville, IA) encoding the indicated insertion or deletion, with a 3’ antibiotic resistance marker, were flanked with homology arms targeting the genomic locus for integration with standard recombineering methodologies^95^. The recombineering plasmid pSIM5 was a kind gift from Donald Court (NCI, https://redrecombineering.ncifcrf.gov/court-lab.html). After genomic modifications were made, single colonies were selected for colony PCR followed by gel electrophoresis to confirm the change in product size before sending the PCR product for sequencing. Primers used to confirm each modification and their corresponding sequences are listed in Supplemental Table 4.

### Cloning and Plasmids

Primers and plasmids used in this study are listed in Supplemental Table 4 and Supplemental Table 1, respectively. Two cloning approaches were used to assemble plasmids: gibson assembly or a standard KLD. For gibson assembly, the fragments were assembled with NEBuilder HiFi DNA Assembly master mix (NEB, Ipswich, MA, Cat# E2621L) according to the manufacturer’s protocol. Reagents for the KLD reactions came from NEB: T4 PNK (Cat #M0236S), T4 DNA ligase (Cat#M0202S), and DpnI (Cat#R0176S) in the T4 ligase reaction buffer. All PCR steps where the product was used in a cloning reaction were performed with Q5 High-Fidelity polymerase (NEB, Ipswich, MA, Cat#M0492S). After confirming the PCR product size with gel electrophoresis the PCR product was incubated with DpnI (NEB, Ipswich, MA, Cat #R0176S) for 1 hour at 37°C to digest the plasmid template. DpnI was heat inactivated at 80°C for 20 minutes. After assembly, each reaction was diluted (1:20) and transformed into electrocompetent E. cloni. The cells were recovered for 1 hour at 37°C and plated on LB agar against the appropriate antibiotic. Single colonies were selected for colony PCR with Taq polymerase (Taq 2X Master Mix, NEB, Ipswich, MA, Cat#M0270L), then sent for sequencing for final confirmation. Primers used for colony PCR and sequencing inserts in plasmids pSMART, pCOLA, pETM6, and pCASCADE are noted in supplemental table 4. Sequence confirmed plasmids were purified (Zyppy Plasmid Miniprep Kit, Zymo Research, Irvine, CA, Cat#D4019) for further studies.

For GFPuv studies, the control plasmid pSMART-HCKan-EV (Addgene Cat#202466) was constructed from the linear pSMART-HCKan vector supplied in the Lucigen CloneSmart kit (Lucigen, Middleton WI, Cat#40708-2) with a single ligation step. All plasmids that express GFPuv under the control of a phoB promoter were obtained from Addgene (supplemental table 1) ^43,46^. Plasmid pETM6-HCAmp-T7-GFPuv (Addgene Cat#202463) was constructed with a gibson assembly of the following two fragments. The first fragment came from PCR amplification of pETM6-HCAmp-EV (Addgene Cat#49795) with primers JNH13 and JNH14. The *GFPuv* gene was amplified from pSMART-HCKan-yibDp-GFPuv (Addgene Cat#127078) with primers JNH15 and JNH16.

For roGFP expression, pSMART-HCKan-yibDp-roGFP2 (Addgene #202462) was constructed as follows. Plasmid pSMART-HCKan-yibDp-ald*-alaE (Addgene Cat#134939) was amplified with primers JNH17 and JNH18. The PCR product was mixed with the gblock encoding roGFP2 (yibD-roGFP2) listed in supplemental table 3 for gibson assembly^96^. For T7 induction pETM6-HCAmp-T7-roGFP was constructed from the following two fragments with gibson assembly. The first fragment was PCR amplified from pETM6-HCAmp-EV (Addgene Cat#49795)(P. Xu et al., 2012) with primers JNH13 and JNH19. The second fragment was amplified from pSMART-HCKan-yibDp-roGFP with primers JNH20 and JNH21 to introduce homology arms.

The silencing plasmid pCASCADE-LCAmp-EV was obtained from addgene (Addgene Cat#65821). Plasmid pCASCADE-LCAmp-trxB (Addgene Cat#202465)was constructed with a KLD reaction using pCASCADE-LCAmp-EV as a PCR template with, 5% DMSO, and primers JNH22 and JNH23. Chaperone plasmids were cloned into pCOLA vectors. First, pCOLA-Gent-ppiB (Addgene Cat#202482) was cloned from the phosphorylated gBlock listed in Supplemental Table 3 with T4 DNA ligase (Cat#M0202S) overnight at room temperature. This plasmid was used as a template to clone pCOLA-Gent-yibD-Erv1p (Addgene Cat#202483) with gibson assembly. Briefly, pCOLA-Gent-ppiB was amplified with JNH24 and JNH25. The Erv1p insert was PCR amplified from strainautoDC REdox with primers JNH26 and JNH27. Primers JNH28 and JNH29 were used to introduce the EM7 promoter with a KLD to assemble plasmid pCOLA-Gent-EM7-Erv1p (Addgene Cat #202484). Primer JNH30 was used to confirm this modification with sequencing. Plasmid pCOLA-Gent-EM7-Erv1p-DsbC (Addgene Cat#202485) and pCOLA-Gent-EM7-Erv1p-PDI were both cloned with 2 fragment gibson assemblies (Addgene Cat#202486). For each assembly, the backbone was amplified from pCOLA-Gent-EM7-Erv1p with primers JNH24 and JNH31. DsbC was amplified with primers JNH32 and JNH33 to introduce homology arms, whereas a PDI gblock was used for assembly. Lastly, the empty vector pCOLA plasmid (Addgene cat#202491) was cloned with a KLD from the PCR product resulting from amplification of pCOLA-Gent-EM7-Erv1p with primers JNH24 and JNH34.

All nanobody expression plasmids were cloned with gibson assembly. For low phosphate induction, plasmid pSMART-HCKan-yibDp-GFPuv (Addgene Cat#127078) was amplified with primers JNH35 and JNH3636. The PCR product was mixed, in individual reactions, with the VHH gblocks listed in supplemental table 3. For T7 induction, pETM6-HCAmp-EV (Addgene Cat#49795) was amplified with primers JNH13 and JNH14. Nanobodies VHH72, GFP minimizer, and C9 BoNT/A were amplified from their respective pSMART plasmids with the primers indicated in supplemental table 4 to introduce homology arms for assembly into pETM6.

### Microfermentations

Microfermentations were conducted in 96-well plates (Genesee Scientific, San Diego, CA, Cat#25-104) using a previously reported method ^43,46,86^. Glycerol stocks of each strain were used to inoculate LB overnight cultures in the 96-well plates, with 2.5μL of glycerol stock in 100μL of LB media per well. Pre-autoclaved sandwich covers (EnzyScreen, Haarlem, The Netherlands, Model #CR1596) were placed on the plates to minimize evaporative loss during incubation. The plates were cultured at 37°C, 300 rpm for 16 hours (shaker orbit of 50 mm). After 16 hours of growth, 1μL of the overnight culture was transferred to 100μL of autoinduction media (either AB or Studier’s lac, as specified) with the appropriate antibiotics. Plates were covered with new sterile sandwich covers, and the cultures were grown at 37°C, 300 rpm for 24 hours before sampling for analysis. For low-temperature microfermentations, plates of inoculated AB were grown at 30°C, 300 rpm for 48 hours.

For IPTG induction, the LB overnight starter culture was used to inoculate 100μL of fresh LB in 96-well plates. When the average plate OD reached ∼0.6 the volume used to measure OD (10μL) was replaced with LB containing concentrated IPTG to achieve a final concentration of 1mM IPTG. The plate was returned to the incubator for 16 hours before the final reading was taken.

### GFPuv expression analysis

GFPuv expression was quantified based on fluorescence and Optical density (OD_600 nm_) measurements taken with a Tecan Infinite 200 plate reader with a method previously described^43^. Biomass levels were quantified based on the conversion factor 1 OD_600 nm_ = 0.35gCDW. Total cellular protein was estimated at 500 mg/gDCW or 50% of dry cell weight^97^. Lastly GFPuv was quantified with the correlation 3.24 e 9 relative fluorescent units corresponding to 1 gram of GFPuv^43^.

### roGFP excitation ratio

The roGFP excitation ratio measurements were taken with a Tecan Infinite 200 plate reader. Measurements were performed using 200μL in black 96 well plates (Greiner Bio-One, Cat#655087). For roGFP fluorescence at 400nm, samples were excited at 412 nm (Omega Optical, Cat# 3024970) and emission was read at 530 nm (Omega Optical, Cat#3032166) using a gain of 50. For roGFP fluorescence at 490nm, samples were excited at 492 nm (Omega Optical, Cat# W2984) and emission was read at 535 nm (Omega Optical, Cat#W4803) using a gain of 50. The ratio is taken by dividing the relative fluorescence units when exciting at 490nm by the relative fluorescence units when exciting at 400nm.

### Hydrogen peroxide addition

Hydrogen peroxide addition was performed after microfermentations. The indicated hydrogen peroxide concentration was freshly prepared in solution by diluting 3% hydrogen peroxide solution and used to fill 195 μL per well in black 96 well plates. Then 5 μL of sample was added to the black 96 well plate and mixed with pipetting and left to sit for 5 minutes based on previous reports^55^. After 5 minutes the roGFP excitation ratio was measured.

### TrxB quantification

Expression levels of TrxB were quantified in the strains by incorporating a C-terminal superfolder GFP tag with standard recombineering methodologies. The resulting strains, JNH_Redox9 and JNH_Redox11, listed in Supplemental Table 2 were used to quantify TrxB expression levels with the GFP ELISA kit (Abcam, Cambridge, UK, Cat# ab171581) according to the manufacturer’s protocol. Strains were grown in triplicate in 96 well plate microfermentations. Cells were harvested by centrifugation at 3500 rpm (2900 rcf) for 10 minutes in a Thermo Sorvall Legend XTR Refrigerated Centrifuge. The pellets were resuspended in 50 μL of BugBuster Protein Extraction Reagent (Millipore Sigma, SKU 70584-4) and vortexed on Benchmark Scientific Incu-Mixer MP at room temperature, 500rpm, for 20 minutes.The lysate was clarified by a second round of centrifugation for use in the ELISA. The TrxB expression level was normalized based on total protein concentration in the sample quantified with a standard Bradford assay.

### Biolector studies

Growth measurements were obtained in a Biolector (m2p labs, 11 Baesweiler, Germany) using a high mass transfer FlowerPlate (m2p-labs, CAT#: MTP-48-B). Biolector settings were as follows: Biomass gain 25, shaking speed 800 rpm, humidity 85%. The temperature was either 37°C or 30°C as indicated. Aeration was varied with fill volume with 800μL used for high aeration and 1500μL used for low aeration. Single colonies of each strain were inoculated into 5 mL LB culture tubes with appropriate antibiotics and incubated at 37 ℃, 150 rpm overnight. The OD_600 nm_ of the overnight cultures was measured and normalized to OD_600 nm_ = 25. The normalized culture was used to inoculate individual wells of the FlowerPlate containing AB media with the appropriate antibiotics with a 1 to 100 dilution. Every strain was analyzed in triplicate. At the end of the Biolector run, the roGFP excitation ratio was measured as described above.

### Shake flask expression

Shake flask expression was performed and adapted from a method previously described^48^. Briefly, an overnight LB culture of the indicated strain and expression plasmid was used to inoculate 20 mL of AB autoinduction media with the appropriate antibiotics in a vented baffled 250mL Erlenmeyer flasks (VWR, cat. no. 89095-270). Flasks were incubated at 37°C, 150 r.p.m. (50 mm diameter) for 24 hours, and at 30°C, 150 r.p.m. (50 mm shaking orbit) for 48 hours before harvest.

### Fermentation seed culture preparation

A 5 mL LB culture, supplemented with the appropriate antibiotics, was inoculated with the specified strain from a glycerol stock and incubated at 37°C overnight. Subsequently, 1% of the overnight culture was transferred into a 250 mL baffled shake flask containing 50 mL of SM10++ media53 and incubated at 37°C overnight. When the culture’s OD6_00 nm_ reached 6-10, a 50 mL culture of SM10+ media (containing half the concentration of cas amino acids and yeast extract as SM10++) was inoculated with 1% of the SM10++ culture and incubated under the same conditions. Again when the culture’s OD_600 nm_ reached 6-10, a 50 mL culture of SM10 media (without cas amino acids or yeast extract) was inoculated similarly. Harvesting took place when the culture’s OD_600 nm_ reached 6-10. The culture was centrifuged at 4,000 rpm for 15 minutes to remove the media, and the harvested cells were resuspended in FGM10 media^43^ to achieve an OD_600 nm_ of 10. Seed vials were created by mixing 6.5 mL of the resuspended culture with 1.5 mL of sterile 50% Glycerol for long-term storage at −70°C.

### One liter fermentation

A parallel bioreactor system (Infors-HT Multifors, Laurel, MD, USA) was employed for conducting 1L fermentation experiments. The bioreactor tanks were initially filled with 800 mL of FGM10 containing an appropriate phosphate concentration to achieve a final *E. coli* biomass of approximately 10 gCDW/L. Antibiotics were added as needed. In the FGM10 medium, essential components, including phosphate, ferrous sulfate, magnesium sulfate, calcium sulfate, glucose, thiamine, and antibiotics, were introduced after sterilizing and cooling the tank vessel containing the remaining media components. Frozen seed vials containing 8 mL of seed culture were used to inoculate the tanks. Temperature and pH within the bioreactors were closely regulated at 37°C and 6.8, respectively, using 14.5 M ammonium hydroxide and 1 M hydrochloric acid as titrants. The reactor was equipped with a PID control system, which maintained control over the dissolved oxygen concentration and the glucose feeding rate was maintained at 1 g/hour 43.

### Autolysis

Autolysis was performed as previously described^58^. After shake flask expression, cells were harvested by centrifugation (4200 r.p.m., 10 minutes, 4°C) in a Thermo Sorvall Legend XTR Refrigerated Centrifuge. The pellets were resuspended in 1/10th of the original flask volume with lysis buffer (20mM Tris,pH=8.0,2mM MgCl_2_, 0.1%Triton-X100). To lyse, the cells were frozen at −80°C for at least one hour. Then the cells were thawed and incubated at 37°C for two hours. Lysates were centrifuged (16,000rpm, 20 minutes, 4°C) to isolate the soluble and insoluble fractions.

### SDS-PAGE and densitometry nanobody expression analysis

Nanobody expression levels were assessed by quantifying the final biomass using OD_600nm_ measurements. The VHH percent expression was determined through SDS-PAGE and densitometry. To quantify the total grams of cellular dry weight (gCDW), we used the final shake flask OD_600nm_, volume, and the conversion factor: 1 OD600nm = 0.35 gCDW/L (eq. 1) ^43^.

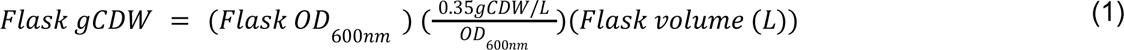

Samples were prepared to analyze the whole cell, soluble, and insoluble fractions. For whole cell analysis, a 0.4mL aliquot was saved from the shake flasks. The whole cell aliquot was resuspended in deionized water to achieve an OD_600 nm_ of 11 and mixed with an equal volume of 2x Laemmli sample buffer (Biorad, CA). The whole cell samples were boiled at 95°C for 20 minutes and then centrifuged at 16,000 rpm for 10 minutes prior to gel loading. The soluble and insoluble fractions were prepared after autolysis. Based on the OD_600nm_, shake flask volume, and autolysis resuspension volume, a 0.1gCDW aliquot was transferred to a fresh microfuge tube (eq.2).

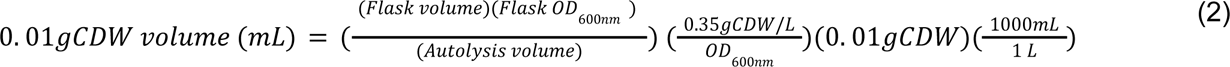

The sample was centrifuged at 16,000 rpm for 20 minutes to separate the soluble fraction from the insoluble fraction. The soluble fraction was immediately removed and protein concentration was quantified with a standard Bradford assay. Soluble samples were normalized to a concentration of 2 mg/mL and mixed with an equal volume of 2x Laemmli sample buffer. The sample was boiled at 95°C for 1 minute and then centrifuged at 16,000 rpm for 1 minute before gel loading. For gel loading, the insoluble fraction was quantified as 0.1gCDW minus the soluble protein (mg), calculated in eq.3.

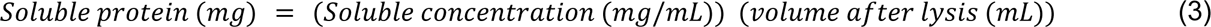

The insoluble fraction was resuspended in deionized water to a protein concentration of 2 mg/mL and mixed with 2x Laemmli sample buffer. Insoluble samples were boiled at 95°C for 20 minutes and then centrifuged at 16,000 rpm for 10 minutes before gel loading. The centrifuged samples (10 μL sample volume) were loaded into a 4-15% gradient Mini-Protean TGX precast protein gel (Biorad, CA, Cat # 4561086) and run at 140 V. The gels were stained with Coomassie Brilliant Blue R-250. Gel images were analyzed using ImageJ (NIH, MD) to determine the percent expression in each fraction.

The total flask VHH mg was quantified by multiplying the total grams of protein by the percent expression obtained from ImageJ. Total cellular protein was estimated at 500 mg/gDCW or 50% of the dry cell weight ^97^(eq. 4).

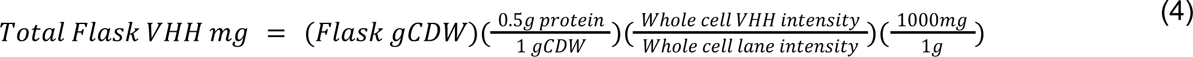

The total soluble protein content was determined through a two-step process. Firstly, the soluble VHH concentration in the autolysis resuspension was calculated with equation 5. This concentration was multiplied by the autolysis volume in equation 6. To quantify the insoluble fraction, the amount of soluble protein was subtracted from the total cellular protein using equation 7.

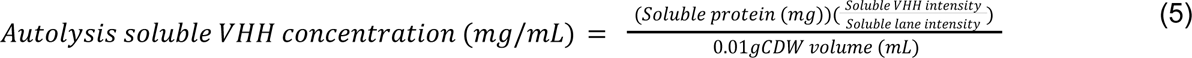

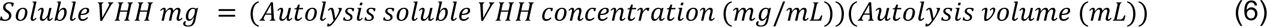

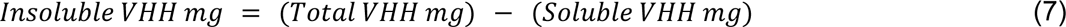

### His Tagged Nanobody quantification via ELISA

The expression levels of his-tagged nanobodies resulting from microfermentations were quantified using the His tag ELISA detection kit (GenScript, Cat# L00436) according to the manufacturer’s protocol. The same method was used to quantify VCAM1 levels after shake flask expression. Briefly, cells were harvested by removing 50μL of sample and serially diluting it (1:5) in the autolysis buffer. The diluted samples were frozen at −80°C and then incubated at 37°C for 2 hours to induce lysis. Afterward, the samples were centrifuged at 16,000rpm for 20 minutes to separate the soluble fraction from the insoluble fraction. The ELISA was performed using the soluble fraction, and incubation steps were performed at 22°C. A 5PL curve fit was used to analyze the standard curve. Samples concentrations were corrected for molar equivalency to the standard (11.3 kDa).

### Filtration based Purification

After freeze-thaw autolysis with 0.1% Triton X-100, lysates were centrifuged (16,000rpm, 20 minutes, 4°C) to isolate the soluble fraction. Filtrations were performed according to the manufacturer’s protocol, and the permeate was analyzed with SDS-PAGE. Samples were diluted to 1 mg/mL of total protein content prior to filtration to prevent filter fouling. Each membrane was pre-soaked for at least 10 minutes with autolysis buffer without Triton X-100 (20mM Tris, pH=8.0, 2mM MgCl_2_). This buffer was removed immediately before adding the sample. Clarified lysates filtered with 30kDa MWCO Amicon Ultra-0.5 Centrifugal Filter(Millipore Sigma, SKU UFC503024) were centrifuged in a table top centrifuge for 30 minutes (14,000 rcf, 4°C).

### VHH fluorimetric functional assay

VHH GFP enhancer and VHH GFP minimizer activity were validated with a fluorescence assay^67^. Fluorescence measurements were taken with a Tecan Infinite 200 plate reader in black 96 well plates (Greiner Bio-One, Cat#655087). Assay components (wtGFP, VHH, PBS buffer) were added to a final volume of 200μL per well. Pure wtGFP (Millipore Sigma, SKU 14-392) diluted in PBS to a final concentration of 100mM in the assay. Purified VHH concentrations quantified after 30kDa MWCO filtration with a Bradford assay and the slope was corrected for the percent of reactive residues relative to BSA to determine sample concentrations. Equimolar VHH GFP enhancer or VHH GFP minimizer (100mM final concentration) were added to their respective assay well and left for 5 minutes. As a control in the purification condition, the EV samples (EV) were added with the average volume of filtered VHH needed for 100nM addition. Fluorescence measurements were taken with sample excitation at 492 nm (Omega Optical, Cat# DF234) and emission was read at 530 nm (Omega Optical, Cat#3032166) using a gain of 80.

