## Supplemental Materials for "Scalable, Robust, High-throughput Expression, Purification & Characterization of Nanobodies Enabled by 2-Stage Dynamic Control"

| Supplemental Table 1: Plasmids | | | | | | |
|---|---|---|---|---|---|---|
| **Plasmid** | **Insert** | **promoter** | **Resistance** | **ori** | **Addgene** | **Source** |
| pSMART-HCKan-EV | None | None | Kanamycin | colE1 | 202466 | Lucigen (Cat # 40708-2), this study |
| pSMART-HCKan-yibDp-GFPuv | GFPuv | yibDp | Kanamycin | colE1 | 127078 | (Menacho-Melgar et al. 2020) |
| pSMART-HCKan-phoHp1-GFPuv | GFPuv | phoHp1 | Kanamycin | colE1 | 139075 | (Moreb et al. 2020) |
| pSMART-HCKan-phoAp-GFPuv | GFPuv | phoAp | Kanamycin | colE1 | 139072 | (Moreb et al. 2020) |
| pSMART-HCKan-ugpBp-GFPuv | GFPuv | ugpBp | Kanamycin | colE1 | 139078 | (Moreb et al. 2020) |
| pSMART-HCKan-phoBp-GFPuv | GFPuv | phoBp | Kanamycin | colE1 | 139073 | (Moreb et al. 2020) |
| pSMART-HCKan-amnp-GFPuv | GFPuv | amnp | Kanamycin | colE1 | 139070 | (Moreb et al. 2020) |
| pSMART-HCKan-ydfH-GFPuv | GFPuv | ydfHp | Kanamycin | colE1 | 139079 | (Moreb et al. 2020) |
| pSMART-HCKan-phoEp-GFPuv | GFPuv | phoEp | Kanamycin | colE1 | 139074 | (Moreb et al. 2020) |
| pSMART-HCKan-mipAp-GFPuv | GFPuv | mipAp | Kanamycin | colE1 | 139071 | (Moreb et al. 2020) |
| pSMART-HCKan-pstSp-GFPuv | GFPuv | pstSp | Kanamycin | colE1 | 139077 | (Moreb et al. 2020) |
| pSMART-HCKan-phoUp-GFPuv | GFPuv | phoUp | Kanamycin | colE1 | 139076 | (Moreb et al. 2020) |

| | | | | | | |
|---|---|---|---|---|---|---|
| pSMART-HCKan-yibDp-GFPuv | GFPuv | yibDp | Kanamycin | colE1 | 127078 | (Menacho-Melgar et al. 2020) |
| pSMART-HCKan-yibDp-ald*-alaE | alanine dehydrogenase and L-alanine exporter | yibDp | Kanamycin | colE1 | 134939 | (Menacho-Melgar et al. 2020) |
| pETM6-Amp-EV | None | T7 | Ampicillin | colE1 | 49795 | (Xu et al. 2012) |
| pETM6-Amp-T7-GFPuv | GFPuv | T7 | Ampicillin | colE1 | 202463 | This study |
| pSMART-HCKan-yibDp-roGFP | roGFP2 | yibDp | Kanamycin | colE1 | 202462 | This study |
| pETM6-Amp-T7-roGFP | roGFP2 | T7 | Ampicillin | colE1 | 202464 | This study |
| pCASCADE-LCAmp-EV | None | ugpB | Chloramphenicol | p15a | 65821 | pCASCADE was a gift from Michael Lynch |
| pCASCADE-LCAmp-trxB | trxB targeting gRNA | ugpB | Chloramphenicol | p15a | 202465 | This study |
| pCOLA-Gent-ppiB | ppiB | yibDp | Gentamicin | colA | 202482 | This Study |
| pCOLA-Gent-yibDp-Erv1p | Erv1p | yibDp | Gentamicin | colA | 202483 | This Study |
| pCOLA-Gent-EM7-Erv1p | Evr1p | EM7 | Gentamicin | colA | 202484 | This Study |
| pCOLA-Gent-EM7-Erv1p-DsbC | Erv1p-DsbC | EM7 | Gentamicin | colA | 202485 | This Study |
| pCOLA-Gent-EM7-Erv1p-PDI | Erv1p-PDI | EM7 | Gentamicin | colA | 202486 | This Study |
| pCOLA-Gent-EV | None | None | Gentamicin | colA | 202491 | This Study |
| pSMART-HCKan-yibDp-VHH72 | VHH72 | yibDp | Kanamycin | colE1 | 202467 | This Study |
| pSMART-HCKan-yibDp-VHH-GFPenhancer | VHH GFP enhancer | yibDp | Kanamycin | colE1 | 202468 | This Study |
| pSMART-HCKan-yibDp-VHH-GFPminimizer | VHH GFP minimizer | yibDp | Kanamycin | colE1 | 202469 | This Study |
| pSMART-HCKan-yibDp-VHH-H6antivenin | VHH H6 antivenin | yibDp | Kanamycin | colE1 | 202470 | This Study |
| pSMART-HCKan-yibDp-VHH-VCAM1 | VHH VCAM1 | yibDp | Kanamycin | colE1 | 202471 | This Study |
| pSMART-HCKan-yibDp-VHH-1B5 | VHH 1B5 | yibDp | Kanamycin | colE1 | 202472 | This Study |

| | | | | | | |
|---|---|---|---|---|---|---|
| pSMART-HCKan-yibDp-VHH-2E7 | VHH 2E7 | yibDp | Kanamycin | colE1 | 202473 | This Study |
| pSMART-HCKan-yibDp-VHH-3E3 | VHH 3E3 | yibDp | Kanamycin | colE1 | 202474 | This Study |
| pSMART-HCKan-yibDp-VHH-C9BoNT-A | VHH C9 (BoNT-A) | yibDp | Kanamycin | colE1 | 202475 | This Study |
| pSMART-HCKan-yibDp-VHH-E7BoNT-A | VHH E7 (BoNT-A) | yibDp | Kanamycin | colE1 | 202476 | This Study |
| pSMART-HCKan-yibDp-VHH-JM3 | VHH JM3 | yibDp | Kanamycin | colE1 | 202477 | This Study |
| pSMART-HCKan-yibDp-VHH-J3 | VHH J3 | yibDp | Kanamycin | colE2 | 202478 | This Study |
| pSMART-HCKan-yibDp-VHH-m36.4 | VHH m36.4 | yibDp | Kanamycin | colE1 | 202479 | This Study |
| pSMART-HCKan-yibDp-VHH-Re5D06 | VHH Re5D06 | yibDp | Kanamycin | colE1 | 202480 | This Study |
| pSMART-HCKan-yibDp-VHH-Re9F06 | VHH Re9F06 | yibDp | Kanamycin | colE1 | 202481 | This Study |
| pETM6-T7-VHH72 | VHH72 | T7 | Ampicillin | colE1 | 202487 | This Study |
| pETM6-T7-VHH-GFPminimizer | VHH GFP minimizer | T7 | Ampicillin | colE1 | 202488 | This Study |
| pETM6-T7-VHH-C9BoNT/A | VHH C9 (BoNT-A) | T7 | Ampicillin | colE1 | 202490 | This Study |

**Supplemental Table 2: Strains**

| Strain | Genotype | Source |
|---|---|---|
| E. coli 10G | F- mcrA Δ(mrr-hsdRMS-mcrBC) endA1 recA1 Φ80dlacZΔM15 ΔlacX74 araD139 Δ(ara,leu)7697galU galK rpsL nupG λ- tonA (StrR) | Lucigen (Cat # 40708-2 ) |
| SHuffle T7 Express | fhuA2 lacZ::T7 gene1 [lon] ompT ahpC gal λatt::pNEB3-r1-cDsbC (SpecR, lacIq) ΔtrxB sulA11 R(mcr-73::miniTn10--TetS)2 [dcm] R(zgb-210::Tn10 --TetS) endA1 Δgor ∆(mcrC-mrr)114::IS10 | NEB (Cat# C3029J) |
| DLF_S0025 | F-, λ-, Δ(araD-araB)567, lacZ4787(del)(::rrnB-3) , rph-1, Δ(rhaD-rhaB)568, hsdR514, ΔackA-pta, ΔpoxB, ΔpflB, ΔldhA, ΔadhE, ΔiclR, ΔarcA, ΔsspB::frt, Δcas3:: ugpBp-sspB-yibDp-casA | (Ye et al, 2020) |
| JNH_Redox | DLF_S0025, ΔgshA-purR | This study |

| | | |
|---|---|---|
| 1 | | |
| JNH_Redox 2 | DLF_S0025, Δgor-phoA-dsbC-purR | This study |
| JNH_Redox 3 | DLF_S0025, Δ:gor-purR | This study |
| JNH_Redox 4 | DLF_S0025, ΔgshA-phoA-dsbC-purR | This study |
| JNH_Redox 5 | DLF_S0025, trxB-DAS4-tetR, Δgor-purR | This study |
| JNH_Redox 6 | DLF_S0025, trxB-das4-tetR, Δgor-phoA-dsbC_purR | This study |
| JNH_Redox 7 | DLF_S0025, trxB-das4-tetR, ΔgshA-phoA-dsbC_purR | This study |
| JNH_Redox 8 | DLF_S0025, trxB-DAS4-tetR, ΔgshA-purR | This study |
| JNH_Redox 9 | DLF_S0025, trxB-sfGFP-tetR | This study |
| JNH_Redox 10 | DLF_S0025, trxB-das4-tetR | This study |
| JNH_Redox 11 | DLF_S0025, trxB-sfGFP-DAS4-tetR | This study |
| JNH_Redox 5E | DLF_S0025, trxB-das4-phoA_erv1p_tetR, Δgor-purR | This study |
| JNH_Redox 6E | DLF_S0025, trxB-das4-phoA_erv1p_tetR, Δgor-phoA-dsbC_purR | This study |
| JNH_Redox 5E_AL | DLF_S0025, trxB-das4-phoA_erv1p_tetR, Δgor-purR, Autolysis_apraR | This study |
| autoDC REdox | DLF_S0025, trxB-das4-phoA_erv1p_tetR, Δgor-phoA-dsbC_purR, Autolysis_apraR | This study |
| DLF_S0025 AL | DLF_S0025, Autolysis_apraR | This study |
| DLF_S0025 _D AL | DLF_S0025, phoA_dsbC_purR, Autolysis_apraR | This study |
| DLF_S0025 _E AL | DLF_S0025, phoA_erv1p_tetR, Autolysis_apraR | This study |
| DLF_S0025 _DE AL | DLF_S0025, phoA_dsbC_purR, phoA_erv1p_tetR, Autolysis_apraR | This study |

| | | |
|---|---|---|
| R04 | R02, Autolysis_apraR | (Menacho-Melgar et al. 2020) |

| Supplemental Table 3: G-Blocks | | | | | | |
|---|---|---|---|---|---|---|
| **G-block** | **g-block sequence** | **Res** | **Conf (F)** | **Conf (R)** | **Conf (F)** | **Conf (R)** |
| Δgor-purR | ATTACGTCTCGCGCTACAATCGCGGTAATCAACGATAAGGACACTTTGTCCAAATAAAACGAAAGGCTCAGTCGAAAGACTGGGCCTTTCGTTTTATTCCTGACGGATGGCCTTTTTGCGTTTCTACAAACTCTTTTTGTTTATTTTTCTAAATACATTCAAATATGTATCCGCTCATGAGACAATAACCCTGATAAATGCTTCAATAATATTGAAAAAGGAAGAGTATGACTGAATACAAGCCCACGGTACGCTTGGCGACGCGCGACGATGTTCCCCGCGCTGTTCGTACATTAGCTGCGGCCTTTGCAGATTACCCAGCGACGCGCCATACGGTCGATCCGGACCGCCATATCGAGCGTGTCACAGAATTGCAGGAACTTTTCTTAACTCGCGTGGGCCTTGACATCGGAAAGGTCTGGGTGGCTGACGATGGCGCTGCAGTGGCTGTTTGGACCACTCCGGAGAGTGTAGAGGCTGGTGCAGTGTTCGCCGAAATTGGTCCTCGTATGGCCGAATTAAGTGGAAGTCGTCTGGCAGCCCAACAACAAATGGAAGGGTTGCTTGCGCCCCACCGTCCGAAAGAACCCGCGTGGTTCCTTGCCACCGTTGGAGTAAGCCCAGATCACCAGGGGAAGGGTTTAGGATCTGCCGTAGTTTTACCAGGTGTGGAGGCAGCAGAACGTGCGGGAGTTCCGGCCTTCCTTGAGACGTCGGCGCCGCGCAATTTACCGTTTTACGAACGTCTTGGATTCACCGTTACGGCGGACGTGGAGGTGCCGGAGGGACCCCGTACTTGGTGTATGACTCGTAAACCGGGAGCCTGATAACATTCACCCAACGGCGGCAGAAGAGTTCGTGACAATGCGTtaaATGTTAA | Puromycin | JNH1 | JNH2 | CAAATTGAACTGGCGGTACTGC | GCCGTTAACTGATGCTCTGG |
| Δgor-phoA-dsbC-purR | ATTACGTCTCGCGCTACAATCGCGGTAATCAACGATAAGGACACTTTGTCCAAATAAAACGAAAGGCTCAGTCGAAAGACTGGGCCTTTCGTTTTATGTTAATCTTTTCAACAGCTGTCATAAAGTTGTCACGGCCGAGACTTATAGTCGCTTTGTTTTTATTTTTTAATGTATTTGTATCTAGAGATTAAAGA | Puromycin | JNH1 | JNH2 | CAAATTGAACTGGCGGTACTGC | GCCGTTAACTGATGCTCTGG |

| | |
|---|---|
| | GGAGAATACTAGatgGATGACGCGGCAATTCAA<br>CAAACGTTAGCCAAAATGGGCATCAAAAGCA<br>GCGATATTCAGCCCGCGCCTGTAGCTGGCAT<br>GAAGACAGTTCTGACTAACAGCGGCGTGTTG<br>TACATCACCGATGATGGTAAACATATCATTCAG<br>GGGCCAATGTATGACGTTAGTGGCACGGCTC<br>CGGTCAATGTCACCAATAAGATGCTGTTAAAG<br>CAGTTGAATGCGCTTGAAAAAGAGATGATCGT<br>TTATAAAGCGCCGCAGGAAAAACACGTCATCA<br>CCGTGTTTACTGATATTACCTGTGGTTACTGC<br>CACAAACTGCATGAGCAAATGGCAGACTACAA<br>CGCGCTGGGGATCACCGTGCGTTATCTTGCT<br>TTCCCGCGCCAGGGGCTGGACAGCGATGCA<br>GAGAAAGAAATGAAAGCTATCTGGTGTGCGA<br>AAGATAAAAACAAAGCGTTTGATGATGTGATG<br>GCAGGTAAAAGCGTCGCACCAGCCAGTTGCG<br>ACGTGGATATTGCCGACCATTACGCACTTGGC<br>GTCCAGCTTGGCGTTAGCGGTACTCCGGCAG<br>TTGTGCTGAGCAATGGCACACTTGTTCCGGG<br>TTACCAGCCGCCGAAAGAGATGAAAGAATTC<br>CTCGACGAACACCAAAAAATGACCAGCGGTA<br>AATAATGAccggcttatcggtcagtttcacctgatttacgtaaaa<br>acccgcttcggcgggtttttgcttttggaggggcagaaagatgaat<br>gactgtccacgacgctatacccaaaagaaaTCCTGACGG<br>ATGGCCTTTTTGCGTTTCTACAAACTCTTTTTG<br>TTTATTTTTCTAAATACATTCAAATATGTATCCG<br>CTCATGAGACAATAACCCTGATAAATGCTTCAA<br>TAATATTGAAAAAGGAAGAGTATGACTGAATAC<br>AAGCCCACGGTACGCTTGGCGACGCGCGAC<br>GATGTTCCCCGCGCTGTTCGTACATTAGCTGC<br>GGCCTTTGCAGATTACCCAGCGACGCGCCAT<br>ACGGTCGATCCGGACCGCCATATCGAGCGTG<br>TCACAGAATTGCAGGAACTTTTCTTAACTCGC<br>GTGGGCCTTGACATCGGAAAGGTCTGGGTGG<br>CTGACGATGGCGCTGCAGTGGCTGTTTGGAC<br>CACTCCGGAGAGTGTAGAGGCTGGTGCAGTG<br>TTCGCCGAAATTGGTCCTCGTATGGCCGAATT<br>AAGTGGAAGTCGTCTGGCAGCCCAACAACAA<br>ATGGAAGGGTTGCTTGCGCCCCACCGTCCGA<br>AAGAACCCGCGTGGTTCCTTGCCACCGTTGG<br>AGTAAGCCCAGATCACCAGGGGAAGGGTTTA<br>GGATCTGCCGTAGTTTTACCAGGTGTGGAGG<br>CAGCAGAACGTGCGGGAGTTCCGGCCTTCCT<br>TGAGACGTCGGCGCCGCGCAATTTACCGTTT |

| | | | | | | |
|---|---|---|---|---|---|---|
| | TACGAACGTCTTGGATTCACCGTTACGGCGG ACGTGGAGGTGCCGGAGGGACCCCGTACTT GGTGTATGACTCGTAAACCGGGAGCCTGATAA CATTCACCCAACGGCGGCAGAAGAGTTCGTG ACAATGCGTtaaATGTTAA | | | | | |
| ΔgshA-pur R | ACCATTACAGTTATGCTAATTAAAACGATTTTG ACAGGCGGGAGGTCAATCAAATAAAACGAAA GGCTCAGTCGAAAGACTGGGCCTTTCGTTTT ATTCCTGACGGATGGCCTTTTTGCGTTTCTAC AAACTCTTTTTGTTTATTTTTCTAAATACATTCA AATATGTATCCGCTCATGAGACAATAACCCTGA TAAATGCTTCAATAATATTGAAAAAGGAAGAGT ATGACTGAATACAAGCCCACGGTACGCTTGG CGACGCGCGACGATGTTCCCCGCGCTGTTCG TACATTAGCTGCGGCCTTTGCAGATTACCCAG CGACGCGCCATACGGTCGATCCGGACCGCCA TATCGAGCGTGTCACAGAATTGCAGGAACTTT TCTTAACTCGCGTGGGCCTTGACATCGGAAA GGTCTGGGTGGCTGACGATGGCGCTGCAGT GGCTGTTTGGACCACTCCGGAGAGTGTAGAG GCTGGTGCAGTGTTCGCCGAAATTGGTCCTC GTATGGCCGAATTAAGTGGAAGTCGTCTGGC AGCCCAACAACAAATGGAAGGGTTGCTTGCG CCCCACCGTCCGAAAGAACCCGCGTGGTTCC TTGCCACCGTTGGAGTAAGCCCAGATCACCA GGGGAAGGGTTTAGGATCTGCCGTAGTTTTAC CAGGTGTGGAGGCAGCAGAACGTGCGGGAG TTCCGGCCTTCCTTGAGACGTCGGCGCCGC GCAATTTACCGTTTTACGAACGTCTTGGATTC ACCGTTACGGCGGACGTGGAGGTGCCGGAG GGACCCCGTACTTGGTGTATGACTCGTAAACC GGGAGCCTGATAACCGCTGATACCGAACCGT TTGCGGTGTGGCTGGAAAAACACGCCtgaCAG | Puro mycin | JNH3 | JNH4 | GCAGT CACGC TATTAT TAAGC TGG | GTTTTC GCCACC CGAACT G |
| ΔgshA-pho A-dsbC-pu rR | ACCATTACAGTTATGCTAATTAAAACGATTTTG ACAGGCGGGAGGTCAATCAAATAAAACGAAA GGCTCAGTCGAAAGACTGGGCCTTTCGTTTT ATGTTAATCTTTTCAACAGCTGTCATAAAGTTG TCACGGCCGAGACTTATAGTCGCTTTGTTTTT ATTTTTTAATGTATTTGTATCTAGAGATTAAAGA GGAGAATACTAGatgGATGACGCGGCAATTCAA CAAACGTTAGCCAAAATGGGCATCAAAAGCA GCGATATTCAGCCCGCGCCTGTAGCTGGCAT GAAGACAGTTCTGACTAACAGCGGCGTGTTG | Puro mycin | JNH3 | JNH4 | GCAGT CACGC TATTAT TAAGC TGG | GTTTTC GCCACC CGAACT G |

| | |
|---|---|
| | TACATCACCGATGATGGTAAACATATCATTCAG GGGCCAATGTATGACGTTAGTGGCACGGCTC CGGTCAATGTCACCAATAAGATGCTGTTAAAG CAGTTGAATGCGCTTGAAAAAGAGATGATCGT TTATAAAGCGCCGCAGGAAAAACACGTCATCA CCGTGTTTACTGATATTACCTGTGGTTACTGC CACAAACTGCATGAGCAAATGGCAGACTACAA CGCGCTGGGGATCACCGTGCGTTATCTTGCT TTCCCGCGCCAGGGGCTGGACAGCGATGCA GAGAAAGAAATGAAAGCTATCTGGTGTGCGA AAGATAAAAACAAAGCGTTTGATGATGTGATG GCAGGTAAAAGCGTCGCACCAGCCAGTTGCG ACGTGGATATTGCCGACCATTACGCACTTGGC GTCCAGCTTGGCGTTAGCGGTACTCCGGCAG TTGTGCTGAGCAATGGCACACTTGTTCCGGG TTACCAGCCGCCGAAAGAGATGAAAGAATTC CTCGACGAACACCAAAAAATGACCAGCGGTA AATAATGAccggcttatcggtcagtttcacctgatttacgtaaaa acccgcttcggcgggtttttgcttttggaggggcagaaagatgaat gactgtccacgacgctatacccaaaagaaaTCCTGACGG ATGGCCTTTTTGCGTTTCTACAAACTCTTTTTG TTTATTTTTCTAAATACATTCAAATATGTATCCG CTCATGAGACAATAACCCTGATAAATGCTTCAA TAATATTGAAAAAGGAAGAGTATGACTGAATAC AAGCCCACGGTACGCTTGGCGACGCGCGAC GATGTTCCCCGCGCTGTTCGTACATTAGCTGC GGCCTTTGCAGATTACCCAGCGACGCGCCAT ACGGTCGATCCGGACCGCCATATCGAGCGTG TCACAGAATTGCAGGAACTTTTCTTAACTCGC GTGGGCCTTGACATCGGAAAGGTCTGGGTGG CTGACGATGGCGCTGCAGTGGCTGTTTGGAC CACTCCGGAGAGTGTAGAGGCTGGTGCAGTG TTCGCCGAAATTGGTCCTCGTATGGCCGAATT AAGTGGAAGTCGTCTGGCAGCCCAACAACAA ATGGAAGGGTTGCTTGCGCCCCACCGTCCGA AAGAACCCGCGTGGTTCCTTGCCACCGTTGG AGTAAGCCCAGATCACCAGGGGAAGGGTTTA GGATCTGCCGTAGTTTTACCAGGTGTGGAGG CAGCAGAACGTGCGGGAGTTCCGGCCTTCCT TGAGACGTCGGCGCCGCGCAATTTACCGTTT TACGAACGTCTTGGATTCACCGTTACGGCGG ACGTGGAGGTGCCGGAGGGACCCCGTACTT GGTGTATGACTCGTAAACCGGGAGCCTGATAA CCGCTGATACCGAACCGTTTGCGGTGTGGCT |

| | | | | | | |
|---|---|---|---|---|---|---|
| | GGAAAAACACGCCtgaCAG | | | | | |
| trxB-das4-tetR | ATGGATCACATTTATCGCCAGGCCATTACTTC<br>GGCCGGTACAGGCTGCATGGCAGCACTTGAT<br>GCGGAACGCTACCTCGATGGTTTAGCTGACG<br>CAAAAGCGGCCAACGATGAAAACTATTCTGAA<br>AACTATGCGGATGCGTCTtaaTGATCCTAATTTT<br>TGTTGACACTCTATCATTGATAGAGTTATTTTA<br>CCACTCCCTATCAGTGATAGAGAAAAGTGAAA<br>TGAATAGTTCGACAAAGATCGCATTGGTAATTA<br>CGTTACTCGATGCCATGGGGATTGGCCTTATC<br>ATGCCAGTCTTGCCAACGTTATTACGTGAATTT<br>ATTGCTTCGGAAGATATCGCTAACCACTTTGG<br>CGTATTGCTTGCACTTTATGCGTTAATGCAGG<br>TTATCTTTGCTCCTTGGCTTGGAAAAATGTCT<br>GACCGATTTGGTCGGCGCCCAGTGCTGTTGT<br>TGTCATTAATAGGCGCATCGCTGGATTACTTAT<br>TGCTGGCTTTTTCAAGTGCGCTTTGGATGCTG<br>TATTTAGGCCGTTTGCTTTCAGGGATCACAGG<br>AGCTACTGGGGCTGTCGCGGCATCGGTCATT<br>GCCGATACCACCTCAGCTTCTCAACGCGTGA<br>AGTGGTTCGGTTGGTTAGGGGCAAGTTTTGG<br>GCTTGGTTTAATAGCGGGGCCTATTATTGGTG<br>GTTTTGCAGGAGAGATTTCACCGCATAGTCCC<br>TTTTTTATCGCTGCGTTGCTAAATATTGTCACT<br>TTCCTTGTGGTTATGTTTTGGTTCCGTGAAAC<br>CAAAAATACACGTGATAATACAGATACCGAAGT<br>AGGGGTTGAGACGCAATCGAATTCGGTATACA<br>TCACTTTATTTAAAACGATGCCCATTTTGTTGA<br>TTATTTATTTTTCAGCGCAATTGATAGGCCAAA<br>TTCCCGCAACGGTGTGGGTGCTATTTACCGA<br>AAATCGTTTTGGATGGAATAGCATGATGGTTG<br>GCTTTTCATTAGCGGGTCTTGGTCTTTTACAC<br>TCAGTATTCCAAGCCTTTGTGGCAGGAAGAAT<br>AGCCACTAAATGGGGCGAAAAAACGGCAGTA<br>CTGCTCGGATTTATTGCAGATAGTAGTGCATTT<br>GCCTTTTTAGCGTTTATATCTGAAGGTTGGTTA<br>GTTTTCCCTGTTTTAATTTTATTGGCTGGTGGT<br>GGGATCGCTTTACCTGCATTACAGGGAGTGAT<br>GTCTATCCAAACAAAGAGTCATCAGCAAGGTG<br>CTTTACAGGGATTATTGGTGAGCCTTACCAAT<br>GCAACCGGTGTTATTGGCCCATTACTGTTTGC<br>TGTTATTTATAATCATTCACTACCAATTTGGGAT<br>GGCTGGATTTGGATTATTGGTTTAGCGTTTTAC | Tetracycline | JNH5 | JNH6 | GAAGTGACCGGCGATCAAATG | GCCTGGGCAATGATCAATATGC |

| | | | | | | |
|---|---|---|---|---|---|---|
| | TGTATTATTATCCTGCTATCGATGACCTTCATGT<br>TAACCCCTCAAGCTCAGGGGAGTAAACAGGA<br>GACAAGTGCTTAGTTATTTCGTCACCAAATGAT<br>TTTTACAAATCAGTAACAAAAGTAAAGAAGGC<br>GACACCATGCGACTATGG | | | | | |
| trxB-sfGFP-das4-tetR | ATGGATCACATTTATCGCCAGGCCATTACTTC<br>GGCCGGTACAGGCTGCATGGCAGCACTTGAT<br>GCGGAACGCTACCTCGATGGTTTAGCTGACG<br>CAAAAGGGGGTTCAGGCGGGTCGGGTGGCgt<br>gagcaagggcgaggagctgttcaccggggtggtgcccatcctgg<br>tcgagctggacggcgacgtaaacggccacaagttcagcgtgcg<br>cggcgagggcgagggcgatgccaccaacggcaagctgaccct<br>gaagttcatctgcaccaccggcaagctgcccgtgccctggcccac<br>cctcgtgaccaccctgacctacggcgtgcagtgcttcagccgctac<br>cccgaccacatgaagcgccacgacttcttcaagtccgccatgccc<br>gaaggctacgtccaggagcgcaccatcagcttcaaggacgacg<br>gcacctacaagacccgcgccgaggtgaagttcgagggcgacac<br>cctggtgaaccgcatcgagctgaagggcatcgacttcaaggagg<br>acggcaacatcctggggcacaagctggagtacaacttcaacag<br>ccacaacgtctatatcaccgccgacaagcagaagaacggcatc<br>aaggccaacttcaagatccgccacaacgtggaggacggcagc<br>gtgcagctcgccgaccactaccagcagaacacccccatcggcg<br>acggccccgtgctgctgcccgacaaccactacctgagcacccag<br>tccgtgctgagcaaagaccccaacgagaagcgcgatcacatgg<br>tcctgctggagttcgtgaccgccgccgggatcactcacggcatgg<br>acgagctgtacaagGGTGGGGGTGGGAGCGGCG<br>GCGGTGGCTCCGCGGCCAACGATGAAAACTA<br>TTCTGAAAACTATGCGGATGCGTCTtaatgaTCC<br>TAATTTTTGTTGACACTCTATCATTGATAGAGTT<br>ATTTTACCACTCCCTATCAGTGATAGAGAAAAG<br>TGAAATGAATAGTTCGACAAAGATCGCATTGG<br>TAATTACGTTACTCGATGCCATGGGGATTGGC<br>CTTATCATGCCAGTCTTGCCAACGTTATTACGT<br>GAATTTATTGCTTCGGAAGATATCGCTAACCAC<br>TTTGGCGTATTGCTTGCACTTTATGCGTTAATG<br>CAGGTTATCTTTGCTCCTTGGCTTGGAAAAAT<br>GTCTGACCGATTTGGTCGGCGCCCAGTGCTG<br>TTGTTGTCATTAATAGGCGCATCGCTGGATTAC<br>TTATTGCTGGCTTTTTCAAGTGCGCTTTGGAT<br>GCTGTATTTAGGCCGTTTGCTTTCAGGGATCA<br>CAGGAGCTACTGGGGCTGTCGCGGCATCGG<br>TCATTGCCGATACCACCTCAGCTTCTCAACGC<br>GTGAAGTGGTTCGGTTGGTTAGGGGCAAGTT | Tetracycline | JNH5 | JNH6 | GAAGTGACCGGCGATCAAATG | GCCTGGGCAATGATCAATATGC |

| | | | | | | |
|---|---|---|---|---|---|---|
| | TTGGGCTTGGTTTAATAGCGGGGCCTATTATT GGTGGTTTTGCAGGAGAGATTTCACCGCATA GTCCCTTTTTTATCGCTGCGTTGCTAAATATTG TCACTTTCCTTGTGGTTATGTTTTGGTTCCGT GAAACCAAAAATACACGTGATAATACAGATACC GAAGTAGGGGTTGAGACGCAATCGAATTCGG TATACATCACTTTATTTAAAACGATGCCCATTTT GTTGATTATTTATTTTTCAGCGCAATTGATAGG CCAAATTCCCGCAACGGTGTGGGTGCTATTTA CCGAAAATCGTTTTGGATGGAATAGCATGATG GTTGGCTTTTCATTAGCGGGTCTTGGTCTTTT ACACTCAGTATTCCAAGCCTTTGTGGCAGGAA GAATAGCCACTAAATGGGGCGAAAAAACGGC AGTACTGCTCGGATTTATTGCAGATAGTAGTG CATTTGCCTTTTTAGCGTTTATATCTGAAGGTT GGTTAGTTTTCCCTGTTTTAATTTTATTGGCTG GTGGTGGGATCGCTTTACCTGCATTACAGGG AGTGATGTCTATCCAAACAAAGAGTCATCAGC AAGGTGCTTTACAGGGATTATTGGTGAGCCTT ACCAATGCAACCGGTGTTATTGGCCCATTACT GTTTGCTGTTATTTATAATCATTCACTACCAATT TGGGATGGCTGGATTTGGATTATTGGTTTAGC GTTTTACTGTATTATTATCCTGCTATCGATGACC TTCATGTTAACCCCTCAAGCTCAGGGGAGTAA ACAGGAGACAAGTGCTTAGTTATTTCGTCACC AAATGATTTTTACAAATCAGTAACAAAAGTAAA GAAGGCGACACCATGCGACTATGG | | | | | |
| trxB-sfGFP-tetR | ATGGATCACATTTATCGCCAGGCCATTACTTC GGCCGGTACAGGCTGCATGGCAGCACTTGAT GCGGAACGCTACCTCGATGGTTTAGCTGACG CAAAAGGGGGTTCAGGCGGGTCGGGTGGCgt gagcaagggcgaggagctgttcaccggggtggtgcccatcctgg tcgagctggacggcgacgtaaacggccacaagttcagcgtgcg cggcgagggcgagggcgatgccaccaacggcaagctgaccct gaagttcatctgcaccaccggcaagctgcccgtgccctggcccac cctcgtgaccaccctgacctacggcgtgcagtgcttcagccgctac cccgaccacatgaagcgccacgacttcttcaagtccgccatgccc gaaggctacgtccaggagcgcaccatcagcttcaaggacgacg gcacctacaagacccgcgccgaggtgaagttcgagggcgacac cctggtgaaccgcatcgagctgaagggcatcgacttcaaggagg acggcaacatcctggggcacaagctggagtacaacttcaacag ccacaacgtctatatcaccgccgacaagcagaagaacggcatc aaggccaacttcaagatccgccacaacgtggaggacggcagc | Tetracycline | JNH5 | JNH6 | GAAGTGACCGGCGATCAAATG | GCCTGGGCAATGATCAATATGC |

| | |
|---|---|
| | gtgcagctcgccgaccactaccagcagaacacccccatcggcg<br>acggccccgtgctgctgcccgacaaccactacctgagcacccag<br>tccgtgctgagcaaagaccccaacgagaagcgcgatcacatgg<br>tcctgctggagttcgtgaccgccgccgggatcactcacggcatgg<br>acgagctgtacaagtaatgaTCCTAATTTTTGTTGACA<br>CTCTATCATTGATAGAGTTATTTTACCACTCCC<br>TATCAGTGATAGAGAAAAGTGAAATGAATAGTT<br>CGACAAAGATCGCATTGGTAATTACGTTACTC<br>GATGCCATGGGGATTGGCCTTATCATGCCAGT<br>CTTGCCAACGTTATTACGTGAATTTATTGCTTC<br>GGAAGATATCGCTAACCACTTTGGCGTATTGC<br>TTGCACTTTATGCGTTAATGCAGGTTATCTTTG<br>CTCCTTGGCTTGGAAAAATGTCTGACCGATTT<br>GGTCGGCGCCCAGTGCTGTTGTTGTCATTAAT<br>AGGCGCATCGCTGGATTACTTATTGCTGGCTT<br>TTTCAAGTGCGCTTTGGATGCTGTATTTAGGC<br>CGTTTGCTTTCAGGGATCACAGGAGCTACTG<br>GGGCTGTCGCGGCATCGGTCATTGCCGATAC<br>CACCTCAGCTTCTCAACGCGTGAAGTGGTTC<br>GGTTGGTTAGGGGCAAGTTTTGGGCTTGGTT<br>TAATAGCGGGGCCTATTATTGGTGGTTTTGCA<br>GGAGAGATTTCACCGCATAGTCCCTTTTTTAT<br>CGCTGCGTTGCTAAATATTGTCACTTTCCTTGT<br>GGTTATGTTTTGGTTCCGTGAAACCAAAAATA<br>CACGTGATAATACAGATACCGAAGTAGGGGTT<br>GAGACGCAATCGAATTCGGTATACATCACTTTA<br>TTTAAAACGATGCCCATTTTGTTGATTATTTATT<br>TTTCAGCGCAATTGATAGGCCAAATTCCCGCA<br>ACGGTGTGGGTGCTATTTACCGAAAATCGTTT<br>TGGATGGAATAGCATGATGGTTGGCTTTTCAT<br>TAGCGGGTCTTGGTCTTTTACACTCAGTATTC<br>CAAGCCTTTGTGGCAGGAAGAATAGCCACTA<br>AATGGGGCGAAAAAACGGCAGTACTGCTCGG<br>ATTTATTGCAGATAGTAGTGCATTTGCCTTTTT<br>AGCGTTTATATCTGAAGGTTGGTTAGTTTTCCC<br>TGTTTTAATTTTATTGGCTGGTGGTGGGATCG<br>CTTTACCTGCATTACAGGGAGTGATGTCTATC<br>CAAACAAAGAGTCATCAGCAAGGTGCTTTACA<br>GGGATTATTGGTGAGCCTTACCAATGCAACCG<br>GTGTTATTGGCCCATTACTGTTTGCTGTTATTT<br>ATAATCATTCACTACCAATTTGGGATGGCTGGA<br>TTTGGATTATTGGTTTAGCGTTTTACTGTATTAT<br>TATCCTGCTATCGATGACCTTCATGTTAACCCC<br>TCAAGCTCAGGGGAGTAAACAGGAGACAAGT |

| | | | | | | |
|---|---|---|---|---|---|---|
| | GCTTAGTTATTTCGTCACCAAATGATTTTTACA AATCAGTAACAAAAGTAAAGAAGGCGACACCA TGCGACTATGG | | | | | |
| phoA_evr1 p_tetR | ATGGATCACATTTATCGCCAGGCCATTACTTC GGCCGGTACAGGCTGCATGGCAGCACTTGAT GCGGAACGCTACCTCGATGGTTTAGCTGACG CAAAAtaaTGACAAATAAAACGAAAGGCTCAGT CGAAAGACTGGGCCTTTCGTTTTATGTTAATC TTTTCAACAGCTGTCATAAAGTTGTCACGGCC GAGACTTATAGTCGCTTTGTTTTTATTTTTTAAT GTATTTGTATCTAGAGATTAAAGAGGAGAATAC TAGATGAAGGCAATTGATAAGATGACCGACAA TCCTCCTCAGGAAGGCTTGTCCGGGCGCAAA ATCATCTATGATGAGGACGGGAAGCCGTGCC GTTCCTGCAATACTTTGCTGGACTTCCAATAC GTCACGGGTAAAATCTCGAATGGATTGAAGAA TTTGTCTTCAAATGGAAAACTGGCAGGGACTG GGGCTCTTACGGGGGAGGCATCGGAACTTAT GCCTGGTTCCCGCACTTATCGCAAAGTAGATC CACCTGATGTTGAGCAGCTTGGCCGTTCTTC GTGGACCTTGTTGCACTCTGTCGCCGCGTCG TATCCAGCTCAGCCTACGGACCAACAGAAGG GCGAGATGAAACAGTTCCTGAACATCTTCTCG CACATTTACCCTTGCAACTGGTGCGCCAAGG ACTTCGAGAAGTACATTCGCGAGAATGCGCC GCAAGTAGAATCCCGCGAAGAATTGGGACGT TGGATGTGTGAAGCACATAATAAGGTGAATAA GAAGTTACGCAAACCAAAGTTCGACTGTAATT TCTGGGAAAAACGCTGGAAAGACGGGTGGG ACGAGTAATGACAAATAAAACGAAAGGCTCAG TCGAAAGACTGGGCCTTTCGTTTTATTCCTAAT TTTTGTTGACACTCTATCATTGATAGAGTTATTT TACCACTCCCTATCAGTGATAGAGAAAAGTGA AATGAATAGTTCGACAAAGATCGCATTGGTAAT TACGTTACTCGATGCCATGGGGATTGGCCTTA TCATGCCAGTCTTGCCAACGTTATTACGTGAA TTTATTGCTTCGGAAGATATCGCTAACCACTTT GGCGTATTGCTTGCACTTTATGCGTTAATGCA GGTTATCTTTGCTCCTTGGCTTGGAAAAATGT CTGACCGATTTGGTCGGCGCCCAGTGCTGTT GTTGTCATTAATAGGCGCATCGCTGGATTACTT ATTGCTGGCTTTTTCAAGTGCGCTTTGGATGC TGTATTTAGGCCGTTTGCTTTCAGGGATCACA | Tetracycline | JNH3 | JNH4 | GAAGTGACCGGCGATCAAATG | GCCTGGGCAATGATCAATATGC |

| | | | | | | |
|---|---|---|---|---|---|---|
| | GGAGCTACTGGGGCTGTCGCGGCATCGGTC<br>ATTGCCGATACCACCTCAGCTTCTCAACGCGT<br>GAAGTGGTTCGGTTGGTTAGGGGCAAGTTTT<br>GGGCTTGGTTTAATAGCGGGGCCTATTATTGG<br>TGGTTTTGCAGGAGAGATTTCACCGCATAGTC<br>CCTTTTTTATCGCTGCGTTGCTAAATATTGTCA<br>CTTTCCTTGTGGTTATGTTTTGGTTCCGTGAA<br>ACCAAAAATACACGTGATAATACAGATACCGAA<br>GTAGGGGTTGAGACGCAATCGAATTCGGTATA<br>CATCACTTTATTTAAAACGATGCCCATTTTGTT<br>GATTATTTATTTTTCAGCGCAATTGATAGGCCA<br>AATTCCCGCAACGGTGTGGGTGCTATTTACCG<br>AAAATCGTTTTGGATGGAATAGCATGATGGTT<br>GGCTTTTCATTAGCGGGTCTTGGTCTTTTACA<br>CTCAGTATTCCAAGCCTTTGTGGCAGGAAGAA<br>TAGCCACTAAATGGGGCGAAAAAACGGCAGT<br>ACTGCTCGGATTTATTGCAGATAGTAGTGCATT<br>TGCCTTTTTAGCGTTTATATCTGAAGGTTGGTT<br>AGTTTTCCCTGTTTTAATTTTATTGGCTGGTGG<br>TGGGATCGCTTTACCTGCATTACAGGGAGTGA<br>TGTCTATCCAAACAAAGAGTCATCAGCAAGGT<br>GCTTTACAGGGATTATTGGTGAGCCTTACCAA<br>TGCAACCGGTGTTATTGGCCCATTACTGTTTG<br>CTGTTATTTATAATCATTCACTACCAATTTGGGA<br>TGGCTGGATTTGGATTATTGGTTTAGCGTTTTA<br>CTGTATTATTATCCTGCTATCGATGACCTTCAT<br>GTTAACCCCTCAAGCTCAGGGGAGTAAACAG<br>GAGACAAGTGCTTAGTTATTTCGTCACCAAAT<br>GATTTTTACAAATCAGTAACAAAAGTAAAGAAG<br>GCGACACCATGCGACTATGG | | | | | |
| trxB-das4-phoA_evr1p_tetR | ATGGATCACATTTATCGCCAGGCCATTACTTC<br>GGCCGGTACAGGCTGCATGGCAGCACTTGAT<br>GCGGAACGCTACCTCGATGGTTTAGCTGACG<br>CAAAAGCGGCCAACGATGAAAACTATTCTGAA<br>AACTATGCGGATGCGTCTtaaTGACAAATAAAA<br>CGAAAGGCTCAGTCGAAAGACTGGGCCTTTC<br>GTTTTATGTTAATCTTTTCAACAGCTGTCATAA<br>AGTTGTCACGGCCGAGACTTATAGTCGCTTTG<br>TTTTTATTTTTTAATGTATTTGTATCTAGAGATTA<br>AAGAGGAGAATACTAGATGAAGGCAATTGATA<br>AGATGACCGACAATCCTCCTCAGGAAGGCTT<br>GTCCGGGCGCAAAATCATCTATGATGAGGAC<br>GGGAAGCCGTGCCGTTCCTGCAATACTTTGC | Tetracycline | JNH3 | JNH4 | GAAGTGACCGGCGATCAAATG | GCCTGGGCAATGATCAATATGC |

| | |
|---|---|
| | TGGACTTCCAATACGTCACGGGTAAAATCTCG AATGGATTGAAGAATTTGTCTTCAAATGGAAAA CTGGCAGGGACTGGGGCTCTTACGGGGGAG GCATCGGAACTTATGCCTGGTTCCCGCACTTA TCGCAAAGTAGATCCACCTGATGTTGAGCAG CTTGGCCGTTCTTCGTGGACCTTGTTGCACT CTGTCGCCGCGTCGTATCCAGCTCAGCCTAC GGACCAACAGAAGGGCGAGATGAAACAGTTC CTGAACATCTTCTCGCACATTTACCCTTGCAA CTGGTGCGCCAAGGACTTCGAGAAGTACATT CGCGAGAATGCGCCGCAAGTAGAATCCCGCG AAGAATTGGGACGTTGGATGTGTGAAGCACA TAATAAGGTGAATAAGAAGTTACGCAAACCAA AGTTCGACTGTAATTTCTGGGAAAAACGCTGG AAAGACGGGTGGGACGAGTAATGACAAATAA AACGAAAGGCTCAGTCGAAAGACTGGGCCTT TCGTTTTATTCCTAATTTTTGTTGACACTCTATC ATTGATAGAGTTATTTTACCACTCCCTATCAGT GATAGAGAAAAGTGAAATGAATAGTTCGACAA AGATCGCATTGGTAATTACGTTACTCGATGCC ATGGGGATTGGCCTTATCATGCCAGTCTTGCC AACGTTATTACGTGAATTTATTGCTTCGGAAGA TATCGCTAACCACTTTGGCGTATTGCTTGCAC TTTATGCGTTAATGCAGGTTATCTTTGCTCCTT GGCTTGGAAAAATGTCTGACCGATTTGGTCG GCGCCCAGTGCTGTTGTTGTCATTAATAGGCG CATCGCTGGATTACTTATTGCTGGCTTTTTCAA GTGCGCTTTGGATGCTGTATTTAGGCCGTTTG CTTTCAGGGATCACAGGAGCTACTGGGGCTG TCGCGGCATCGGTCATTGCCGATACCACCTC AGCTTCTCAACGCGTGAAGTGGTTCGGTTGG TTAGGGGCAAGTTTTGGGCTTGGTTTAATAGC GGGGCCTATTATTGGTGGTTTTGCAGGAGAG ATTTCACCGCATAGTCCCTTTTTTATCGCTGCG TTGCTAAATATTGTCACTTTCCTTGTGGTTATG TTTTGGTTCCGTGAAACCAAAAATACACGTGA TAATACAGATACCGAAGTAGGGGTTGAGACGC AATCGAATTCGGTATACATCACTTTATTTAAAAC GATGCCCATTTTGTTGATTATTTATTTTTCAGC GCAATTGATAGGCCAAATTCCCGCAACGGTGT GGGTGCTATTTACCGAAAATCGTTTTGGATGG AATAGCATGATGGTTGGCTTTTCATTAGCGGG TCTTGGTCTTTTACACTCAGTATTCCAAGCCTT TGTGGCAGGAAGAATAGCCACTAAATGGGGC |

| | | | | | | |
|---|---|---|---|---|---|---|
| | GAAAAAACGGCAGTACTGCTCGGATTTATTGC<br>AGATAGTAGTGCATTTGCCTTTTTAGCGTTTAT<br>ATCTGAAGGTTGGTTAGTTTTCCCTGTTTTAAT<br>TTTATTGGCTGGTGGTGGGATCGCTTTACCTG<br>CATTACAGGGAGTGATGTCTATCCAAACAAAG<br>AGTCATCAGCAAGGTGCTTTACAGGGATTATT<br>GGTGAGCCTTACCAATGCAACCGGTGTTATTG<br>GCCCATTACTGTTTGCTGTTATTTATAATCATTC<br>ACTACCAATTTGGGATGGCTGGATTTGGATTAT<br>TGGTTTAGCGTTTTACTGTATTATTATCCTGCTA<br>TCGATGACCTTCATGTTAACCCCTCAAGCTCA<br>GGGGAGTAAACAGGAGACAAGTGCTTAGTTAT<br>TTCGTCACCAAATGATTTTTACAAATCAGTAAC<br>AAAAGTAAAGAAGGCGACACCATGCGACTAT<br>GG | | | | | |
| Autolysis_<br>apraR | AAGAAATTAAAAAACGAAAATACCAGCTATAGC<br>CAGATTGTCACAGAGTGTCGTATGCGTTACGC<br>CGT<br>ACAGATGTTATTGATGGATAACAAAAATATCAC<br>TCAGGTGGCGCAATTATGTGGCTATAGCAGCA<br>CGT<br>CGTACTTTATCTCTGTTTTTAAGGCGTTTTACG<br>GCCTGACACCGTTGAATTATCTCGCCAAACAG<br>CGAC<br>AAAAAGTGATGTGGtgaAGGGCAAAGCGGAAA<br>CGGATAAGACGGGCATAAATGAGGAAGAAAT<br>GGCT<br>CGACCTAGCATAACCCCGCGGGGCCTCTTCG<br>GGGGTCTCGCGGGGTTTTTTGCTGAAAGAAG<br>CTTCAA<br>ATAAAACGAAAGGCTCAGTCGAAAGACTGGG<br>CCTTTCGTTTTATCTGTTGTTTGTCGCTGCGG<br>CCGGGT<br>CAGGTATGATTTAAATGGTCAGTAACGGGTCT<br>TGAGGGGTTTTTTGCATATGTGCGTAATTGTG<br>CTGAT<br>CTCTTATATAGCTGCTCTCATTATCTCTCTACC<br>CTGAAGTGACTCTCTCACCTGTAAAAATAATAT<br>CTCA | Apra<br>mycin | JNH5 | JNH6 | GCATT<br>GCTTTT<br>TACCG<br>TATTGT<br>CTAAC | GATTATT<br>ATGGTG<br>TCACGC<br>CATCTC |

| | | | | | | |
|---|---|---|---|---|---|---|
| yibD-roGFP2 | CTGGAAAAAGGAGATATACCATGGTGAGCAAGGGCGAGGAGCTGTTCACCGGGGTGGTGCCCATCCTGGTCGAGCTGGACGGCGACGTAAACGGCCACAAGTTCAGCGTGTCCGGCGAGGGCGAGGGCGATGCCACCTACGGCAAGCTGACCCTGAAGTTCATCTCCACCACCGGCAAGCTGCCCGTGCCCTGGCCCACCCTCGTGACCACCCTGACCTACGGCGTGCAGTGCTTCAGCCGCTACCCCGACCACATGAAGCAGCACGACTTCTTCAAGTCCGCCATGCCCGAAGGCTACGTCCAGGAGCGCACCATCTTCTTCAAGGACGACGGCAACTACAAGACCCGCGCCGAGGTGAAGTTCGAGGGCGACACCCTGGTGAACCGCATCGAGCTGAAGGGCATCGACTTCAAGGAGGACGGCAACATCCTGGGGCACAAGCTGGAGTACAACTACAACTGCCACAACGTCTATATCATGGCCGACAAGCAGAAGAACGGCATCAAGGTGAACTTCAAGATCCGCCACAACATCGAGGACGGCAGCGTGCAGCTCGCCGACCACTACCAGCAGAACACCCCCATCGGCGACGGCCCCGTGCTGCTGCCCGACAACCACTACCTGAGCACCTGCTCCGCCCTGAGCAAAGACCCCAACGAGAAGCGCGATCACATGGTCCTGCTGGAGTTCGTGACCGCCGCCGGGATCACTCTCGGCATGGACGAGCTGTACAAGTAATGAGCTCTTCTAATACGACTCAC | - | SL1 | SR2 | CAGTCCAGTTACGCTGGAGTC | GGTCAGGTATGATTTAAATGGTCAGT |
| pCOLA-Gent-ppiB | GAAGTGCCAGACGATGGAAGGACGCAATGAAGCTGGAGGATTGGCGTGGGAATCGTGCTTCTGTCTAAGCAAGAATGCCTAGCGTACAGGGTGCACTTTGTAACGATTTGGGAGTCCAGAGACTCGCTGTTTTCGAAATTctcggtcgctacgctccgggcgtgagactATGCAGCGCAGAAACGTCCTAGAAGATGCCAGGAGGATACTTAGCAGAGAGACAATAAGGCCGGAGCGAAGCCGTTTTTCCATAGGCTCCGCCCCCCTGACGAACATCACGAAATCTGACGCTCAAATCAGTGGTGGCGAAACCCGACAGGACTATAAAGATACCAGGCGTTTCCCCCTGATGGCTCCCTCTTGCGCTCTCCTGTTCCCGTCCTGCGGCGTCCGTGTTGTGGTGGAGGCTTTACCCAAATCACCACGTCCCGTTCCGTGTAGACAGTTCGCTCCAAGCTGGGCTGTGTGCAAGAACCCCCCGTTCAGCCCGACTGCTGCGCCTTATCCGGTAACTATCATCTTGAGTCCAACCCGGAAAGACACGACAAAACGCCACTGGCAGCAGCCATTGG | Gentamicin | JNH7 | JNH8 | GTGCGTAATTGTGCTGATCTCTT | AAGTTGGAACCTCTTACGTGC |

| | |
|---|---|
| | TAACTGAGAATTAGTGGATTTAGATATCGAGAG<br>TCTTGAAGTGGTGGCCTAACAGAGGCTACAC<br>TGAAAGGACAGTATTTGGTATCTGCGCTCCAC<br>TAAAGCCAGTTACCAGGTTAAGCAGTTCCCCA<br>ACTGACTTAACCTTCGATCAAACCGCCTCCCC<br>AGGCGGTTTTTTCGTTTACAGAGCAGGAGATT<br>ACGACGATCGTAAAAGGATCTCAAGAAGATCC<br>TTTACGGATTCCCGACACCATCACTCTAGATTT<br>CAGTGCAATTTATCTCTTCAAATGTAGCACCTG<br>AAGTCAGCCCCATACGATATAAGTTGTAATTCT<br>CATGTTAGTCATGCCCCGCGCCCACCGGAAG<br>GAGCTGACTGGGTTGAAGGCTCTCAAGGGCA<br>TCGGTCGAGATCCCGGTGCCTAATGAGTGAG<br>CTAACTTACATTAATTGCGTGTGCGTAATTGTG<br>CTGATCTCTTATATAGCTGCTCTCATTATCTCT<br>CTACCCTGAAGTGACTCTCTCACCTGTAAAAA<br>TAATATCTCACAGGCTTAATAGTTTCTTAATACA<br>AAGCCTGTAAAACGTCAGGATAACTTCTGTGT<br>AGGAGGATAATCTATGGTAACATTCCACACTAA<br>CCACGGGGACATCGTTATTAAAACATTCGATG<br>ATAAAGCTCCGGAAACAGTCAAGAACTTTCTG<br>GACTATTGTCGCGAGGGGTTCTACAATAATAC<br>AATCTTCCACCGTGTTATTAACGGGTTCATGAT<br>CCAAGGCGGAGGGTTCGAGCCGGGCATGAA<br>GCAGAAAGCTACTAAAGAGCCAATTAAGAACG<br>AAGCAAACAATGGTCTGAAGAACACTCGTGG<br>TACGCTTGCGATGGCCCGTACCCAAGCGCCA<br>CATAGCGCCACCGCCCAATTCTTTATTAATGTT<br>GTGGATAATGATTTTCTGAACTTCTCCGGAGA<br>ATCGTTACAGGGTTGGGGCTACTGCGTTTTC<br>GCAGAAGTCGTCGACGGTATGGATGTCGTGG<br>ATAAGATCAAAGGAGTTGCGACGGGTCGTAG<br>CGGTATGCACCAAGACGTGCCGAAGGAGGAT<br>GTGATTATCGAGTCTGTGACTGTTTCTGAATAA<br>GGATCCgagtcacactggctcaccttcgggtgggcctttctgcg<br>tttatTGATCGGCACGTAAGAGGTTCCAACTTTC<br>ACCATAATGAAATAAGATCACTACCGGGCGTAT<br>TTTTTGAGTTATCGAGATTTTCAGGAGCTGGA<br>AGGAAGTAAAAATGTTGCGTAGCTCTAACGAT<br>GTGACGCAACAAGGTTCGCGTCCAAAGACAA<br>AATTGGGAGGCAGTAGCATGGGGATCATTCG<br>CACTTGTCGCCTGGGGCCAGACCAGGTGAA<br>GTCAATGCGTGCGGCTCTGGACTTATTCGGG<br>CGCGAATTTGGAGATGTAGCCACTTACTCACA |

| | | | | | | |
|---|---|---|---|---|---|---|
| | GCACCAACCGGACAGTGATTACTTGGGGAAT TTACTTCGCAGTAAAACTTTTATCGCTTTGGCC GCTTTCGACCAGGAGGCTGTAGTAGGTGCGT TGGCAGCCTATGTTCTTCCTAAATTCGAGCAA CCGCGTAGCGAAATTTACATCTATGATCTTGCA GTCTCCGGCGAACATCGCCGTCAGGGGATCG CCACAGCTTTAATCAACCTTTTGAAGCATGAG GCTAATGCACTTGGAGCGTACGTGATTTATGT GCAGGCTGACTACGGTGATGATCCTGCAGTC GCTCTGTACACCAAACTGGGTATCCGCGAGG AGGTCATGCACTTTGATATTGACCCGTCTACG GCTACCTAATAGAAGCTT | | | | | |
| PDI | GGGTGGGACGAGTAATGATCTAGAGATTAAAG AGGAGAATACTAGATGGATGCGCCAGAAGAA GAAGATCATGTTCTTGTGCTTCGTAAATCCAAT TTCGCGGAAGCTTTGGCAGCGCATAAATATTT GCTGGTTGAATTCTACGCCCCTTGGTGTGGT CATTGCAAGGCGTTGGCGCCCGAGTATGCGA AGGCTGCTGGCAAGCTTAAAGCGGAGGGCTC GGAAATCCGTTTAGCGAAAGTGGATGCCACG GAGGAATCAGACCTTGCACAACAGTACGGCG TCCGCGGTTATCCGACGATTAAATTCTTCCGC AATGGAGATACAGCATCCCCAAAGGAGTACAC GGCTGGGCGTGAGGCAGATGACATTGTCAAT TGGCTTAAAAAACGCACAGGCCCAGCGGCTA CGACTCTGCCAGATGGTGCTGCGGCAGAAAG TCTTGTTGAATCGAGTGAAGTTGCTGTAATTG GATTTTTTAAGGACGTCGAGTCAGATAGTGCT AAGCAATTTTTACAAGCCGCCGAAGCAATCGA CGACATTCCTTTCGGCATCACAAGTAATTCTG ATGTCTTCTCGAAGTACCAATTGGACAAGGAT GGAGTCGTGTTATTTAAAAAGTTCGATGAGGG CCGTAATAATTTCGAAGGCGAGGTGACTAAGG AGAACTTATTAGATTTCATCAAACACAACCAAC TTCCTCTTGTCATTGAATTCACTGAGCAAACT GCGCCGAAGATTTTCGGGGGAGAAATTAAGA CCCACATCTTGTTGTTTCTGCCGAAATCAGTT TCGGACTACGATGGGAAACTGAGCAATTTTAA GACTGCGGCGGAATCATTTAAAGGGAAGATTT TATTCATCTTCATTGACAGTGACCACACTGAC AACCAGCGTATTCTGGAGTTCTTCGGATTAAA GAAAGAGGAATGTCCAGCCGTACGTTTGATTA CTCTGGAGGAGGAGATGACTAAATACAAACC | - | JNH30 | JNH8 | CTCAAGGGCATCGGTCGAGA | AAGTTGGAACCTCTTACGTGC |

| | | | | | | |
|---|---|---|---|---|---|---|
| | CGAGTCTGAAGAGCTGACAGCGGAGCGTATT<br>ACCGAGTTCTGCCATCGCTTTTTAGAAGGCAA<br>AATTAAGCCACATCTTATGTCGCAAGAACTGC<br>CGGAAGACTGGGACAAGCAGCCTGTCAAGG<br>TGCTTGTCGGAAAAAACTTCGAGGATGTTGC<br>GTTTGACGAGAAGAAGAATGTCTTTGTTGAGT<br>TCTACGCCCCCTGGTGCGGGCATTGTAAACA<br>GTTGGCCCCCATTTGGGACAAGCTTGGCGAG<br>ACGTACAAAGACCATGAGAATATTGTAATTGC<br>GAAAATGGATTCAACCGCAAATGAGGTCGAA<br>GCTGTGAAAGTACACAGCTTCCCAACACTGA<br>AATTTTTTCCTGCATCGGCCGATCGCACAGTT<br>ATCGACTACAACGGAGAACGTACCCTGGACG<br>GCTTTAAAAAATTCCTGGAGTCTGGGGGTCA<br>GGATGGTGCTGGGGATGACGACGACTTAGAG<br>GACCTTGAGGAAGCCGAAGAACCCGACATGG<br>AGGAGGATGATGATCAGAAGGCGGTTAAAGA<br>TGAGTTGTAATGAGGATCCgagtcacactgg | | | | | |
| VHH72 | ATGCAGGTGCAGTTACAGGAATCCGGCGGCG<br>GCCTTGTTCAAGCTGGTGGATCGCTTCGCTTA<br>TCCTGCGCCGCCTCGGGCCGCACTTTCTCG<br>GAATACGCAATGGGCTGGTTCCGCCAAGCAC<br>CGGGGAAAGAGCGCGAGTTCGTAGCTACTAT<br>TTCATGGTCCGGGGGGTCCACCTATTATACCG<br>ATTCAGTAAAGGGCCGTTTCACTATTTCCCGC<br>GATAATGCGAAGAATACTGTTTACTTGCAGATG<br>AACTCGCTTAAGCCCGATGATACCGCAGTATA<br>TTACTGTGCAGCAGCAGGCCTTGGGACGGTC<br>GTATCTGAATGGGACTACGATTATGACTATTGG<br>GGACAGGGTACTCAAGTTACTGTTTCGAGCG<br>GCTCGCACCATCATCATCACCACTAATGAGGA<br>TCCCCGGC | - | SL1 | SR2 | CAGTC<br>CAGTT<br>ACGCT<br>GGAGT<br>C | GGTCAG<br>GTATGA<br>TTTAAAT<br>GGTCAG<br>T |
| VHH GFP enhancer | TTCTGTGTAGGAGGATAATCTATGATGGCACA<br>AGTTCAGTTGGTAGAAAGTGGAGGTGCCCTG<br>GTCCAACCCGGGGGTTCTCTTCGCTTGTCAT<br>GTGCTGCAAGCGGGTTTCCGGTGAATCGCTA<br>CTCGATGCGTTGGTATCGCCAAGCACCCGGC<br>AAGGAGCGCGAATGGGTCGCAGGCATGTCGT<br>CCGCAGGCGATCGTAGTAGTTACGAGGATAG<br>TGTCAAAGGTCGCTTTACGATCAGCCGCGAT<br>GACGCTCGCAATACCGTTTACTTACAAATGAA<br>CAGCCTTAAGCCAGAAGATACCGCAGTCTACT<br>ACTGCAATGTAAACGTGGGCTTTGAATACTGG | - | SL1 | SR2 | CAGTC<br>CAGTT<br>ACGCT<br>GGAGT<br>C | GGTCAG<br>GTATGA<br>TTTAAAT<br>GGTCAG<br>T |

| | | | | | | |
|---|---|---|---|---|---|---|
| | GGCCAGGGTACTCAAGTTACGGTTTCCAGTCACCACCACCACCATCATTAATGAGGATCCCCGGC | | | | | |
| VHH GFP minimizer | TTCTGTGTAGGAGGATAATCTATGGCCGATGTCCAGCTGCAAGAGTCCGGCGGTGGGAGCGTTCAAGCAGGTGGGAGTCTGCGCCTTTCGTGCGCAGCCTCAGGAGATACATTTAGTTCATATTCTATGGCGTGGTTCCGTCAAGCACCGGGGAAGGAATGTGAACTTGTCAGCAACATTCTGCGCGACGGCACCACAACATACGCCGGTAGCGTAAAGGGACGTTTCACAATTAGCCGCGACGATGCAAAGAACACGGTCTACTTACAAATGGTCAATCTGAAATCTGAAGACACCGCCCGTTACTACTGCGCGGCTGACAGTGGAACTCAGCTGGGCTATGTCGGGGCTGTAGGATTGTCGTGTTTAGACTACGTTATGGATTATTGGGGAAAAGGTACACAGGTAACTGTTAGTAGTCATCATCATCACCACCACTAATGAGGATCCCCGGC | - | SL1 | SR2 | CAGTCCAGTTACGCTGGAGTC | GGTCAGGTATGATTTAAATGGTCAGT |
| VHH H6 antivenin | TTCTGTGTAGGAGGATAATCTATGCAAGTCCAGCTTCAAGAAAGTGGGGGCGGATTAGTTCAGGCGGGTGGGTCACTTACACTGGCATGCACTGCATCAGGCCGTACCTTCGACCGTTACGCGGTTGGTTGGTTCCGCCAGACTCCTGGAAAGGATCGTGAGTTTGTCGCAACGATCTCGTGGTCAGGAGGAACCACTCGTTACGCGGATAGCGTGAAGGGTCGCTTCACAGTGTCTCGCGACAACGCTAAGAACACTGTGTACCTTCAGATGAACACCTTGAAGCCAGAGGACACGGCCGTTTATTACTGCGCAGCTGACCTGGCGTTGTCCACGGTCGATGAAGCGGTCGCACGCTATTGGGGGCAGGGGACTCAAGTAACAGTATCTTCCCATCACCATCACCACCACTAATGAGGATCCCCGGC | - | SL1 | SR2 | CAGTCCAGTTACGCTGGAGTC | GGTCAGGTATGATTTAAATGGTCAGT |
| VHH VCAM1 | TTCTGTGTAGGAGGATAATCTATGCAAGTCCAACTTCAGGAGTCAGGAGGAGGCTCAGTGCAGACTGGTGGATCTCTGCGCTTGTCGTGTGCGGCCAGCGGCTACACAAACAGCATCATGTATATGGCGTGGTTCCGCCAGGCTCCGGGGAAAAAGCGCGAAGGAGTTGCAGCGATCCGTTTTCCCGACGATTCAGCCTACTATGCGGGTTCGGTTAAAGGACGTTTTACTATCTCTCACGACAACGCCAAAAACACGGTTTACTTGCAGATGAACAACCTTAACCCTGAAGACACAGCAATGTATTACTGTGCAGCTCGCTCATCACCTTACAGTTTTGCGTGGA | - | SL1 | SR2 | CAGTCCAGTTACGCTGGAGTC | GGTCAGGTATGATTTAAATGGTCAGT |

| | | | | | | |
|---|---|---|---|---|---|---|
| | ACGACCCTAGCAACTATAACTACTGGGGCCAGGGCACGCAGGTCACCGTAAGCAGCCATCATCACCACCACCATTAATGAGGATCCCCGGC | | | | | |
| VHH 1B5 | TTCTGTGTAGGAGGATAATCTATGGAGGTTCAACTTGTTGAGAGCGGGGGTGGGCTGGTACAACCGGGTGGATCACTGCGCCTGAGCTGCGCCGCGAGTGGCCGTGCTACATTTGATGATTACGCTATTGGCTGGTTCCGCCAAGCACCGGGAAAGGAGCGTGAGGGCGTATCTTATATTGGATGTAATGATGGCGCTACCTATTATGCGGGCAGCGTCAAAGGCCGCTTTACCATTAGTTGTGATTACGCGAAAAATACGGTCTATTTACAGATGAATTCCTTAAAACCTGAGGATACTGCAGTTTACTATTGTGCAGCGGCAGCTCAATGGGCAACTATCCGCTGGATTCACGAGTATGACTACAATATCTGGGGCCAGGGGACTCAGGTCACGGTGTCTAGCCATCACCATCATCACCACTAATGAGGATCCCCGGC | - | SL1 | SR2 | CAGTCCAGTTACGCTGGAGTC | GGTCAGGTATGATTTAAATGGTCAGT |
| VHH 2E7 | TTCTGTGTAGGAGGATAATCTATGGAGGTACAATTAGTAGAATCTGGCGGTGGGTTAGTACAGCCAGGTGGGTCCTTGCGCTTATCATGCGCCGCCTCAGGCAATATCGTCTCCATCGACGCTGCTGGCTGGTTCCGCCAAGCCCCTGGTAAACAACGCGAACCTGTCGCTACAATCTTGACGGGTGGGGCTACTAACTACGCTGACTCCGTCAAAGGTCGCTTCACGATCTCTCGCGACAATGCAAAAAACACTGTTTATCTGCAAATGAACTCTCTGAAGCCAGAAGATACCGCGGTGTATTATTGTTATGCACCGATGATCTATTATGGCGGCCGTTATAGCGACTATTGGGGCCAGGGCACACAAGTCACAGTATCGTCACACCACCATCATCACCACTAATGAGGATCCCCGGC | - | SL1 | SR2 | CAGTCCAGTTACGCTGGAGTC | GGTCAGGTATGATTTAAATGGTCAGT |
| VHH 3E3 | TTCTGTGTAGGAGGATAATCTATGGAGGTGCAACTTGTTGAGAGTGGCGGGGGATTGGTTCAACCTGGCGGTAGTTTGCGCTTATCGTGTGCGGCAAGCCAATTTACATTAGAATCATACGCCATTGGTTGGTTCCGCCAGGCTCCCGGAAAAGATAGTGAGGGTGTAGCTTGCATTAGCTCGTCTACTTACTATGCGGACTCTGTGAAGGGACGTTTTACAATCTCGCGCGATAACGCGAAAAACACGGTCTATCTTCAGATGGAAAGTCTTAAACCCGAAGACACTGCTGTCTACCACTGTGCCACAAGCGGGGCTGGCAGTTACTGCACACTTCGCGCCTTCGGGTCATGGGGTCAGGGCACGCAGGTCACAGTCA | - | SL1 | SR2 | CAGTCCAGTTACGCTGGAGTC | GGTCAGGTATGATTTAAATGGTCAGT |

| | | | | | | |
|---|---|---|---|---|---|---|
| | GTTCTCATCATCACCATCATCACTAATGAGGATCCCCGGC | | | | | |
| VHH C9 (BoNT-A) | TTCTGTGTAGGAGGATAATCTATGGAAGTACAACTGCAGGAATCCGGTGGTGGTGTCGTCCAACCGGGTGGTAGTCTGAAACTGTCTTGTAGTGGAAGCGGCGCTATTTTTGACACATACGACGTTGGGTGGTACCGTCAAGCTCCTGGGAAACGCCGTGAATTAGTTGCCTCAATCACCGCGACTGCCCGTGGTCGTATTGATTACAATTACTTCGCACAAGGCCGCTTTACGATTTCTAAAGATAATGCTGCAAATACTGTCTACTTGCAAATGGACCACCTTGAACCGGGCGACACCGCTGTGTATTACTGTAAGACGTCGGTTACCGGCTGGGGCCGCGGGACTCAAGTTACAGTAAGCTCGGCTATGCAGGGCACACTGCCCACACATCATCACCATCATCATTAATGAGGATCCCCGGC | - | SL1 | SR2 | CAGTCCAGTTACGCTGGAGTC | GGTCAGGTATGATTTAAATGGTCAGT |
| VHH E7 (BoNT-A) | TTCTGTGTAGGAGGATAATCTATGGATGTTCAGTTACAAGCGAGTGGTGGCGGTTCTGCGCAGGCTGGAGGTAGCTTAACCTTATCTTGTGCCGCCAGCGGCTTGATCTTTTCAAATTATATTATGGGCTGGTTCCGCCAGGCACCGGGTAAAGACCGCGAGTTCGTAGCTGCGATTAGTCGCCAAGGTACTACACCGTATTACGTCAACTCGGGAAATGACCGTTTCACAATTAGTCGTGACAACGCGAAAAACACTGGCACGTTACAAATGAATCATCTGAAGCCAGAGGACACCGCTGTATACTATTGCGCGGTCGATAGTTTTCTGAGCAACATCCGTGTCGGGAAGGCGGATTATTGGGGGCAGGGGACGCAGGTCACGCACCATCACCATCATCACTAATGAGGATCCCCGGC | - | SL1 | SR2 | CAGTCCAGTTACGCTGGAGTC | GGTCAGGTATGATTTAAATGGTCAGT |
| VHH JM3 | TTCTGTGTAGGAGGATAATCTATGGAAGTGCAGTTAGTCGAGAGCGGTGGAGGGCTGGTTCAGCCGGGTGGATCTTTGCGTTTGTCATGTGCGGCCAGTGGCTTCACACTTGAGGATTATTCCATCGGCTGGTTCCGCCAGGCACCGGGTAAAGAGCGCGAAGGCGTGAGTTGTATTTCTGACTCAGACGGACGTACATACTACGCCGATTCGGTCAAGGGCCGTTTTACCATTTCCCGCGATAACGCCAAGAACACGGTGTATCTTCAGATGAACTCTTTAAAGCCTGAAGATACGGCAGTCTACTATTGTGCGACAGATTGCACTGTAGTGCCAAGTTTGTTGTATGCGATGGATAGTGGAAAAGGAACCCAAGTAACAGTTAGTTCGCATCACCACCATCACCATTA | - | SL1 | SR2 | CAGTCCAGTTACGCTGGAGTC | GGTCAGGTATGATTTAAATGGTCAGT |

| | | | | | | |
|---|---|---|---|---|---|---|
| | ATGAGGATCCCCGGC | | | | | |
| VHH m36.4 | TTCTGTGTAGGAGGATAATCTATGCAAGTCCAGTTAGTACAAAGCGGAGGGGGCTTAGTACAACCGGGGGGCTCCCTGCGTTTGAGTTGTGCAGCGTCCGCGTTTGACTTTTCAGATTATGAGATGAGTTGGGTCCGCGAGGCTCCCGGCAAAGGCCTGGAATGGATCGGCGAGATTAATGATTCGGGTAATACAATCTATAATCCTTCGCTGAAATCTCGTGTTACGATCTCCCGTGACAATAGCAAGAACACTTTATATCTTCAGATGAATACGTTACGCGCTGAAGACACTGCTATCTATTACTGCGCTATTTACGGTGGCAATTCAGGCGGCGAATACTGGGGTCAAGGGACGTTAGTCACTGTCAGCAGTCACCATCATCACCATCACTAATGAGGATCCCCGGC | - | SL1 | SR2 | CAGTCCAGTTACGCTGGAGTC | GGTCAGGTATGATTTAAATGGTCAGT |
| VHH Re5D06 | TTCTGTGTAGGAGGATAATCTATGGGCTCTCAGGTACAGCTGGTTGAATCTGGGGGTGGCCTGGTTCAACCCGGAGGATCGCTTCGTTTGTCATGCGCAGCCTCCGGCATCACCCTGGATTATTATGCAATCGGTTGGTTTCGCCAAGCTCCAGGAAAAGAACGTGAGGGTGTTAGTCGCATCCGTTCATCAGATGGATCGACAAACTATGCAGACTCTGTGAAAGGGCGTTTTACGATGAGCCGCGATAACGCGAAGAACACGGTCTACTTGCAGATGAACTCTTTAAAGCCTGAGGACACGGCAGTATACTATTGTGCTTATGGCCCGTTAACGAAATACGGCTCAAGTTGGTATTGGCCTTATGAATATGACTACTGGGGACAGGGAACACAGGTCACTGTTTCAAGCACGTCCCATCATCATCATCACCACTAATGAGGATCCCCGGC | - | SL1 | SR2 | CAGTCCAGTTACGCTGGAGTC | GGTCAGGTATGATTTAAATGGTCAGT |
| VHH Re9F06 | TTCTGTGTAGGAGGATAATCTATGGGATCTCAAGTACAGTTGGTGGAATCAGGTGGTGGACTTGTTCAGGCGGGCGGGTCACTGCGCTTGAGCTGTGCTGCTAGTGGACGCACTTTCAGCAATGATGCGTTAGGATGGTTTCGCCAAGCACCTCGCAAGGAGCGTGAGTTTGTCGCGGCTATCAATTGGAATAGTGGCACCTATTATGCAGATTCTGTCAAAGGGCGTTTTACCATTTCGCGTGATAATGCTAAAAATACCGTGTACTTGCAAATGAACAGCTTAAAACCTGAAGACACTGCTGTCTATAGCTGCGCAGCAGCCTCTGATTACGGGCTGCCCCGCGAGGATTTCTTGTACGATTACTGGGGTCAGGGAACGCAGGTAACTGTTAGCTCAACATCACACCATCATCATCACCATTAATGAGGATCCCCGGC | - | SL1 | SR2 | CAGTCCAGTTACGCTGGAGTC | GGTCAGGTATGATTTAAATGGTCAGT |

| Supplemental Table 4: Primers | | |
|---|---|---|
| **Primer** | **Sequence (5'-3')** | **Purpose** |
| SL1 | CAGTCCAGTTACGCTGGAGTC | Sequence pSMART insert |
| SR2 | GGTCAGGTATGATTTAAATGGTCAGT | Sequence pSMART insert |
| JNH1 | CAAATTGAACTGGCGGTACTGC | Recombineering confirmation |
| JNH2 | GCCGTTAACTGATGCTCTGG | Recombineering confirmation |
| JNH3 | GCAGTCACGCTATTATTAAGCTGG | Recombineering confirmation |
| JNH4 | GTTTTCGCCACCCGAACTG | Recombineering confirmation |
| JNH5 | GAAGTGACCGGCGATCAAATG | Recombineering confirmation |
| JNH6 | GCCTGGGCAATGATCAATATGC | Recombineering confirmation |
| JNH5 | GCATTGCTTTTTACCGTATTGTCTAAC | Recombineering confirmation |
| JNH6 | GATTATTATGGTGTCACGCCATCTC | Recombineering confirmation |
| JNH7 | GTGCGTAATTGTGCTGATCTCTT | Sequence yibD pCOLA insert |
| JNH8 | AAGTTGGAACCTCTTACGTGC | Sequence pCOLA insert |
| JNH9 | TTGTGAGCGGATAACAATTCCC | Sequence pETM6 insert |
| JNH10 | ACCCCTCAAGACCCGTTTAG | Sequence pETM6 insert |
| JNH11 | GGGAGACCACAACGG | Sequence pCASCADE gRNA insert |
| JNH12 | CGCAGTCGAACGACCG | Sequence pCASCADE gRNA insert |
| JNH13 | TGTATATCTCCTTCTTAAAGTTAAACAAAATTAT T | Amplify pETM6 backbone for cloning for T7 gene insert |
| JNH14 | CTGAGCAATAACTAGCATAACCCC | Amplify pETM6 backbone for cloning for T7 gene insert |
| JNH15 | aataattttgtttaactttaagaaggagatatacatATGGCTAG CAAAGGAGAAG | Amplify GFPuv with homology arms for pETM6 gibson assembly |
| JNH16 | ccaaggggttatgctagttattgctcagTTATTTGTAGAGCT CATCCATG | Amplify GFPuv with homology arms for pETM6 gibson assembly |
| JNH17 | CTGGAAAAAGGAGATATACCATGGTGAGCAA GGGCGAGG | Amplify Addgene Cat#134939 for roGFP2 insert |
| JNH18 | GTGAGTCGTATTAGAAGAGCtcattaCTTGTACA GCTCGTCCATGCC | Amplify Addgene Cat#134939 for roGFP2 insert |
| JNH19 | CTGAGCAATAACTAGCATAACCCC | Amplify pETM6 for roGFP2 insert |
| JNH20 | AATAATTTTGTTTAACTTTAAGAAGGAGATATAC AATGGTGAGCAAGGGCGAGG | Amplify roGFP for pETM6 gibson asssembly |

| | | |
|---|---|---|
| JNH21 | CCAAGGGGTTATGCTAGTTATTGCTCAGTTAG TGATGGTGATGGTGATGAGATCTCTT | Amplify roGFP for pETM6 gibson asssembly |
| JNH22 | cagacgtaaaaaaagTCGAGTTCCCCGCGCCAGC GGGGATAAACCGAAAAAAAAACCCC | Amplify pCASCADE for trxB gRNA insert |
| JNH23 | taaattccctacaatCGGTTTATCCCCGCTGGCGCG GGGAACTCGAGGTGGTACCAGATC | Amplify pCASCADE for trxB gRNA insert |
| JNH24 | GGATCCGAGTCACACTGG | Amplify pCOLA backbone for chaperone insertion |
| JNH25 | AGATTATCCTCCTACACAGAAGTTATC | Amplify pCOLA backbone for Evr1p insertion |
| JNH26 | AACTTCTGTGTAGGAGGATAATCTATGAAGGC AATTGATAAGATGAC | Amplify Evr1p with homology arms for gibson assembly into pCOLA |
| JNH27 | CCAGTGTGACTCGGATCCTCATTACTCGTCCC ACCC | Amplify Evr1p with homology arms for gibson assembly into pCOLA |
| JNH28 | tatatcggcatagtataatacgacGTGTAGGAGGATAAT CTATGAAG | Amplify pCOLA to insert the EM7 promoter |
| JNH29 | ctatgccgatgattaattgtcaacACGCAATTAATGTAAG TTAGC | Amplify pCOLA to insert the EM7 promoter |
| JNH30 | CTCAAGGGCATCGGTCGAGA | Sequence EM7 pCOLA insert |
| JNH31 | TCATTACTCGTCCCACCC | Amplify pCOLA Erv1p for DsbC or PDI insertion |
| JNH32 | GGGTGGGACGAGTAATGATCTAGAGATTAAAG AGGAGAATACT | Amplify DsbC for gibson assembly into EM7-Erv1p-DsbC-pCOLA |
| JNH33 | ccagtgtgactcGGATCCTCATTATTTACCGCTGGT CA | Amplify DsbC for gibson assembly into EM7-Erv1p-DsbC-pCOLA |
| JNH34 | ACGCAATTAATGTAAGTTAGCTCAC | Cloning pCOLA empty vector |
| JNH35 | TAATGAGGATCCCCGGCTTATCG | Amplify pSMART backbone for yibDp VHH insert |
| JNH36 | CATAGATTATCCTCCTACACAGAAGTTATCCTG ACGTTTTA | Amplify pSMART backbone for yibDp VHH insert |
| JNH37 | aataattttgtttaactttaagaaggagatatacatATGCAGGT GCAGTTACAGG | Amplify VHH72 with homology arms for pETM6 assembly |
| JNH38 | ccaaggggttatgctagttattgctcagTCATTAGTGGTGA TGATGATGG | Amplify VHH72 with homology arms for pETM6 assembly |
| JNH39 | aataattttgtttaactttaagaaggagatatacatATGGCCGA TGTCCAGCT | Amplify VHH GFP minimizer with homology arms for pETM6 assembly |
| JNH40 | ccaaggggttatgctagttattgctcagTCATTAGTGGTGG | Amplify VHH GFP minimizer with homology |

| | | |
|---|---|---|
| | TGATGATGAT | arms for pETM6 assembly |
| JNH41 | aataattttgtttaactttaagaaggagatatacatATGGAAGT ACAACTGCAGGAAT | Amplify VHH C9 BoNT/A with homology arms for pETM6 assembly |
| JNH42 | ccaaggggttatgctagttattgctcagTCATTAATGATGAT GGTGATGAT | Amplify VHH C9 BoNT/A with homology arms for pETM6 assembly |

**Supplemental Table 5: Nanobodies**

| VHH name | Amino acid sequence | MW (kDa) | Cys | Disulfs | Immunogen | Source |
|---|---|---|---|---|---|---|
| VHH72 | MQVQLQESGGGLVQAGGSLRLSCAASGRTFSEYAMGWFRQAPGKEREFVATISWSGGSTYYTDSVKGRFTISRDNAKNTVYLQMNSLKPDDTAVYYCAAAGLGTVVSEWDYDYDYWGQGTQVTVSSGSHHHHHH | 14.7 | 2 | 1 | SARS-CoV-2 | (Huo, J. et al. 2020) |
| J3 | MEVQLVESGGGLVQAGGFLRLSCELRGSIFNQYAMAWFRQAPGKEREFVAGMGAVPHYGEFVKGRFTISRDNAKSTVYLQMSSLKPEDTAIYFCARSKSTYISYNSNGYDYWGRGTQVTVSSHHHHHH | 14.4 | 2 | 1 | HIV | (Weiss, R. A. & Verrips, C. T. 2019) |
| H6 antivenin | MQVQLQESGGGLVQAGGSLTLACTAS GRTFDRYA VGWFRQTPGKDREFVAT ISWSGGTT RYADSVKGRFTVS RDNAKN TVYL QMNT LKPEDTAVYYC AADLALSTVDEAVARYWGQGTQVTVSSHHHHH | 14.2 | 2 | 1 | B. atrox venom | (Bailon Calderon, H. et al. 2020) |
| Re9F06 | M GSQVQLVESGGGLVQAGGSLRLSCAASGRTFSNDALGWFRQAPRKEREFVAAINWNSGTYYADSVKGRFTISRDNAKNTVYLQMNSLKPEDTAVYSCAAASDYGLPREDFLYDYWGQGTQVTVSSTS HHHHHH | 14.8 | 2 | 1 | SARS-CoV-2 | (Güttler, T. et al. 2021) |
| Re5D06 | M GSQVQLVESGGGLVQPGGSLRLSCAASGITLDYYAIGWFRQAPGKEREGVSRIRSSDGSTNYADSVKGRFTMSRDNAKNTVYLQMNSLKPEDTAVYYCAYGPLTKYGSSWYWPYEYDYWGQGTQVTVSSTS HHHHHH | 15.4 | 2 | 1 | SARS-CoV-2 | (Güttler, T. et al. 2021) |
| VCAM1 | MQVQLQESGGGSVQTGGSLRLSCAASGY | 14.8 | 2 | 1 | Human | ( Ta, D. T. |

| | | | | | | |
|---|---|---|---|---|---|---|
| | TNSIMYMAWFRQAPGKKREG VAAIRFPDDSAYYAGSVKGRFTISHDNAKNT VYLQMNNLNPEDTAMYYC AARSSPYSFAWNDPSNYNYWGQGTQVTV SSHHHHHH | | | | VCAM1 | 2015) (Broisat, A. et al. 2012) |
| GFP enhancer | MAQVQLVESGGALVQPGGSLRLSCAASGF PVNRYSMRWYRQAPGKEREWVAGMSSAG DRSSYEDSVKGRFTISRDDARNTVYLQMN SLKPEDTAVYYCNVNVGFEYWGQGTQVTV SSHHHHHH | 13.7 | 2 | 1 | GFP | (Kirchhofer, A. et al. 2010) |
| GFP minimizer | MADVQLQESGGGSVQAGGSLRLSCAASG DTFSSYSMAWFRQAPGKECELVSNILRDGT TTYAGSVKGRFTISRDDAKNTVYLQMVNLK SEDTARYYCAADSGTQLGYVGAVGLSCLD YVMDYWGKGTQVTVSSHHHHHH | 15.0 | 4 | 2 | GFP | (Kirchhofer, A. et al. 2010) |
| E7 anti BoNT/A | M DVQLQASGGGSAQAGGSLTLSCAASGLIFS NYIMGWFRQAPGKDREFVAAISRQGTTPYY VNSGNDRFTISRDNAKNTGTLQMNHLKPE DTAVYYCAVDSFLSNIRVG-KADYWGQGTQ VT HHHHHH | 13.9 | 2 | 1 | BoNT/A | (Goldman, E. R. et al. 2008) |
| C9 anti BoNT/A | M EVQLQESGGGVVQPGGSLKLSCSGSGAIF DTYDVGWYRQAPGKRRELVASITATARGRI DYNYFAQGRFTISKDNAANTVYLQMDHLEP GDTAVYYCKTSVTGWGRGTQVTVSSAMQ GTLPT HHHHHH | 14.0 | 2 | 1 | BoNT/A | (Goldman, E. R. et al. 2008) |
| m36.4 | M QVQLVQSGG GLVQPGGSLRLSCAAS AFDF SDYE MSWVREAPGKGLEWIGE INDS GNT IYNPSLKSRVTISRDNSKNTLYLQMNTLRAE DTAIYYC AIYGGNSGGEY WGQGTLVTVSS HHHHHH | 13.6 | 2 | 1 | HIV | (Chen, W. et al. 2014) |
| 1B5 | M EVQLVESGGGLVQPGGSLRLSCAASGRAT FD DYAIG WFRQAPGKEREGVS YIGCNDGATYYAGSVKG RFTISCDYAKNTVYLQMNSLKPEDTAVYYC AA AAQWATIRWIHEYDYNI WGQGTQVTVSS HHHHHH | 14.8 | 4 | 2 | HIV | (Strokappe, N. et al. 2012) |
| 3E3 | M | 13.6 | 2 | 1 | HIV | (McCoy, L. |

| | | | | | | |
|---|---|---|---|---|---|---|
| | EVQLVESGGGLVQPGGSLRLSCAASQFTLE SYAIGWFRQAPGKDSEGVACISSSTYY ADSVKGRFTISRDNAKNTVYLQMESLKPED TAVYHCATSGAGSYCTLRAFGSWGQGTQV TVSS HHHHHH | | | | | E. et al. 2014) |
| 2E7 | M EVQLVESGGGLVQPGGSLRLSCAASGNIVS IDAAG WFRQAPGKQREPVA TILTGGATNYADSVKGRFTISRDNAKNTVYL QMNSLKPEDTAVYYCYA PMIYYGGRYSDY WGQGTQVTVSS HHHHHH | 13.8 | 2 | 1 | HIV | (Strokappe, N. et al. 2012) |
| JM3 | M EVQLVESGGGLVQPGGSLRLSCAAS GFTLEDYS IGWFRQAPGKEREGVS CISDSDGRT YYADSVKGRFTISRDNAKNTVYLQMNSLKP EDTAVYYC ATDCTVVPSLLYAMDS GKGTQVTVSS HHHHHH | 14 | 4 | 2 | HIV | (Matz, J. et al. 2013) |

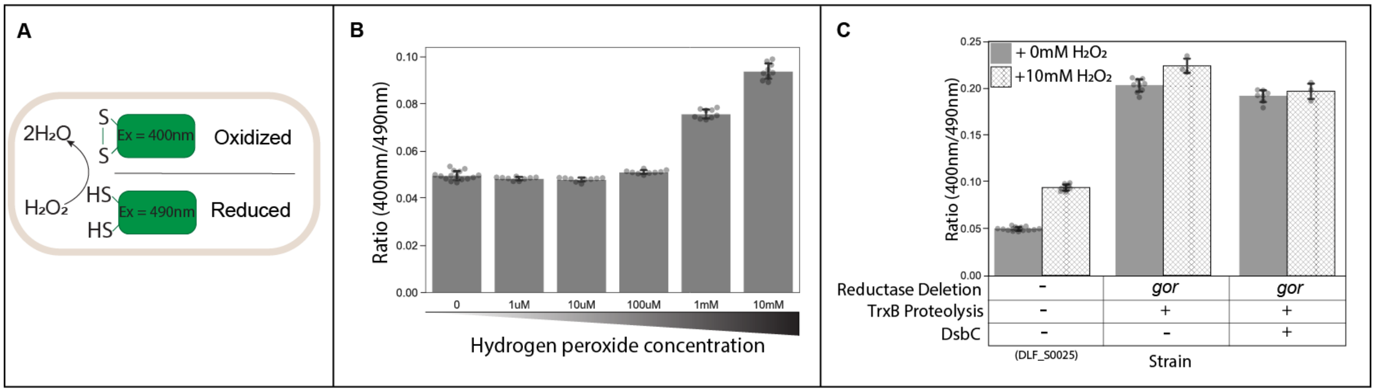

**Figure S1: Peroxide addition for validation of the roGFP redox sensor.**

A) Overview of the redox-sensitive GFP (roGFP) assay: Assessing cytoplasmic redox state by measuring the excitation spectrum and emission ratio of roGFP (400nm/490nm). Increasing ratio indicates higher roGFP oxidation. B) Hydrogen peroxide (0-10mM) addition to the *E. coli* control strain (DLF_S0025) expressing roGFP under the yibD promoter after 37°C microfermentations. Raw data points indicate the number of replicates. C) Comparison of roGFP ratio before (solid bar) and after (patterned bar) 10mM hydrogen peroxide addition. Strains include control (DLF_S0025), reductase control (JNH_Redox5), and reductase control with dynamic overexpression of DsbC (JNH_Redox6).

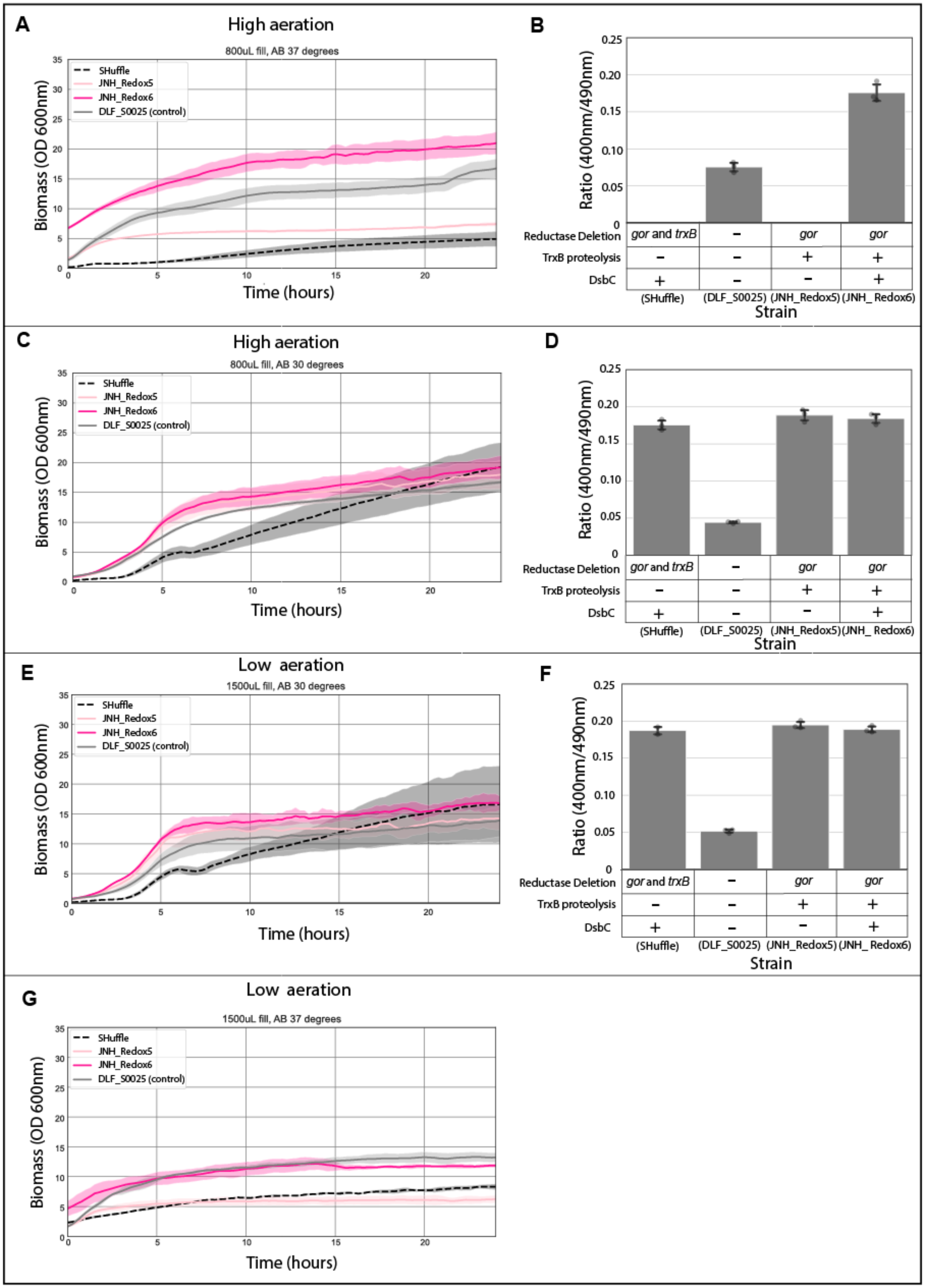
A
High aeration
800uL fill, AB 37 degrees
SHuffle
JNH_Redox5
JNH_Redox6
DLF_S0025 (control)
Biomass (OD 600nm)
Time (hours)
B
Ratio (400nm/490nm)
Reductase Deletion
gor and trxB
gor
TrxB proteolysis
DsbC
(SHuffle)
(DLF_S0025)
(JNH_Redox5)
(JNH_ Redox6)
Strain
C
High aeration
800uL fill, AB 30 degrees
D
E
Low aeration
1500uL fill, AB 30 degrees
F
G
Low aeration
1500uL fill, AB 37 degrees

**Figure S2: Biolector studies with SHuffle and dynamic control strains.**

Dynamic control enhances strain robustness in SHuffle *E. coli* based on Biolector data. A) Biolector growth curves of SHuffle and dynamic control strains, JNH_Redox5 and JNH_Redox6, expressing yibD-roGFP-pSMART in AB autoinduction media at 37°C under high aeration (800µL fill volume). Color indicates strain as labeled in the legend. B) Redox emission ratios measured at the end of the Biolector run from panel A, with corresponding strain modifications marked below each bar. C) Biolector growth curves of the strains at 30°C under low aeration (1500µL fill volume). D) Redox emission ratios measured at the end of the Biolector run from panel C. E) Biolector growth curves of the strains at 30°C under low aeration (1500µL fill volume). F) Endpoint redox emission ratios measured at the end of the Biolector experiment in panel E. G) Biolector growth curves of the strains at 37°C under low aeration (1500µL fill volume). The roGFP reporter did not induce under these conditions.

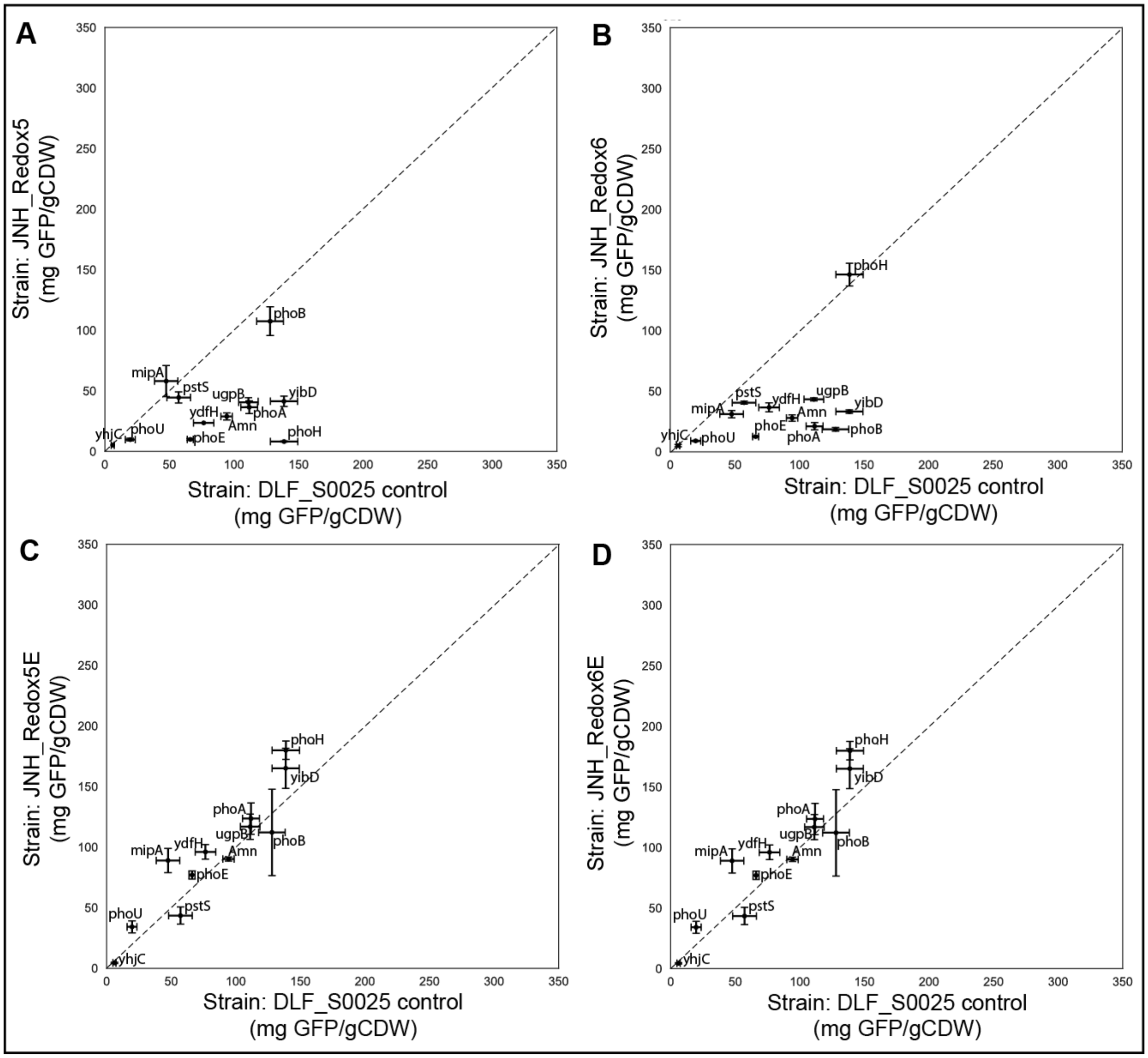

**Figure S3: Erv1p impact on GFPuv expression robustness in DC Redox strains**

Screening a phoB promoter library in 5 strains at 37°C in AB media. Data points represent GFPuv expression controlled by phoB promoters, as labeled. Below the dashed line indicates improved expression in the control strain (DLF_S0025), above the dotted line indicates improved expression in the engineered strain. Data on the dashed line indicates robust expression. A) Control strain

(DLF_S0025) x-axis, and JNH_Redox5 y-axis. B) Control strain (DLF_S0025) x-axis, and JNH_Redox6 y-axis. C) Control strain (DLF_S0025) x-axis, and JNH_Redox5E y-axis. D) Control strain (DLF_S0025) x-axis, and JNH_Redox6E y-axis.

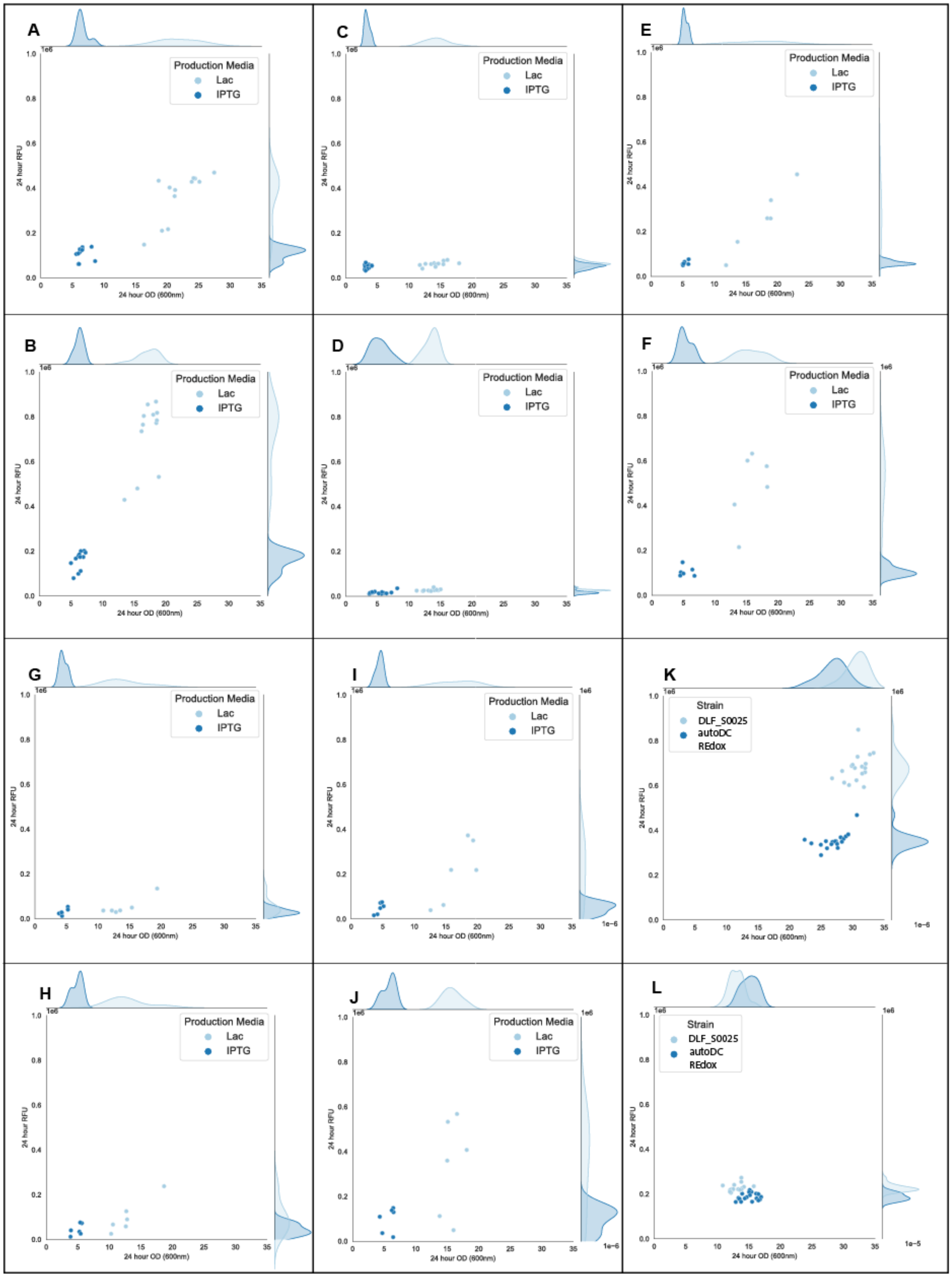
A
B
C
D
E
F
G
H
I
J
K
L
Production Media
Lac
IPTG
Strain
DLF_S0025
autoDC REdox
24 hour RFU
24 hour OD (600nm)
1e6
1e-6
1e-5
0.0
0.2
0.4
0.6
0.8
1.0
0
5
10
15
20
25
30
35

### Figure S4: Strain robustness studies and protein expression studies with GFPuv

Growth ($OD_{600\ nm}$) and GFPuv expression robustness were assessed under various conditions: A) BL21 (DE3), 37°C. B) BL21 (DE3), 30°C. C) SHuffle, 37°C. D) SHuffle, 30°C. E) BL21 (DE3) with EM7-Erv1p, 37°C. F) BL21 (DE3) with EM7-Erv1p, 30°C. G) BL21(DE3) + EM7-Erv1p-DsbC, 37°C. H) BL21(DE3) + EM7-Erv1p-DsbC, 30°C. I) BL21(DE3) + EM7-Erv1p-PDI, 37°C. J) BL21(DE3) + EM7-Erv1p-PDI, 30°C. K) Strains DLF_S0025 and autoDC REdox, 37°C. L) Strains DLF_S0025 and autoDC REdox, 30°C. In panels A-J, color indicates the T7-GFPuv induction method: studier's lactose autoinduction media (Lac, light blue) or IPTG induction (dark blue). In panels K-L, color indicates the strain. AB autoinduction media was used for yibD-GFPuv phosphate induction.

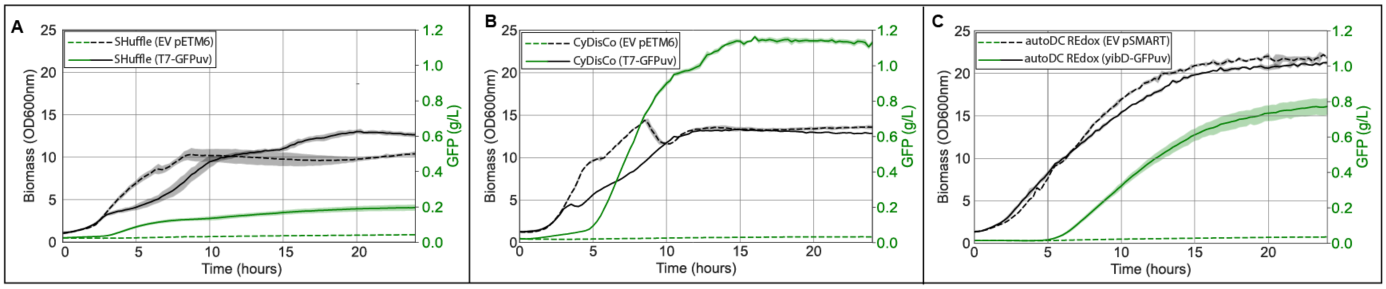

### Figure S5: BioLector growth robustness studies

Biolector studies of SHuffle (A), CyDisCo (B), and autoDC Redox (C) at 37 degrees with 800µL fill volume. Growth curves shown in black, GFPuv expression in green. Dashed line indicates strain with empty vector plasmid, solid line indicates GFPuv plasmid. Shading represents standard deviation (n=3). SHuffle and CyDisCo studies performed in Studier's lactose autoinduction media, while autoDC REdox used AB autoinduction media.

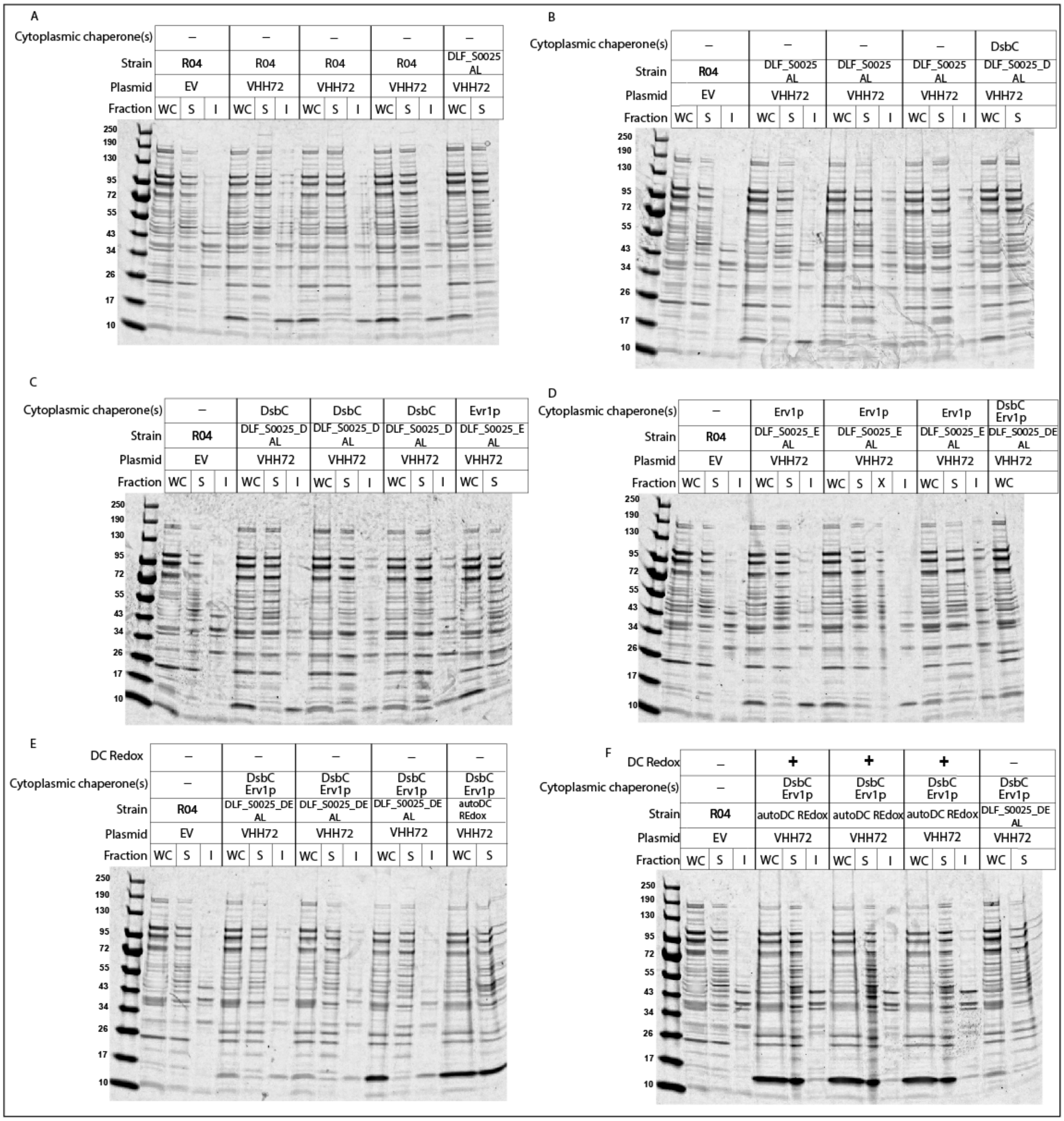

### Figure S6: SDS-PAGE gels from dynamic control chaperone studies

SDS-PAGE gels utilized for the analysis of VHH72 expression levels (14.7 kDa) in different fractions: whole cell (WC), soluble (S), and insoluble (I). The quantification of these fractions can be found in Figure 3H. Each gel represents triplicate VHH72 expression in a specific strain, with strain modifications indicated above each gel. A) VHH72 expression in control strain R04. B) VHH72 expression in control strain DLF_S0025 AL, which incorporates DC "off" valves. C) VHH72 expression in DLF_S0025_D AL (with DsbC dynamic overexpression). D) VHH72 expression in DLF_S0025_E AL (with Erv1p dynamic overexpression). E) VHH72 expression in DLF_S0025_DE AL (with both DsbC and Erv1p dynamic overexpression). F) VHH72 expression in auto DC REdox with combined DC redox state, DsbC, and Erv1p dynamic overexpression.

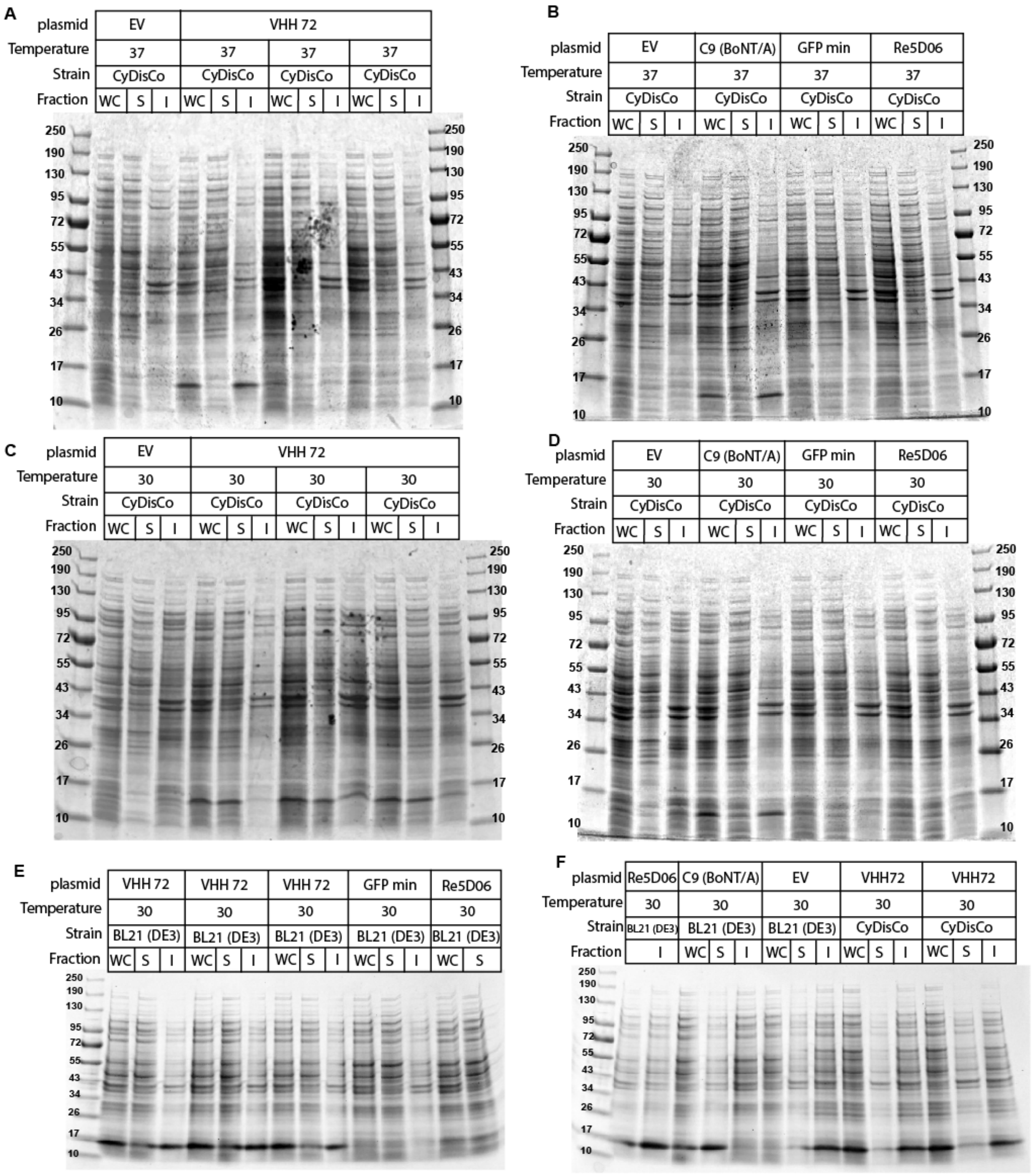
A
plasmid EV VHH 72
Temperature 37 37 37 37
Strain CyDisCo CyDisCo CyDisCo CyDisCo
Fraction WC S I WC S I WC S I WC S I
250 190 130 95 72 55 43 34 26 17 10
B
plasmid EV C9 (BoNT/A) GFP min Re5D06
Temperature 37 37 37 37
Strain CyDisCo CyDisCo CyDisCo CyDisCo
Fraction WC S I WC S I WC S I WC S I
250 190 130 95 72 55 43 34 26 17 10
C
plasmid EV VHH 72
Temperature 30 30 30 30
Strain CyDisCo CyDisCo CyDisCo CyDisCo
Fraction WC S I WC S I WC S I WC S I
250 190 130 95 72 55 43 34 26 17 10
D
plasmid EV C9 (BoNT/A) GFP min Re5D06
Temperature 30 30 30 30
Strain CyDisCo CyDisCo CyDisCo CyDisCo
Fraction WC S I WC S I WC S I WC S I
250 190 130 95 72 55 43 34 26 17 10
E
plasmid VHH 72 VHH 72 VHH 72 GFP min Re5D06
Temperature 30 30 30 30 30
Strain BL21 (DE3) BL21 (DE3) BL21 (DE3) BL21 (DE3) BL21 (DE3)
Fraction WC S I WC S I WC S I WC S I WC S
250 190 130 95 72 55 43 34 26 17 10
F
plasmid Re5D06 C9 (BoNT/A) EV VHH72 VHH72
Temperature 30 30 30 30 30
Strain BL21 (DE3) BL21 (DE3) BL21 (DE3) CyDisCo CyDisCo
Fraction I WC S I WC S I WC S I WC S I
250 190 130 95 72 55 43 34 26 17 10

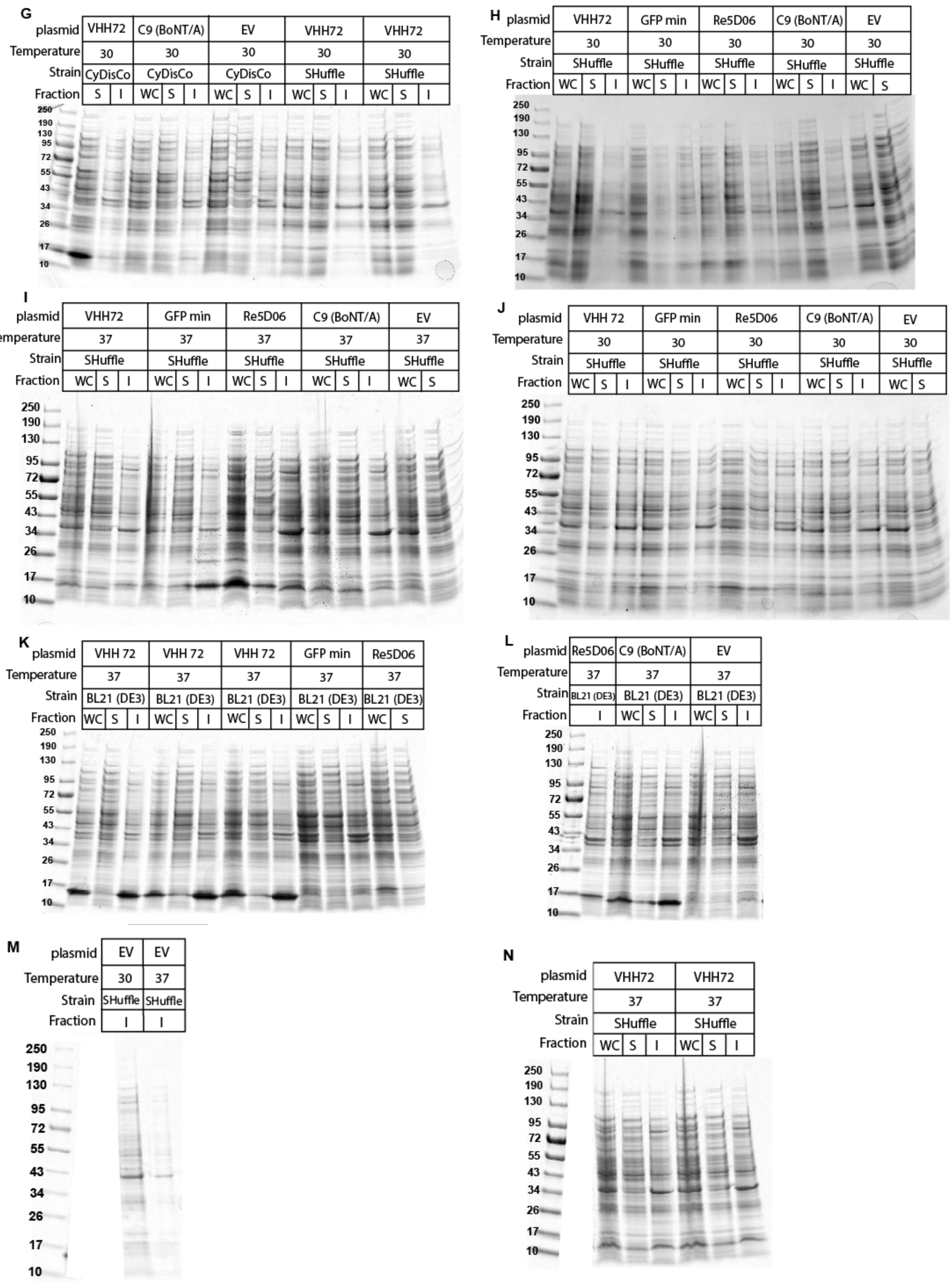
G
plasmid VHH72 C9 (BoNT/A) EV VHH72 VHH72
Temperature 30 30 30 30 30
Strain CyDisCo CyDisCo CyDisCo SHuffle SHuffle
Fraction S I WC S I WC S I WC S I WC S I
250 190 130 95 72 55 43 34 26 17 10
H
plasmid VHH72 GFP min Re5D06 C9 (BoNT/A) EV
Temperature 30 30 30 30 30
Strain SHuffle SHuffle SHuffle SHuffle SHuffle
Fraction WC S I WC S I WC S I WC S I WC S
250 190 130 95 72 55 43 34 26 17 10
I
plasmid VHH72 GFP min Re5D06 C9 (BoNT/A) EV
Temperature 37 37 37 37 37
Strain SHuffle SHuffle SHuffle SHuffle SHuffle
Fraction WC S I WC S I WC S I WC S I WC S
250 190 130 95 72 55 43 34 26 17 10
J
plasmid VHH 72 GFP min Re5D06 C9 (BoNT/A) EV
Temperature 30 30 30 30 30
Strain SHuffle SHuffle SHuffle SHuffle SHuffle
Fraction WC S I WC S I WC S I WC S I WC S
250 190 130 95 72 55 43 34 26 17 10
K
plasmid VHH 72 VHH 72 VHH 72 GFP min Re5D06
Temperature 37 37 37 37 37
Strain BL21 (DE3) BL21 (DE3) BL21 (DE3) BL21 (DE3) BL21 (DE3)
Fraction WC S I WC S I WC S I WC S I WC S
250 190 130 95 72 55 43 34 26 17 10
L
plasmid Re5D06 C9 (BoNT/A) EV
Temperature 37 37 37
Strain BL21 (DE3) BL21 (DE3) BL21 (DE3)
Fraction I WC S I WC S I
250 190 130 95 72 55 43 34 26 17 10
M
plasmid EV EV
Temperature 30 37
Strain SHuffle SHuffle
Fraction I I
250 190 130 95 72 55 43 34 26 17 10
N
plasmid VHH72 VHH72
Temperature 37 37
Strain SHuffle SHuffle
Fraction WC S I WC S I
250 190 130 95 72 55 43 34 26 17 10

### Figure S7: SDS-PAGE gels from CyDisCo VHH expression

SDS-PAGE gels that were used to analyze VHH expression levels in shake flask studies quantified in Figure 3I and Figure 3J. CyDisCo indicates BL21(DE3) + EM7-Erv1p-PDI and Studier's lactose autoinduction media was used for induction. Fractions are labeled as the whole cell (WC), soluble (S), and insoluble (I). The VHH expressed in each gel, along with the strain and expression temperature are indicated above the gel. A and C) VHH72, 14.7 kDa , B and D) VHH C9 BoNT/A, 14kDa, VHH GFP minimizer, 15 KDa, and VHH Re9F06, 14.8 kDa.

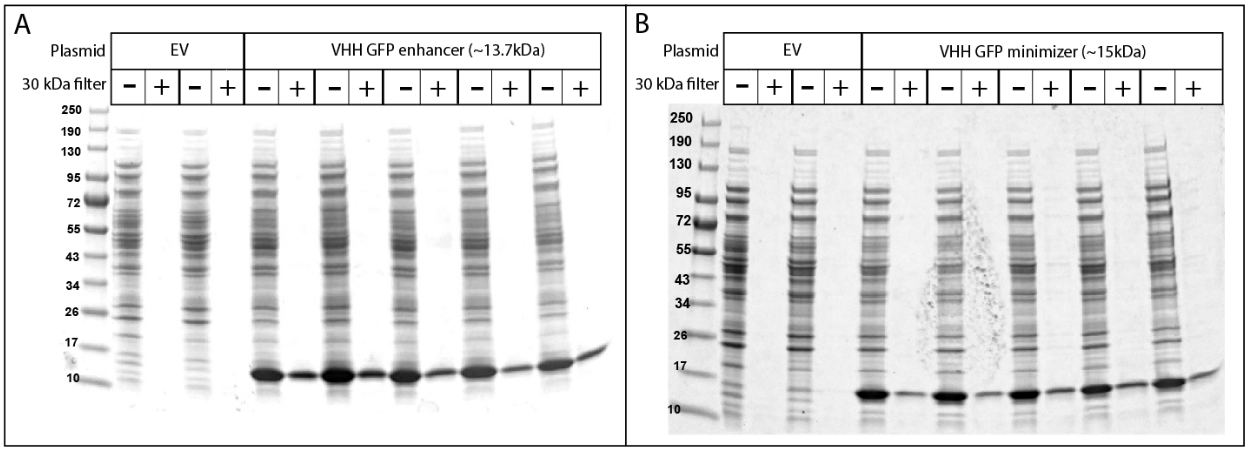

### Figure S8: VHH GFP enhancer and VHH GFP minimizer SDS-PAGE gels before and after filtration

SDS-PAGE analysis was performed on five biological replicates of each VHH, comparing the 30kDa filtrate (+) with the unfiltered soluble fraction (-). Panel A shows the SDS-PAGE gel for VHH GFP enhancer (~13.7 kDa), while panel B displays the SDS-PAGE gel for VHH GFP minimizer (~15 kDa).

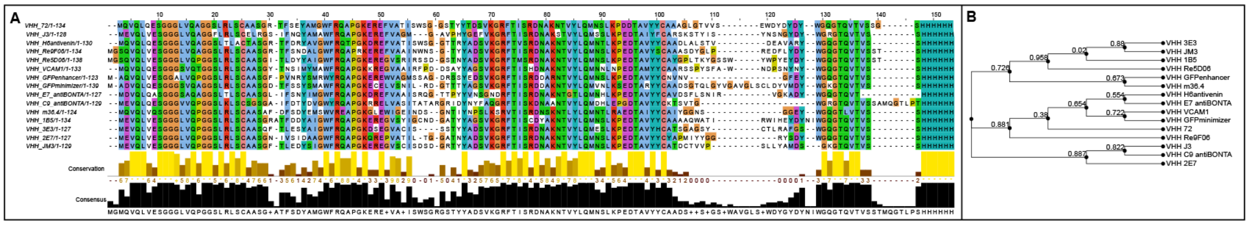

### Figure S9: VHH sequence alignment and phylogenetic tree

VHH sequence alignment (A) and phylogenetic tree (B) obtained using the bacterial and viral bioinformatics resource center multiple sequence alignment tool. The clustal color schemes highlight amino acid similarities: hydrophobic (blue), positive charge (red), negative charge (magenta), polar (green), cysteines (pink), glycines (orange), prolines (yellow), aromatic (cyan), while unconserved regions are displayed in white.

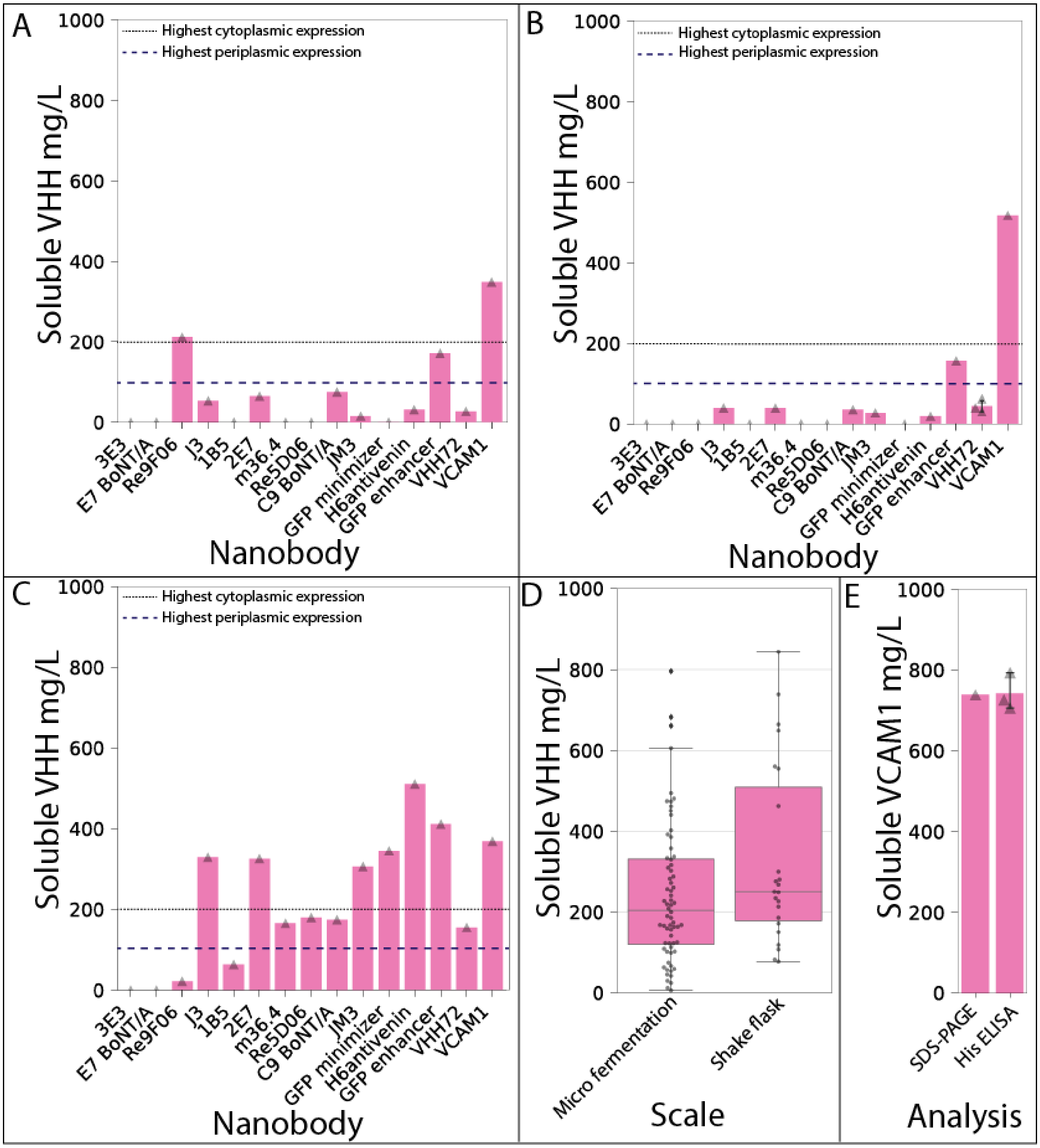

**Figure S10: Shake flask VHH expression in the control strain (R04) and autoDC REdox**

Soluble VHH titers were obtained through shake flask expression at both 30°C and 37°C. The gray dotted line represents the highest reported yield of soluble VHHs with cytoplasmic expression (~200 mg/L) found in the literature, while the blue dashed line indicates the upper limit of periplasmic VHH expression yield (<100 mg/L) (Zarschler et al. 2013; Hussack et al. 2011; Liu and Huang 2018). Panels A, B, and C illustrate the strain and temperature conditions for cytoplasmic expression of soluble VHHs: A) Control strain R04 at 30°C, B) Control strain R04 at 37°C, and C) autoDC REdox strain at 30°C. D) Box plots from all VHHs produced in micro fermentations (100μL) and shake flasks (20mL) (n= 12 nanobodies) additional data points represent technical replicates. E) Comparison of soluble VHH VCAM1 expression level, produced in a shake flask, obtained from each quantification method: SDS-PAGE and densitometry versus the His tag ELISA. VCAM1 was expressed in autoDC REdox at 37°C.

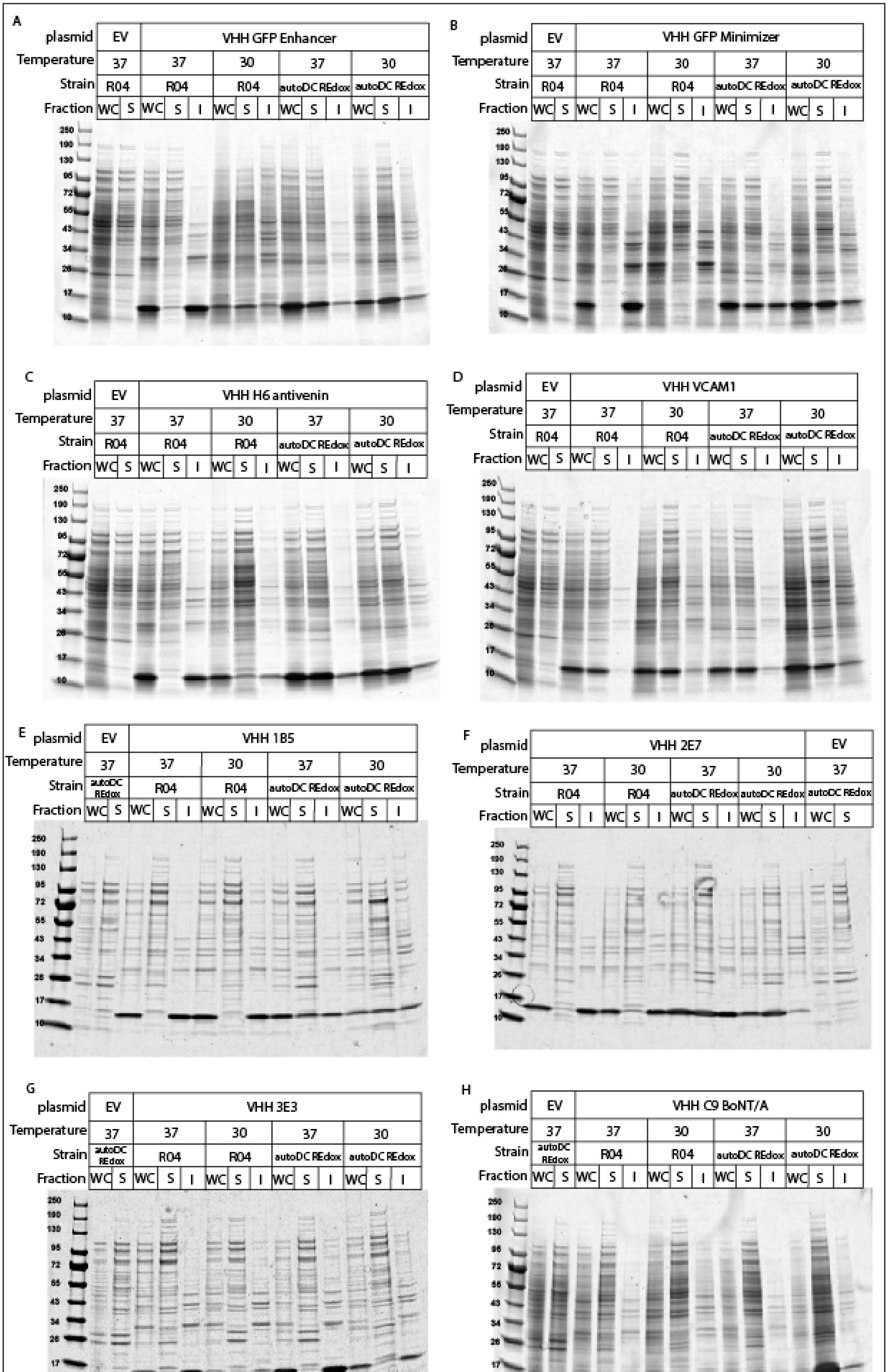
A
plasmid EV VHH GFP Enhancer
Temperature 37 37 30 37 30
Strain R04 R04 R04 autoDC REdox autoDC REdox
Fraction WC S WC S I WC S I WC S I WC S I
250 180 130 95 72 55 43 34 26 17 10
B
plasmid EV VHH GFP Minimizer
Temperature 37 37 30 37 30
Strain R04 R04 R04 autoDC REdox autoDC REdox
Fraction WC S WC S I WC S I WC S I WC S I
C
plasmid EV VHH H6 antivenin
Temperature 37 37 30 37 30
Strain R04 R04 R04 autoDC REdox autoDC REdox
Fraction WC S WC S I WC S I WC S I WC S I
D
plasmid EV VHH VCAM1
Temperature 37 37 30 37 30
Strain R04 R04 R04 autoDC REdox autoDC REdox
Fraction WC S WC S I WC S I WC S I WC S I
E
plasmid EV VHH 1B5
Temperature 37 37 30 37 30
Strain autoDC REdox R04 R04 autoDC REdox autoDC REdox
Fraction WC S WC S I WC S I WC S I WC S I
F
plasmid VHH 2E7 EV
Temperature 37 30 37 30 37
Strain R04 R04 autoDC REdox autoDC REdox autoDC REdox
Fraction WC S I WC S I WC S I WC S I WC S
G
plasmid EV VHH 3E3
Temperature 37 37 30 37 30
Strain autoDC REdox R04 R04 autoDC REdox autoDC REdox
Fraction WC S WC S I WC S I WC S I WC S I
H
plasmid EV VHH C9 BoNT/A
Temperature 37 37 30 37 30
Strain autoDC REdox R04 R04 autoDC REdox autoDC REdox
Fraction WC S WC S I WC S I WC S I WC S I

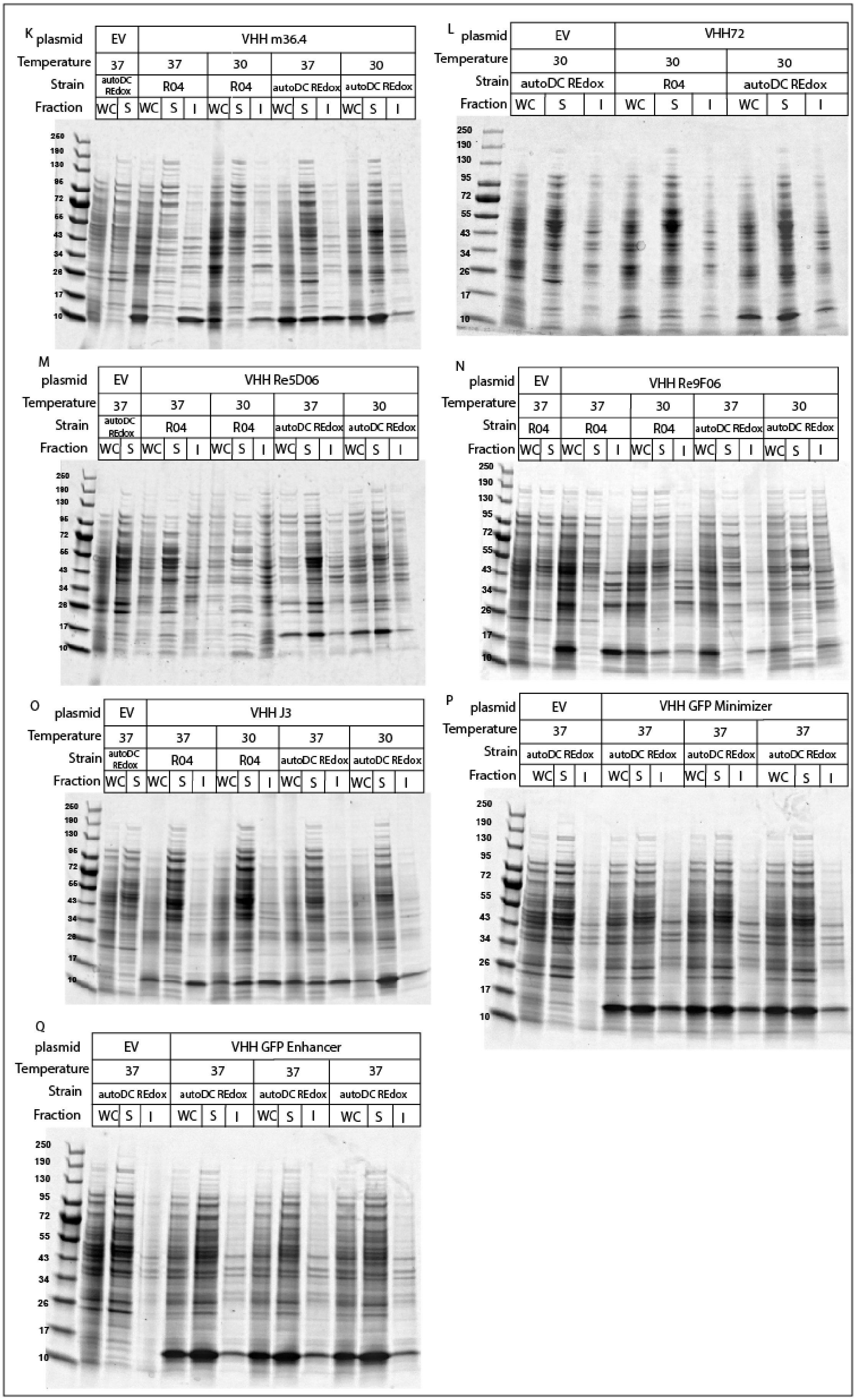
K
plasmid EV VHH m36.4
Temperature 37 37 30 37 30
Strain autoDC REdox R04 R04 autoDC REdox autoDC REdox
Fraction WC S WC S I WC S I WC S I WC S I
250 190 130 95 72 55 43 34 26 17 10
L
plasmid EV VHH72
Temperature 30 30 30
Strain autoDC REdox R04 autoDC REdox
Fraction WC S I WC S I WC S I
250 190 130 95 72 55 43 34 26 17 10
M
plasmid EV VHH Re5D06
Temperature 37 37 30 37 30
Strain autoDC REdox R04 R04 autoDC REdox autoDC REdox
Fraction WC S WC S I WC S I WC S I WC S I
250 190 130 95 72 55 43 34 26 17 10
N
plasmid EV VHH Re9F06
Temperature 37 37 30 37 30
Strain R04 R04 R04 autoDC REdox autoDC REdox
Fraction WC S WC S I WC S I WC S I WC S I
250 190 130 95 72 55 43 34 26 17 10
O
plasmid EV VHH J3
Temperature 37 37 30 37 30
Strain autoDC REdox R04 R04 autoDC REdox autoDC REdox
Fraction WC S WC S I WC S I WC S I WC S I
250 190 130 95 72 55 43 34 26 17 10
P
plasmid EV VHH GFP Minimizer
Temperature 37 37 37 37
Strain autoDC REdox autoDC REdox autoDC REdox autoDC REdox
Fraction WC S I WC S I WC S I WC S I
250 190 130 95 72 55 43 34 26 17 10
Q
plasmid EV VHH GFP Enhancer
Temperature 37 37 37 37
Strain autoDC REdox autoDC REdox autoDC REdox autoDC REdox
Fraction WC S I WC S I WC S I WC S I
250 190 130 95 72 55 43 34 26 17 10

**Figure S11: VHH expression SDS-PAGE gels.**

SDS-PAGE gels used to analyze VHH expression levels in the whole cell (WC), soluble (S), and insoluble (I) fractions quantified in Figure 4 and Figure S9. The VHH expressed in each gel, along with the strain (R04 or autoDC REdox) and expression temperature are indicated above the gel. A) VHH GFP-enhancer, 13.7kDa B) VHH GFP minimizer, 15 KDa C) VHH H6 antivenin, 14.2 kDa D) VHH VCAM1, 14.8 kDa E) VHH 1B5, 14.8kDa F) VHH 2E7, 13.8 kDa G) VHH 3E3, 13.6 kDa H) VHH C9 BoNT/A, 14kDa I) VHH E7 BoNT/A, 13.9 kDa J) VHH JM3, 14 kDa K) VHH m36.4, 13.6 kDa L) VHH72, 14.7 kDa M) VHH Re5D06, 15.4 kDa N) VHH Re9F06, 14.8 kDa. O) VHH J3, 14.4 kDa. P) VHH GFP minimizer, 15 KDa (biological replicates n=3). Q) VHH GFP-enhancer (biological replicates n=3), 13.7kDa. R) VHH GFP minimizer (biological replicates, n=3), 15 kDa.

Zarschler, Kristof, Stefanie Witecy, Franz Kapplusch, Christian Foerster, and Holger Stephan. 2013. “High-Yield Production of Functional Soluble Single-Domain Antibodies in the Cytoplasm of Escherichia Coli.” *Microbial Cell Factories* 12 (October): 97.
